# Multiview super-resolution microscopy

**DOI:** 10.1101/2021.05.21.445200

**Authors:** Yicong Wu, Xiaofei Han, Yijun Su, Melissa Glidewell, Jonathan S. Daniels, Jiamin Liu, Titas Sengupta, Ivan Rey-Suarez, Robert Fischer, Akshay Patel, Christian Combs, Junhui Sun, Xufeng Wu, Ryan Christensen, Corey Smith, Lingyu Bao, Yilun Sun, Leighton H. Duncan, Jiji Chen, Yves Pommier, Yun-Bo Shi, Elizabeth Murphy, Sougata Roy, Arpita Upadhyaya, Daniel Colón-Ramos, Patrick La Riviere, Hari Shroff

## Abstract

We enhance the performance of confocal microscopy over imaging scales spanning tens of nanometers to millimeters in space and milliseconds to hours in time, improving volumetric resolution more than 10-fold while simultaneously reducing phototoxicity. We achieve these gains via an integrated, four-pronged approach: 1) developing compact line-scanners that enable sensitive, rapid, diffraction-limited imaging over large areas; 2) combining line-scanning with multiview imaging, developing reconstruction algorithms that improve resolution isotropy and recover signal otherwise lost to scattering; 3) adapting techniques from structured illumination microscopy, achieving super-resolution imaging in densely labeled, thick samples; 4) synergizing deep learning with these advances, further improving imaging speed, resolution and duration. We demonstrate these capabilities on more than twenty distinct fixed and live samples, including protein distributions in single cells; nuclei and developing neurons in *Caenorhabditis elegans* embryos, larvae, and adults; myoblasts in *Drosophila* wing imaginal disks; and mouse renal, esophageal, cardiac, and brain tissues.

## Introduction

Confocal microscopy^1^ (CM) remains the dominant workhorse in biomedical optical microscopy due to its reliability and flexibility in imaging a wide variety of three-dimensional samples. Strengths of CM include diffraction-limited imaging in transparent samples; high-contrast imaging due to optical sectioning in densely labeled, thick specimens; and rapid selection of magnification and resolution tailored to the specimen of interest, often obtained simply by selecting the appropriate objective lens. Moreover, speed, signal-to-noise ratio (SNR), and optical sectioning^2^ may be further balanced^3^ by choosing an appropriate implementation, e.g., multifocal spinning disk CM for thin, living samples near the coverslip vs. single point-scanning CM for thicker, fixed samples.

Drawbacks of traditional CM include substantial point spread function (PSF) anisotropy (axial resolution 2-3 fold worse than lateral resolution, complicating the spatial analysis of subcellular structures); spatial resolution limited to the diffraction limit; depth-dependent degradation in scattering samples leading to signal loss at distances far from the coverslip; and three-dimensional illumination and volumetric bleaching, which may rapidly diminish the pool of available fluorescent molecules and lead to unwanted phototoxicity^4^.

Possible solutions to these problems include (1) decoupling illumination from detection^5^, detecting fluorescence from multiple directions^6^ and combining the results^7^, resulting in a more nearly isotropic PSF; (2) using the diffraction-limited illumination structure in combination with digital^8,9^ or optical post-processing^10–12^ to improve resolution; (3) or using light-sheet fluorescence microscopy (LSFM) to achieve higher SNR imaging while imparting less dose to the specimen^13^. However, these solutions have their own drawbacks. Multiview imaging approaches^14,15^ are usually performed at moderate numerical aperture (NA), limiting resolution. Point-scanning super-resolution methods deliver more dose and are less sensitive than the confocal architectures upon which they are built, limiting SNR and imaging duration. Imaging performance in LSFM remains tied to the propagation length of the illumination beam, forcing a compromise between axial resolution and field-of-view, and attempts to attain the highest detection NA in LSFM compromise imaging field^16,17^ or sensitivity and spatial resolution^18,19^. These problems impede direct interrogation of cellular and subcellular structure and function, particularly in large, three-dimensional samples such as tissues.

Here we address these challenges in the context of CM, improving imaging resolution and duration in diverse specimens including single cells, living embryos, and scattering tissues. First, we develop a compact line-scanning illuminator that enables sensitive, rapid, and diffraction-limited confocal imaging over a 175 x 175 μm^2^ area, which can be readily incorporated into multiview imaging systems^14,20^. Second, we develop reconstruction algorithms that fuse three line-scanning confocal sample views, enabling ^~^2-fold improvement in axial resolution relative to conventional confocal microcopy and recovery of signal otherwise lost to scattering. Third, we use deep learning algorithms to lower the illumination dose imparted by confocal microscopy, enabling clearer imaging than LSFM in living, light-sensitive, and scattering samples. Fourth, we use sharp, line illumination introduced from three directions to further improve spatial resolution along these directions, enabling better than ten-fold volumetric resolution enhancement relative to traditional CM. Finally, we show that combining deep learning with traditional multiview fusion approaches can produce super-resolution data from single confocal images, providing a route to rapid, optically sectioned, super-resolution imaging with higher sensitivity and speed than otherwise possible. We demonstrate the power of these advances on more than twenty distinct fixed and live samples, spanning orders of magnitude in spatial and temporal scale.

## Results

### Multiview confocal microscopy

We constructed our confocal imaging platform with several goals. First, we wished to take advantage of state-of-the-art multiview imaging workflows^21^ in which multiple specimen views are rapidly captured, registered, and fused to yield reconstructions with improved resolution, lessening the anisotropic distortion and axial confusion that results when imaging three-dimensional samples with conventional single-view microscopy. Second, we desired good optical sectioning in tissue tens of microns in thickness, i.e., thicker than a single cell. Third, we aimed to achieve efficient fluorescence detection without the reflective losses introduced by descanning (and potentially rescanning) the fluorescent emissions, as is typical in CM.

To meet these goals, we developed a multiview confocal platform (**Fig. 1a, S1**) with three objectives (0.8 NA, 0.8 NA, 1.2 NA)^20^ to serially scan sharp line illumination along each axis, simultaneously collecting fluorescence in epi-mode from each objective. We chose line-illumination as line-scanning confocal microscopy is much faster than point-scanning confocal microscopy, yet provides better optical sectioning than spinning-disk confocal microscopy^10^, thereby providing a good compromise between imaging speed and optical sectioning^3^.

**Fig. 1.**
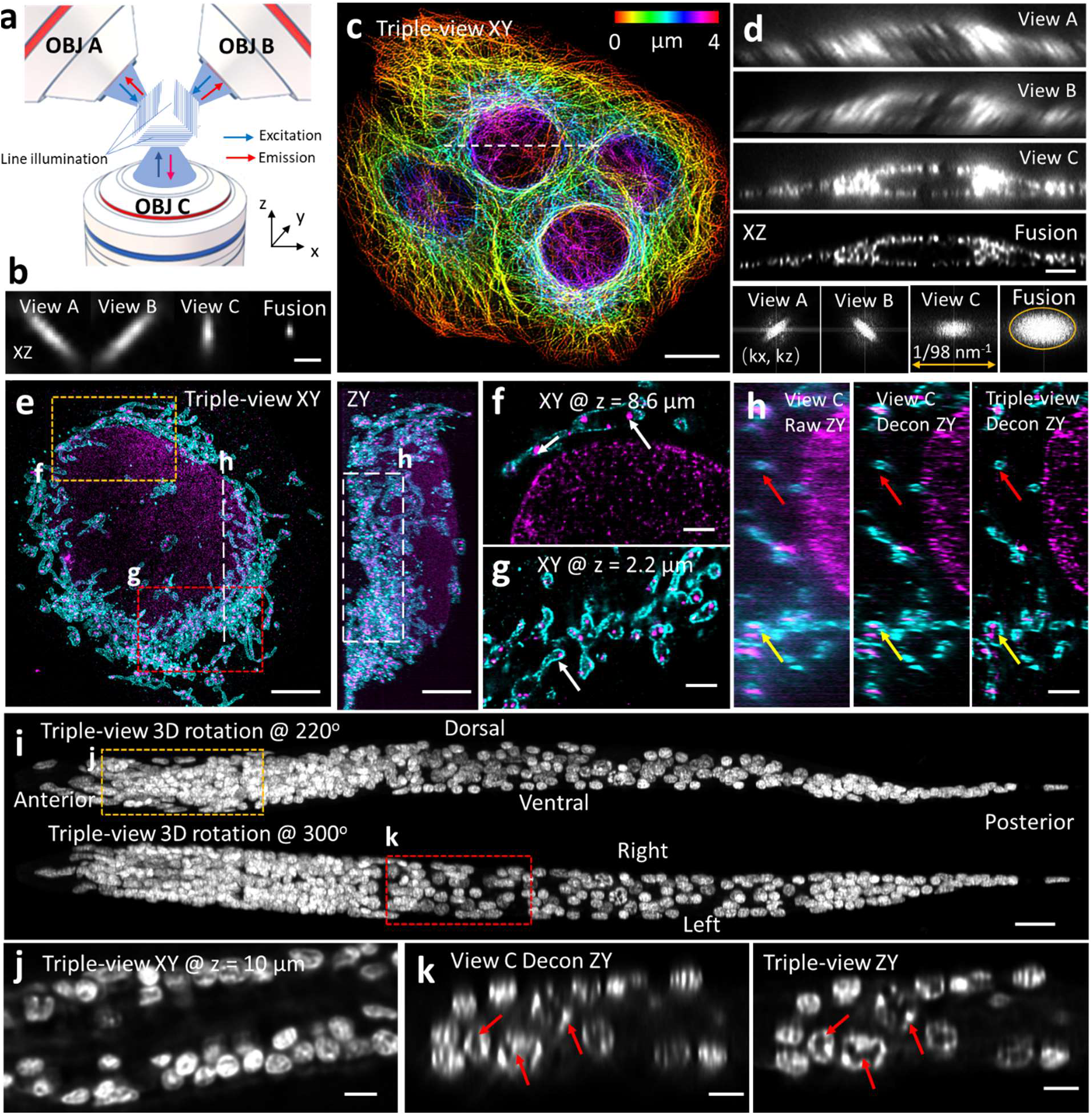
Multiview line confocal microscopy. **a)** Concept. Diffraction-limited line illumination (blue) is serially introduced into three objectives (OBJ A, B, C) and scanned through the sample, collecting fluorescence (red) from each view. See also **Fig. S1. b)** 100 nm beads, approximating PSF, as imaged in each view, and fused image after registering/deconvolving all three views. **c)** Lateral maximum intensity projection image of fixed U2OS cells immunolabeled with mouse-anti-alpha tubulin, anti-mouse-biotin, streptavidin Alexa Fluor 488, marking microtubules (fused result after registration and deconvolution of all three raw views). Height: color bar. **d)** Raw and fused axial views along the dashed line in **c)**. Fourier transforms of axial view are shown in last row, showing resolution enhancement after fusion. Orange oval: 1/260 nm^−1^ in half width and 1/405 nm^−1^ in half height. **e)** Lateral (left) and axial (right) maximum intensity projection images of 4x expanded U2OS cell immunolabeled against mitochondrial outer membrane (cyan, rabbit-anti-Tomm20, goat anti-rabbit Biotin, streptavidin Alexa Fluor 488) and double stranded DNA (magenta, mouse anti-dsDNA and donkey anti-Janelia Fluor 549). Triple-view result after registration/deconvolution is shown. **f, g)** Higher magnification views of yellow and red rectangular regions in **e)** at indicated axial distance from the beginning of the volume. White arrows: mitochondrial nucleoids encapsulated within Tomm20 signal. **h)** Axial view corresponding to dashed white line/rectangles in **e)** highlighting progressive increase in resolution from raw single view (left), to deconvolved single-view (middle), to deconvolved triple-view (right). Red, yellow arrows: mitochondria, nucleoids. See also **Movie S1. i)** Triple-view reconstruction of whole fixed L1 stage larval worm, nuclei labeled with NucSpot 488. Two maximum intensity projections are shown, taken after rotating volume 220 degrees (top) and 300 degrees (bottom) about x axis. **j)** Higher magnification lateral view 10 μm from the beginning of the volume, corresponding to the yellow dashed region in **i). k)** Higher magnification axial views (single planes) corresponding to the red rectangular region in **i)**, comparing single-view deconvolved result (left) to triple-view result (right). Red arrows highlight subnuclear structure better resolved in triple-view result. Scale bars: **b, f-h)** 1 μm, **c, i)** 10 μm, **d, e, j, k)** 3 μm.

To introduce and scan diffraction-limited line illumination, we developed compact, fiber-coupled, microelectromechanical systems (MEMS)-based line scanners that could be easily integrated into all three views (**Fig. S1b-d**). Careful characterization of these line scanners confirmed diffraction-limited performance over an imaging field at least 175 μm in each dimension (**Fig. S1e**). To provide confocality, we synchronized line illumination with the ‘rolling shutter’^22,23^ of our scientific complementary metal-oxide-semiconductor (sCMOS) detectors, enabling rapid, optically-sectioned imaging limited primarily by detector read out rate (25 Hz for 175 x 175 μm^2^, **Fig. S1d, e**). Since fluorescence descanning is unnecessary, only a dichroic, emission filter, tube lens and beam-folding mirrors were needed to relay the fluorescence from objective to detector, enabling light-efficient detection. Fluorescent volumes can be collected by using piezo-electric collars mounted to each objective to scan the focal plane over their range (100-150 μm), or over considerably larger extent by keeping the imaging plane stationary and scanning the sample stage (to 1 mm extent in this work). Further details, including descriptions of the software used to operate the microscope, are provided in **Methods**.

### Improved resolution in cellular and multicellular samples

We began our characterization of the multiview system by imaging 0.1 μm fluorescent beads. As expected, since such beads approximate the point spread function (PSF), they appeared highly anisotropic in each view: the full width at half maximum size of individual beads was 452 ± 42 nm laterally and 1560 ± 132 nm axially from the top two 0.8 NA views (*N* = 40) and 311 ± 17 nm laterally and 724 ± 88 nm axially from the bottom 1.2 NA view (*N* = 20). However, after performing registration and joint deconvolution^21,24,25^ of all three views by using illumination and detection PSFs derived from theory^26^ (Methods), the complementary information encoded in each view produced triple-view reconstructions with smaller, more nearly isotropic extent (**Fig. 1b**, lateral 235 ± 24 nm and axial 381 ± 23 nm, **Table S1**), better approximating the true bead size.

These gains extended to biological samples. First, we fixed U2OS cells and imaged immunolabeled microtubules using all three objectives, registering and fusing the volumes to produce the triple-view reconstruction (**Fig. 1c**). The triple-view result offered noticeably improved resolution, which we confirmed with decorrelation analysis^27^: 259 ± 22 nm laterally and 436 ± 31 nm axially (*N* = 10 slices) vs. 283 ± 11 nm and 772 ± 61 nm axially in the raw lower view (**Fig. 1d**). Second, we used our microscope to enhance nanoscale imaging of subcellular structures acquired by expansion microscopy (ExM^28^), a technique that enlarges fixed samples to enhance resolution. We expanded fixed U2OS cells immunolabeled to highlight DNA and Tomm20, imaged them from all three views, and reconstructed the triple-view result (**Fig. 1e, Movie S1**) to clearly resolve sub-organelle detail, including individual nucleoids residing within mitochondria (**Fig. 1f, g**). While the individual views also offered super-resolution detail due to the four-fold expansion procedure (e.g., resolution 112 ± 19 nm laterally, 152 ± 38 nm axially, *N* = 20 slices each, averaged over both colors, based on decorrelation analysis^27^), their anisotropic PSFs distorted the shapes of mitochondria and DNA puncta. Deconvolution of the single views offered some resolution improvement (64 ± 8 nm laterally, 100 ± 3 nm axially), but less than the triple-view result (54 ± 6 nm laterally, 78 ± 17 nm axially, **Fig. 1h, Movie S1**). In a third multicellular example, we fixed and imaged a whole *C. elegans* L1 larva marked with nuclear label NucSpot Live 488 (**Fig. 1i**), stitching and reconstructing the data (**Methods**) from two subvolumes to capture submicron structure (**Fig. 1j**) over the entire ^~^250 μm sized whole organism. The improved axial resolution of the triple-view reconstruction offered clearer views of subnuclear structure than did the individual views (**Fig. 1k**), which were again distorted due their anisotropic PSFs.

### Improved imaging in larger, scattering samples

Confocal microscopy is often used to image large, densely labeled samples, as the pinhole enhances contrast due to the rejection of out-of-focus background and scattered emission. A challenge, however, is the depth-dependent attenuation of both illumination and emission in scattering tissue. Encouraged by our results on the L1 stage *C. elegans*, we sought to image an adult *C. elegans* animal, which ^~^30-40 fold larger in volume than an L1. Despite an invariant embryonic cell lineage^29^ and decades of work on this well-characterized model organism, it has not been possible to segment all nuclei in an intact adult *C. elegans* animal (let alone identify all cells), in part because of optical imaging difficulties in resolving neighboring nuclei in the deeper tissues. We reasoned that the complementary coverage provided by our triple-view geometry could allow us to alleviate scattering losses, especially compared to single-view imaging. To investigate this capability, we fixed and imaged an adult *C. elegans* nematode stained for nuclei with NucSpot Live 488, stitching and reconstructing the entire worm volume spanning ^~^871 μm x 125 μm x 56 μm from ten imaging subvolumes (**Fig. 2a, S2a, Movie S2**).

**Fig. 2.**
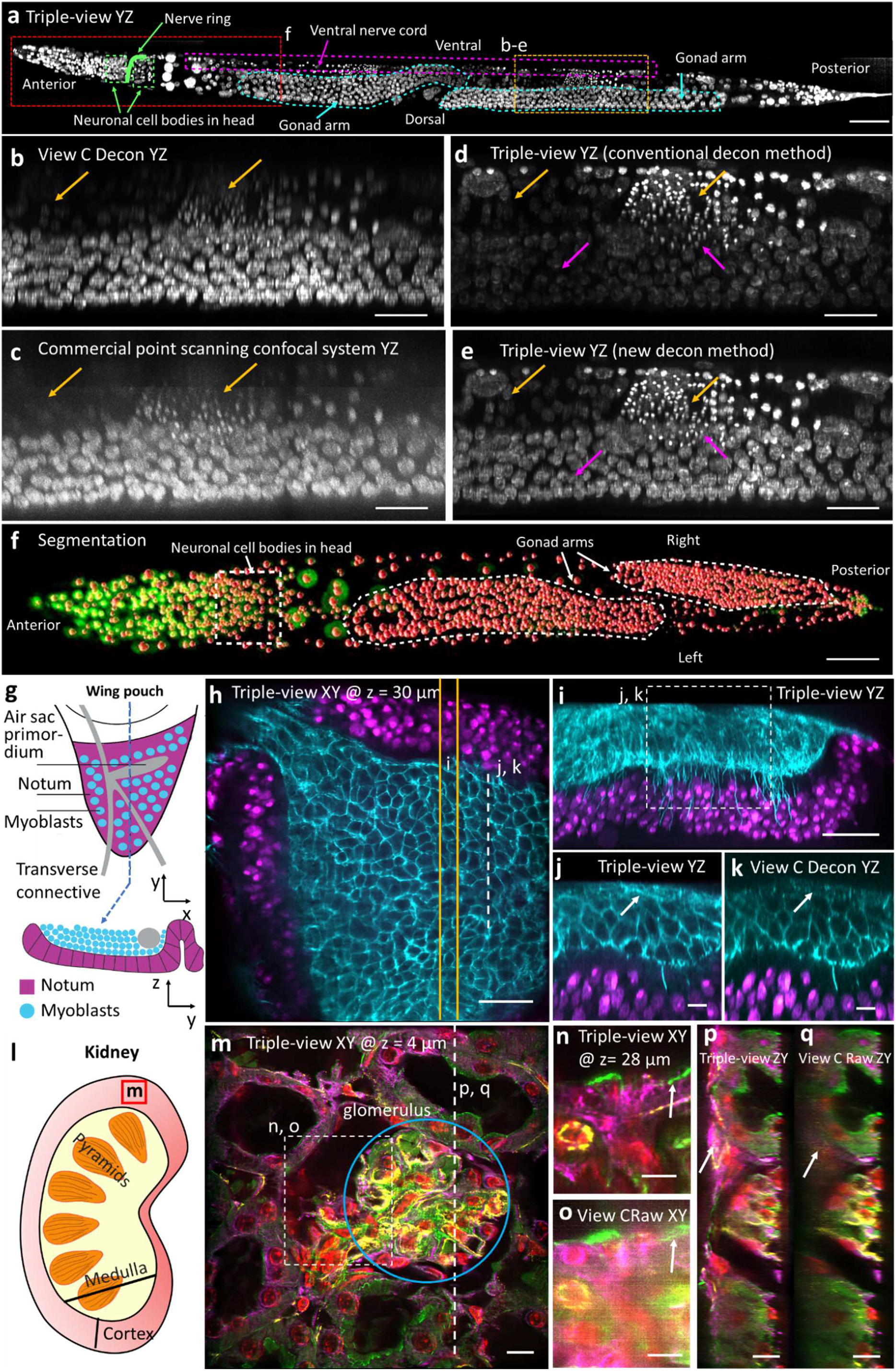
Multiview confocal microscopy offers improved imaging in scattering samples. **a)** Triple-view reconstruction of whole fixed adult *C. elegans*, labeled with NucSpot Live 488. Axial maximum intensity projection is shown. **b-e)** Comparative higher magnification views of dashed yellow rectangular region in **a)**, with bottom deconvolved view **b)**, commercial Leica SP8 confocal microscope **c)**, conventional triple-view deconvolution **d)**, attenuation-compensated triple-view deconvolution **e)**. Colored arrows highlight comparisons, orange: single-vs. triple-view, magenta: deconvolution methods. **f)** Triple-view reconstruction (green) with segmented nuclei overlaid in red (red), corresponding to red dashed rectangular region in **a)**. See also **Movie S2. g)** Schematic of larval wing disc, lateral (top) and axial (bottom) views, including adult muscle precursor myoblasts and notum. **h)** Lateral plane from triple-view reconstruction, 30 μm from sample surface. Notum nuclei (NLS-mCherry, magenta) and myoblast membranes (CD2-GFP, cyan) labeled. **i)** Axial maximum intensity projection derived from 6 μm thick yellow rectangle in **h). j, k)** Higher magnification view of white dashed line/rectangle in **h/i)**, comparing triple-view result **j)** to single View C deconvolution **k)**. White arrows: membrane observed in **j)** but absent in **k). l)** Schematic of kidney: approximate region where tissue was extracted. **m)** Four color triple-view reconstruction of mouse kidney slice. Lateral image at indicated axial height from beginning of the volume, highlighting glomerulus surrounded by convoluted tubules. Red: nuclei stained with DAPI, green: actin stained with phalloidin-Alexa Fluor 488; magenta: tubulin immunolabeled with mouse-α-Tubulin primary, α-Mouse-JF549 secondary; yellow: CD31 immunolabeled with Goat-α-CD31 primary, α-Goat AF647 secondary. **n, o)** Higher magnification of white rectangular region in **m)** at indicated axial distance; triple-view reconstruction **n)** vs. single view C **o). p, q)** Axial view along dashed line in **m)**, comparing triple-view reconstruction **p)** to single view C **q)**. White arrows: structures that are dim in single view but restored in triple-view. Scale bars: **a)** 50 μm, **b-e, n-q)** 10 μm, **h, i, m)** 20 μm, **j, k)** 5 μm.

In this sample, we noticed pronounced depth-dependent attenuation in the raw views acquired by our system, (**Fig. 2b**). The same degradation was also evident when imaging the same sample on a state-of-the-art point-scanning confocal microscope (**Fig. 2c**), which also imaged the whole sample ^~^26-fold more slowly (^~^70 mins) than our triple-view approach (^~^160 s for all three views). Fusion of all three views improved signal at the periphery of the imaging volume, where attenuation is minimal in any single view. However, signal in the interior of the animal still appeared dim (**Fig. 2d**). To remedy this, we modeled the signal attenuation as exponential with depth, estimating the decay parameter directly from the data, and explicitly incorporated the decay in our deconvolution algorithm (**Fig. S2b, Methods**). This procedure allowed us to compensate for the attenuation, producing clear reconstructions throughout the sample volume (**Fig. 2e**). Image quality and spatial resolution in the attenuation-compensated result were sufficiently good that we could use a modified deep learning routine based on Mask-RCNN^30,31^, in conjunction with manual editing, to segment 2136 nuclei in the worm (**Fig. 2f, Methods, Movie S2**). Such holistic segmentation was impossible using either the point-scanning data (**Fig. 2c**) or the conventional triple-view fusion (**Fig. 2d**).

In another example, we examined the notum region of *Drosophila* larval wing imaginal discs. The basal side of the wing disc columnar epithelium in the disc notum is a complex niche of two different adult progenitor cells: tracheoblasts that form the air-sac primordium (ASP), and myoblasts that are adult flight muscle precursors (**Fig. 2g**)^32^. Using CD2-GFP to label myoblast membranes and NLS-mCherry to mark the notum, we obtained clear triple-view reconstructions of these cell populations throughout the ^~^60 μm thick volume in lateral (**Fig. 2h**) and axial (**Fig. 2i**) views, highlighting a population of membrane extensions similar to cytonemes^32^ that oriented towards the discs^33^. Axial views confirmed that fusing all three views (**Fig. 2j**) gave better reconstructions near the top surface of the image volume than either lower View C alone (**Fig. 2k**) or higher NA single-view spinning-disk confocal reconstructions of a similar sample (**Fig. S2c, d**).

Next, we isolated a 30 μm thick tissue section from fixed mouse kidney (**Fig. 2l**), immunolabeling four targets to highlight subcellular constituents of glomeruli and convoluted tubules (**Fig. 2m**). Again, the triple-view reconstruction provided subcellular detail throughout the stack volume (**Fig. 2n, p**), unlike the raw data collected from individual views (**Fig. 2o, q**), which exhibited so much attenuation that cells and blood vessels appeared absent at the far side of the stack.

Finally, we imaged mouse cardiac (**Fig. S2e**) and brain (**Fig. S2g**) tissue slices spanning ^~^0.5 x 0.5 mm^2^ in lateral extent. Two-color imaging marking nuclei and proteins produced reconstructions reminiscent of hematoxylin and eosin stain, but with volumetric information. Again, the triple-view reconstructions produced better lateral (**Fig. S2f**) and axial (**Fig. S2h**) views than did conventional single-view confocal microscopy.

### Two-step deep learning for denoising and resolution enhancement in living samples

Next, we investigated the use of our multiview system in living samples (**Fig. 3**). As a first example, we used MitoTracker Green and Lysotracker Deep Red to label mitochondria and lysosomes in human colon carcinoma (HCT-116) cells, acquiring two-color, triple-view volumes in 1.2 s, every 10 s, over a total of 60 volumes (**Fig. 3a, Movie S3**). Mitochondria and lysosomes were clearly defined in the triple view reconstructions. The clarity in axial views enabled us to discern transient interactions between lysosomes and mitochondria that appeared to result in mitochondrial fission (**Fig. 3b**), which were obscured in the raw confocal views (**Fig. S3a, b**). In a second example, we imaged contractile cardiomyocytes^34^ that we labeled with MitoTracker Red CMXRos, acquiring triple-view volumes in 2 s, every 20 s, for 100 volumes (**Fig. 3c, Movie S3**). As for the HCT-116 cells, triple-view reconstructions displayed better spatial resolution and contrast than the raw single-view confocal data (**Fig. S3c**).

**Fig. 3.**
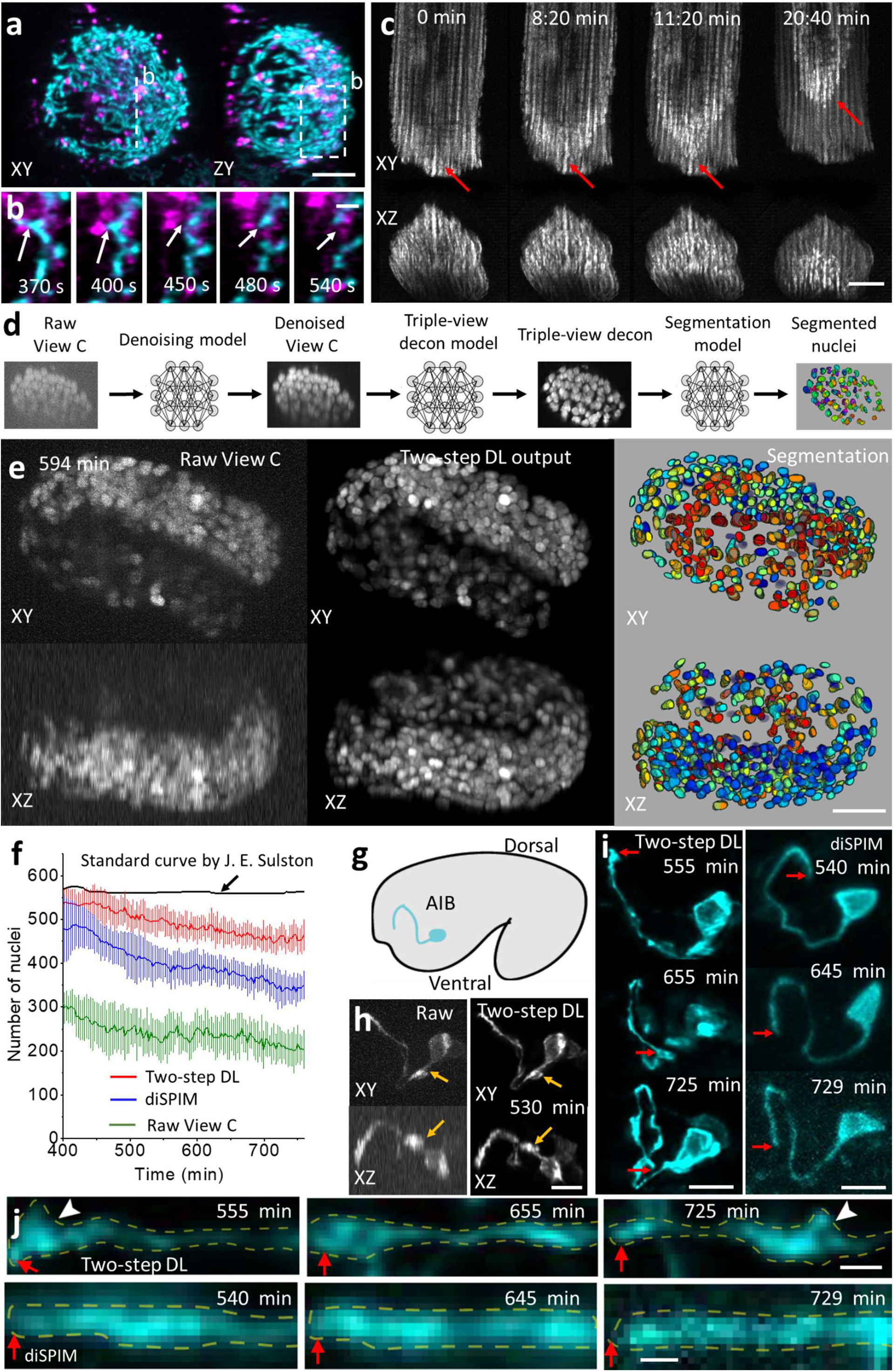
Multiview confocal imaging in living samples. **a)** HCT-116 human colon carcinoma cells labeled with MitoTracker Green (cyan) and LysoTracker Deep Red (magenta), imaged with triple-view CM. Triple view data acquired in 1.15 s, every 10 s, 60 time points. Lateral (left) and axial (right) maximum intensity projections are shown for a single time point after registration/deconvolution. **b)** Higher magnification of dashed line/rectangle in **a)**, highlighting association between lysosomes and mitochondria, and eventual mitochondrial fission (arrows). See also **Fig. S3a, b. c)** Cardiomyocytes expressing EGFP-Tomm20, labeled with MitoTracker Red CMXRos, imaged with triple-view CM. Triple view data were acquired in 2.03 s, every 20 s, 100 time points. Lateral (top) and axial (bottom) maximum intensity projections shown at indicated times after registration/deconvolution. See also **Movie S3. d)** Deep learning workflow. Raw data captured from objective C are subjected to three neural networks that successively denoise, enhance resolution, and segment nuclei. See **Fig. S3, Methods**, for further details. **e)** Example derived from live *C. elegans* embryos expressing H2B-GFP, at different stages of deep learning. Left: raw data from View C. Lateral (top) and axial (bottom) maximum intensity projections. Volumes acquired in 0.6 s, every 3 minutes, for 130 volumes. Middle: as in Left, but after denoising/ resolution enhancement CARE networks are applied. Right: after Mask-RCNN segmentation network and post-processing. Displayed segmentations (colored nuclei) are derived from 3D surface rendering. See also **Movie S4. f)** Quantifying nuclear segmentation from twitch to hatch on data acquired with raw View C (green), diSPIM (blue) and View C after two-step denoising and resolution enhancement (red), vs. number recorded by John E. Sulston in ref.^29^ Means and standard deviations are shown from 16 embryos. Times are minutes post fertilization, with twitching occurring at 435 minutes. **g)** Schematic of AIB neuron in *C. elegans* embryo. **h)** Example lateral (top) and axial (bottom) maximum intensity projections derived from single view volumetric input, comparing raw View C (left) vs. two-step deep learning (right) at indicated time post fertilization. AIB is marked with membrane targeted GFP. Single view volumes acquired in 0.75 s, every 5 min, for 65 volumes. Arrows: features better resolved after deep learning. Neurite widths in raw data: 470 ± 17 nm laterally, 1268 ± 127 nm axially; after deep learning: 355 ± 17 nm laterally and 754 ± 46 nm axially (*N* = 10). **i)** Additional comparisons highlighting elaboration of the AIB neurite (neurite tip indicated by red arrows) during development, two-step deep learning on View C input (left), vs. diSPIM images of a different embryo at similar times (right). **j)** Higher magnification of neurites in **i)**. Neurites straightened using ImageJ; corresponding neurite tip regions are indicated by red arrows in **i, j**. White arrowheads in **j)**: varicosities evident in deep learning prediction (top) but obscured in the diSPIM data. Yellow dashed lines outline neurite. Scale bars: **a)** 3 μm, **b)** 1 μm, **c, e)** 10 μm, **h, i)** 5 μm, **j)** 2 μm.

However, the cardiomyocyte reconstructions also illustrate drawbacks in our multiview confocal approach. Some reconstructed volumes showed obvious motion blur, indicating the need for more rapid imaging, which is made more difficult since three confocal views are serially acquired for each reconstruction. Additionally, we observed a progressive ‘front’ of fluorescence loss over time, indicating light-induced signal change (**Fig. 3c, Movie S3**). We had previously observed fluctuations in MitoTracker Red using light-sheet microscopy^20^, which we attributed to transient fluctuations in mitochondrial membrane population. Although the effects in this dataset (**Fig. S3c, d**) were similar, the more global loss in fluorescence is also suggestive of phototoxicity, which could lead to reduction in membrane potential and thus loss of mitochondrially-associated dye.

The requirement for rapid imaging on the one hand and minimal phototoxicity on the other is more stringent when imaging developing animals, e.g. *C. elegans* embryos^35^. Imaging is particularly difficult in the latter half of embryogenesis when muscular twitching causes the nematode to writhe within the eggshell. For example, motion blur, photobleaching, developmental delay or embryonic arrest confounded our attempts to obtain high SNR triple-view confocal recordings of a pan-nuclear, GFP-histone label over the seven-hour period from embryonic twitching to hatching. By recording from only the single 1.2 NA confocal view and lowering the illumination intensity, we improved imaging speed and recorded volumes every three minutes over the entire imaging period without obvious photobleaching or developmental delay. However, under these conditions poor SNR and scattering prevented visualization of many nuclei (**Fig. 3d, e**). These problems motivated us to develop computational solutions to restore our data.

Deep learning has emerged as a powerful, data driven method for denoising and resolution enhancement^36–38^. We trained and implemented sequential ‘content aware image restoration’ (CARE^36^) neural networks that (1) denoised the raw, low SNR lower view recordings and then (2) predicted the high SNR triple-view output, given this higher SNR single-view confocal input (**Fig. 3d, Fig. S3e-h**). Key to this two-step procedure is obtaining sufficient high quality training data, both for denoising and for resolution improvement. To achieve this goal, we paralyzed late-stage *C. elegans* embryos expressing the pannuclear label, then acquired 100 matched low-SNR (^~^4) and high-SNR (^~^60) volumes from all three views to train the denoising networks, and 68 matched volumes to train the triple-view deconvolution network, reserving 10 volumes for testing (**Methods, Fig. S3i**). Intriguingly, this two-step output produced better output than when trying to train a single network to predict high SNR triple-view output directly from noisy single-view raw data (**Fig. S3f, g**).

An advantage of the *C. elegans* embryonic system is that the number of nuclei is known exactly, as they were manually scored by John Sulston by visually examining many individual embryos^29^. This effort, achieved four decades ago, has not been repeated with fluorescence microscopy due to imaging challenges. Indeed, inspection of the noisy confocal single-view input revealed that scattering severely attenuated the signal of nuclei on the ‘far side’ of the embryo (i.e., furthest from the detection lens). Qualitatively, the two-step prediction appeared strikingly better than the single-view input, restoring many nuclei even at increasing depths within the embryo (**Fig. 3e, Movie S4**). To quantify the improvement, we then used a third neural network, our previous Mask RCNN (**Fig. 2f, Fig. S3i, j**), to segment and count the number of nuclei. Against the Sulston ground truth, the raw single confocal view found fewer than half of all nuclei. The two-step prediction fared much better, capturing the majority of the nuclei, outperforming single-view light-sheet microscopy data passed through a neural network designed to improve resolution isotropy^21^, and even dual-view light-sheet microscopy data (diSPIM^14,39,40^, **Fig. 3f, Fig. S3k**).

To examine the usefulness of our two-step restoration method in resolving nanoscale structures through time, we imaged the development of neurites in the *C. elegans* nerve ring, the equivalent of the nematode brain (**Fig. 3g**). We focused our analyses on the AIB interneurons, a pair of “functional hub”^41^ neurons which bridge communication between modular and distinct regions of the neuropil.^42^ The architecture of AIB is precisely laid out during development to bridge nerve ring neighborhoods, with a coordinated assembly of synaptic varicosities during the establishment of the circuits^43,44^. While the morphology of AIB had been known for three decades, the developmental events leading to its precise positioning in the brain remain unknown owing to limitations in optical imaging. We accumulated 45 matched volumes of low SNR single view input and high SNR triple-view data, using 40 volumes for training and 5 for testing. Denoising and triple-view prediction improved SNR, resolution, and contrast relative to raw single-view input data (**Fig. 3h, Fig. S3g**). We also compared the two-step predictions (**Fig. 3h**) to another embryo imaged over the same period with diSPIM, imaging the elaboration of the AIB neurite over a span of 450 minutes (**Fig. 3i**). The superior resolution offered by the deep learning prediction enabled us to detect, in developing embryos, growth cone dynamics at the tip of the neurite and the emergence of presynaptic boutons as the neurite engaged with its postsynaptic partners (**Fig. 3j**). The fluorescent intensity observed for the growth cone labeling, using our cytoplasmic marker^45^, decreased as the AIB neurite was correctly positioned into its final neighborhood, concomitant with the emergence (and increase of intensity) of presynaptic boutons, indicating a developmental transition during synaptogenesis. Other imaging approaches, including diSPIM, were incapable of capturing the subcellular structural changes described here for a single neurite and in the context of the developing brain (**Fig. 3j**).

### Multiview super-resolution imaging

The discrepancy in size between the optical point spread function and a fluorescently labeled target usually hinders the observation of subcellular phenomena with diffraction-limited microscopy. To ameliorate this problem, we next sought to integrate super-resolution methods into our imaging platform. Within the umbrella of super-resolution microscopy techniques, linear structured illumination microscopy (SIM^46^) stands out for its ability to provide resolution enhancement at rapid speed and relatively low dose, enabling extended volumetric super-resolution imaging in living cells. In SIM, sharp, sparse illumination is used to encode otherwise inaccessible super-resolution information into the passband of the microscope, where it is decoded using digital or optical processing. We reasoned that the diffraction-limited line structure in our microscope could be used for SIM, provided we modified our acquisition scheme to isolate sparse fluorescence emission in the vicinity of the line focus (**Fig. 4a, b**). Therefore, we (1) used an acousto-optic tunable filter (AOTF) to periodically shutter the beam during each confocal image, producing sparse illumination structure; (2) phase shifted the structured illumination in the direction of the line scan, collecting the resulting images; (3) developed post-processing schemes to detect the positions of fluorescence emissions and reassign them^8,9^, thereby improving spatial resolution in the direction of line scanning (**Fig. 4b, Methods, Fig. S4a-h**). Performing this procedure in any single view improved spatial resolution in 1D. However, the three objectives in our microscope allowed us to scan the illumination line along three orthogonal directions (**Fig. 4a**). Collecting and processing five images per direction enabled three-dimensional super-resolution imaging, improving volumetric resolution 5.3-fold, from 335 x 285 x 575 nm^3^ to 225 x 165 x 280 nm^3^ in triple-view 1D SIM mode, as verified on immunolabeled microtubules in fixed U2OS cells (**Fig. 4b**).

**Fig. 4.**
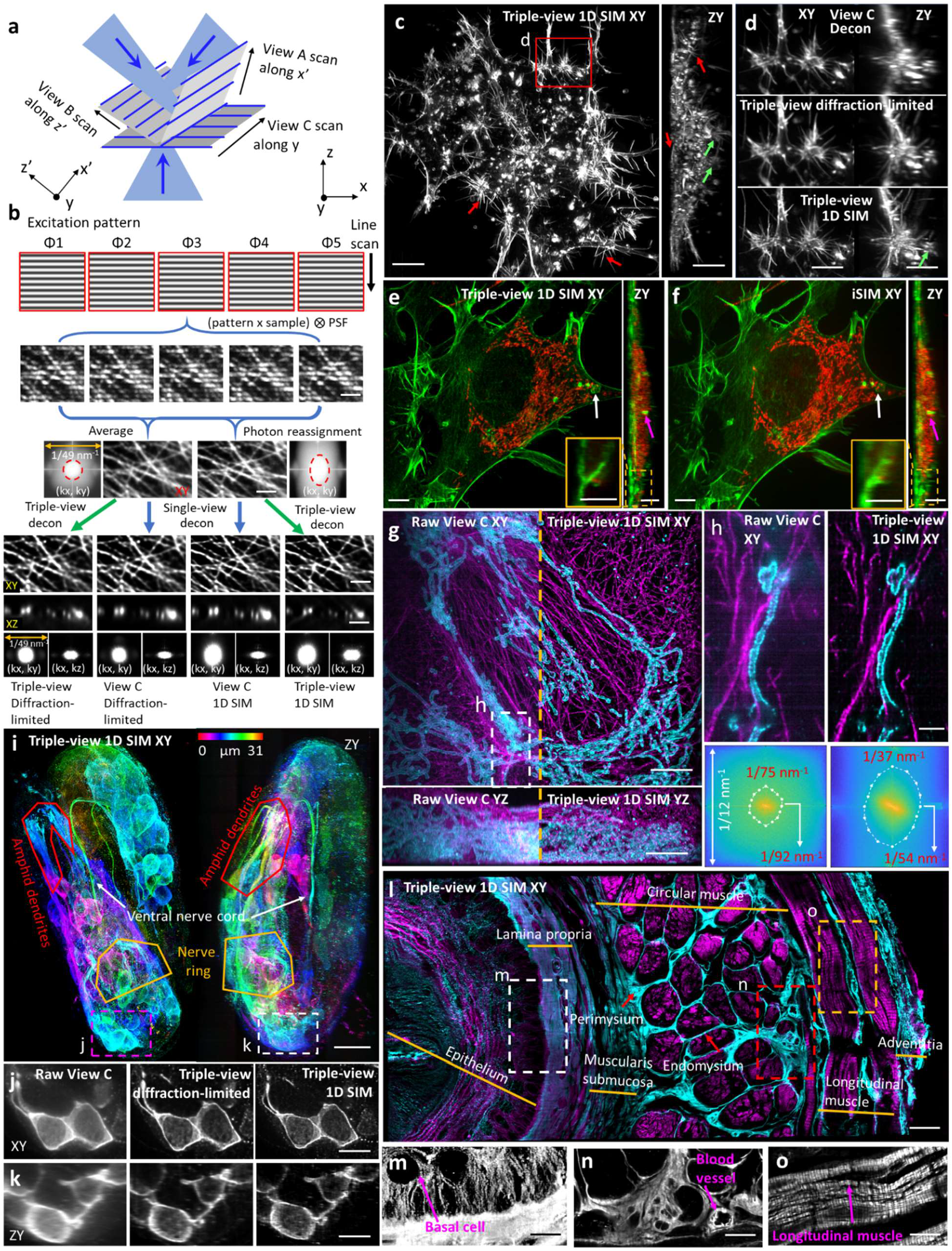
Multiview super-resolution microscopy in fixed cells and tissue. **a)** Illumination geometry. Sparse, periodic line illumination is scanned in orthogonal directions (x’, y, z’), enhancing 3D resolution. **b)** Workflow for a single scanning direction. Five confocal images are acquired per plane, each with illumination structure shifted 2π/5 in phase relative to the previous position. Averaging produces diffraction-limited images. Detecting each illumination maximum and reassigning the fluorescence signal around it (photon reassignment) improves spatial resolution in the direction of the line scan. Combining image volumes acquired from multiple views further improves volumetric resolution. Images are of immunolabeled microtubules in fixed U2OS cells and corresponding Fourier transforms. **c)** Triple-view SIM lateral (left) and axial (right) maximum intensity projections of fixed B16F10 mouse melanoma cells labeled with Alexa Fluor 488 phalloidin and embedded in 1.8% collagen gel. Red/green arrows highlight filopodia/invadopodia. See also **Movie S5. d)** Higher magnification lateral (left) and axial (right) views corresponding to red rectangular region in c), comparing deconvolved view C (top), diffraction-limited triple-view deconvolution (middle), and triple-view SIM result. See also **Movie S5. e, f)** Comparative triple-view SIM **e)** and instant SIM **f)** maximum intensity projections of the same fixed HEY-T30 cell embedded in collagen gel, labeled with MitoTracker Red CMXRos and Alexa Fluor 488 phalloidin. White and magenta arrows: the same features in lateral (left) and axial (right) projections. Insets are higher magnification axial views of dashed rectangular regions. **g)** Comparative raw single view C (left of dashed orange line) and triple-view SIM (right of dashed line) maximum intensity projections of fixed and 4x expanded U2OS cell, immunolabeled with rabbit anti-Tomm20 primary, anti-rabbit-biotin, streptavidin Alexa Fluor 488 marking mitochondria (cyan) and mouse anti-alpha tubulin primary, anti-mouse JF 549, marking tubulin. Lateral (top) and axial (bottom) views are shown. **h)** Higher magnification views of white dashed region in **g)**, comparing triple-view (left) and triple-view SIM (right) reconstructions. Bottom: sector-based decorrelation analysis to estimate spatial resolution in images at top. Resolution values for horizontal and vertical directions are indicated. **i)** Lateral (left) and axial (right) triple-view SIM maximum intensity projections from fixed, 3.3-fold expanded *C. elegans* embryo, immunostained with rabbit-anti-GFP, anti-rabbit-biotin, and streptavidin Alexa Fluor 488 to amplify GFP signal marking neuronal structures as indicated. Images are depth coded as indicated. Higher magnification slices of lateral (**j**) and axial (**k**) views indicated by magenta and white dashed rectangles in **i)**, illustrating progressive improvement in resolution from raw single view (left), triple-view deconvolved (middle), and triple-view 1D SIM mode (right). **l)** XY image of mouse esophageal tissue slab, immunostained for tubulin (mouse-anti-alpha tubulin, anti-mouse JF 549, cyan) and actin (phalloidin Alexa Fluor 488, magenta), imaged in triple-view SIM mode. Anatomical regions are highlighted. **m, n, o)** Higher magnification single-color views of white (**m**, tubulin), red (**n**, tubulin), and orange (**o**, actin) dashed regions in **l)**, anatomical features highlighted. Contrast has been adjusted in **m**) to better observe outline of basal cell. Scale bars: **b, j, k)** 2 μm, **c, m-o)** 10 μm, **d-g, i)** 5 μm, **h)** 1 μm, **l)** 20 μm.

An additional advantage of this implementation of SIM is its ability to interrogate thicker, more densely labeled samples due to the strong suppression of out-of-focus light. For example, when imaging Alexa Fluor 488 phalloidin-labeled actin in fixed B16F10 mouse melanoma cells embedded in a 1.8% collagen gel^47^, the triple-view SIM reconstruction offered clear views of three-dimensional actin-rich filopodia and invadopodia (**Fig. 4c**), especially those that protruded in the axial direction. Comparing the triple-view 1D SIM results to the raw single-view or triple-view confocal results confirmed this impression, also highlighting the progressive improvement in spatial resolution (**Fig. 4d, Movie S5**). We observed similar gains on fixed and immunolabeled *C. elegans* embryos (**Fig. S4i-k**). In another example, we imaged a fixed, collagen gel embedded HEY-T30 ovarian cancer cell dual-labeled for actin and mitochondria in triple-view 1D SIM mode (**Fig. 4e**), acquiring comparative images of the same sample using instant SIM^10^, a rapid single-view SIM implementation (**Fig. 4f**). Although the lateral resolution of both SIM implementations was similar, our triple-view method provided better axial resolution due to the super-resolution capability introduced by the upper two specimen views.

Triple-view 1D SIM also improved images of physically expanded samples (**Fig. 4g-j**). Triple-view SIM reconstructions on 4x expanded U2OS cells immunostained for Tomm20 and microtubules showed better contrast (**Fig. 4g**) and resolution (**Fig. 4h**) relative to single-view input. For example, using decorrelation analysis to compare spatial resolution, we obtained ^~^40-50 nm lateral resolution in SIM mode vs. 60-80 nm resolution in confocal mode (**Fig. 4h**). To investigate performance in expanded multicellular samples, we adapted and optimized methods^48,49^ to expand *C. elegans* embryos 3.3-fold (**Methods, Fig. S5**), imaging dense neural markers in triple-view 1D SIM mode to visualize amphid neurites, the nerve ring, and ventral nerve cord (**Fig. 4i**). Comparisons on cellular membranes with raw single- and triple-view data highlighted the improvement offered in the SIM mode, both laterally (**Fig. 4j**) and axially (**Fig. 4k**).

Finally, we immunolabeled actin and microtubules in fixed mouse esophageal tissue, then imaged and reconstructed the triple-view SIM output over a 172 x 365 x 25 μm^3^ slab by stitching together 21 subvolumes (**Fig. 4l**). At larger length scales, we identified distinct, radially arranged anatomic regions, including epithelia, lamina propia, submucosa, perimysium, endomysium, longitudinal muscle and adventitial tissue layers. At smaller length scales, the subcellular detail afforded by our reconstruction also enabled crisp visualization of cells, small vessels, and the striated, periodic actin arrangements within longitudinal muscle (**Fig. 4m-o**).

### Deep learning enables isotropic in-plane super-resolution imaging from confocal input

Operating our line confocal system in SIM mode enables 1D super-resolution imaging in any individual volumetric view, requiring three views for 3D super-resolution. Given that five diffraction-limited images are required per view, the 3D SIM mode thus requires 5x more images than the triple-view confocal mode, and 15x as many images as conventional single-view confocal microscopy. Although acceptable when imaging brightly labeled, fixed samples (**Fig. 4**), the need for extra images hampers live imaging applications due to the associated increase in illumination dose and decrease in temporal resolution. We thus considered additional technologies to achieve super-resolution, particularly those capable of significantly improving imaging speed or duration.

Deep learning offers such a method, especially given that 1) robust training data consisting of matched confocal input and 1D super-resolved output can be readily obtained by our microscope (**Fig. 4b**); and 2) the desired resolution enhancement (^~^2-fold better than the confocal input) is relatively modest. If the training data is assembled from samples arranged in random orientations, as is typical when imaging biological specimens, we reasoned that 1D resolution could be improved in any arbitrary direction relative to the input data simply by rotating the input image and reapplying the trained neural network. Combining all such rotated, super-resolved views using joint deconvolution could then enable isotropic, in-plane resolution enhancement (**Fig. 5a**), from a single confocal input image.

**Fig. 5.**
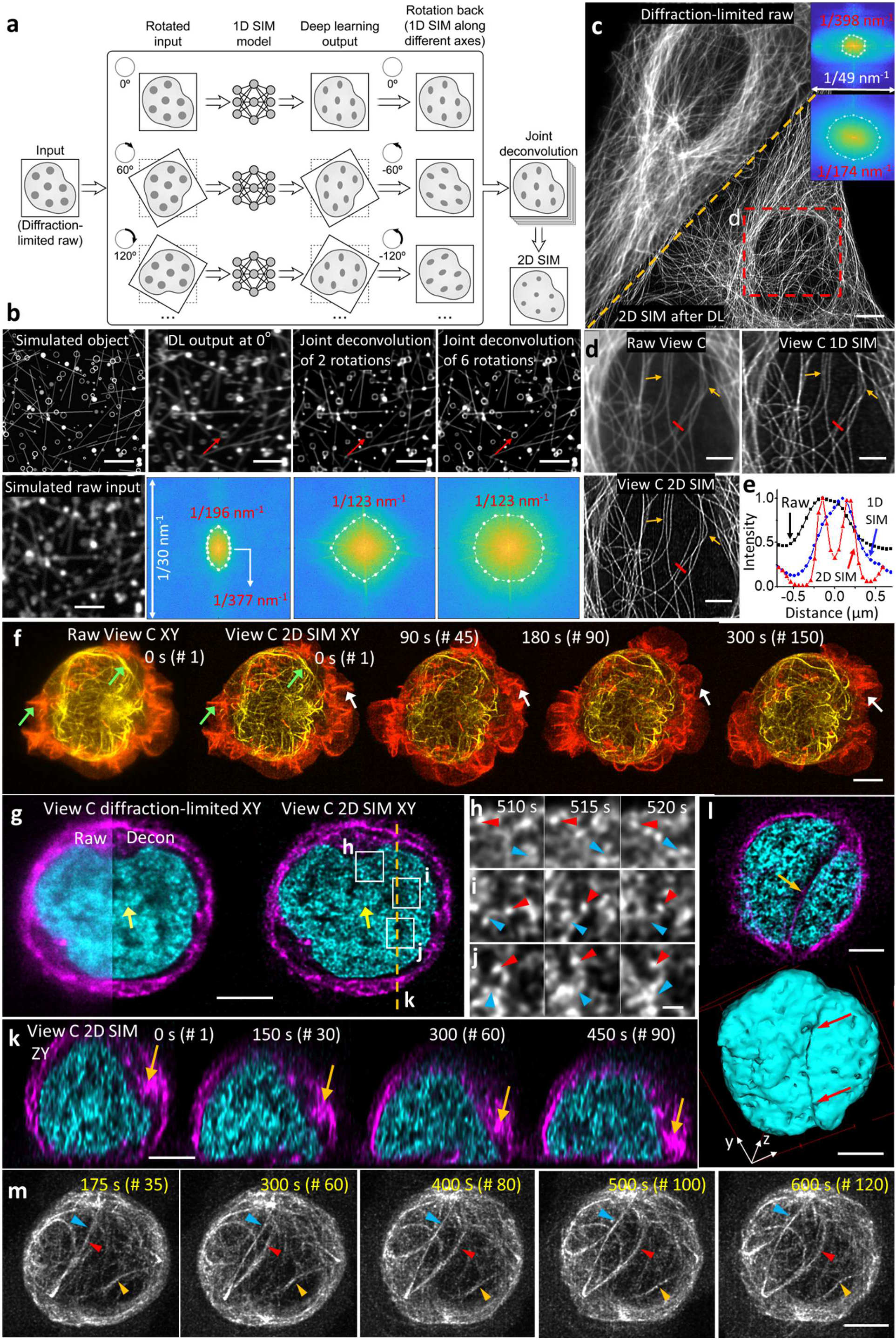
Deep learning for isotropic in-plane super-resolution imaging. **a)** Deep learning and joint deconvolution produce 2D super-resolution images from diffraction-limited input. A library of diffraction-limited and matched 1D SIM super-resolved image volumes is used to train a neural network to produce 1D super-resolved images. Given new diffraction-limited images rotated with respect to each other, application of the neural network produces rotated 1D super-resolved images. Rotating these images back to the initial reference frame and using joint deconvolution to combine them produces a 2D SIM super-resolution image. **b)** Simulations of the method in **a)**. An object consisting of lines and hollow spheres can be blurred to resemble diffraction-limited confocal input (left column). Outputs after deep learning (DL) at a single orientation (second column), after joint deconvolution of two orientations (third column), or six orientations (last column) show progressive improvement in resolution isotropy (red arrows), confirmed with Fourier transforms of the images (second row). Ellipses bound decorrelation estimates of resolution (numerical values from ellipse boundary indicated in red text). See also **Fig. S6a. c)** CM volumes of immunolabeled microtubules (above of dashed line) in fixed U2OS cells, processed as in **a)** to produce 2D SIM predictions, improving lateral spatial resolution (below dashed line). Fourier transforms in inset show improvement in spatial resolution to ^~^170 nm after deep learning and joint deconvolution. See also **Fig. S6b. d)** Higher magnification view of red dashed region in **c)**, comparing raw confocal input, 1D SIM output after deep learning and deconvolution, and 2D SIM output after deep learning and joint deconvolution after six rotations. Arrows highlight regions for comparison. **e)** Profiles along red line in **d)**, comparing the resolution of two filaments in 2D SIM mode (red), 1D SIM mode (blue), and confocal input (black). **f)** Lateral maximum intensity projections of Jurkat T cell expressing EMTB-3XGFP (yellow) and F-tractin-tdTomato (red), volumetrically imaged every 2 s, as imaged with raw line confocal input (View C, left) and 2D SIM output after deep learning and joint deconvolution with six rotations. Four time points from 150 volume series are shown; green arrows highlight features better resolved in 2D SIM output vs. raw input; white arrows indicate actin dynamics at cell periphery. See also **Movie S6. g)** Single lateral plane within Jurkat T cell expressing H2B-GFP (cyan) and EMTB-mCherry (magenta), volumetrically imaged every 5 s for 200 time points. Diffraction-limited View C input (left) is compared to 2D SIM output (right), with View C raw data also compared to deconvolved data. Yellow arrow indicates fine nuclear structure for comparison. **h-j)** Higher magnification views of white rectangular regions in **g)** illustrating dynamic movements of H2B nanodomains (red arrowheads: the same nanodomain; blue arrowheads: splitting or merging of nanodomains). Images are maximum intensity projections over 500 nm axial range. **k)** Axial plane corresponding to dashed orange line in **g**), highlighting centrosome movement towards coverslip (orange arrow) and accompanying nuclear shape distortion. See also **Fig. S6d. l)** Lateral plane (top) and surface rendering (bottom) of nuclear indentation by microtubule bundles. **m)** Maximum intensity projections over bottom half of EMTB-mCherry volumes at indicated times, showing coordinated movement of the microtubule cytoskeleton. Colored arrowheads mark regions for comparison. See also **Movie S6**. Scale bars: **b)** 2 μm, **c, f, g, k-m)** 5 μm, **d)** 3 μm, **h-j)** 500 nm.

We investigated the validity of such an approach using phantom objects, consisting of hollow spheres and lines with subdiffractive width (**Fig. 5b**). Blurring the phantoms to resemble confocal data and producing matched 1D SIM counterparts allowed us to train a 3D residual channel attention network (RCAN), recently shown to enable resolution improvement in fluorescence microscopy image volumes^38^. Applying the trained network produced 1D resolution enhancement (**Fig. S6a**) but rotating the input image, reapplying the network, and combining the associated network output with joint deconvolution isotropized the resolution gain (**Fig. 5b**). We then evaluated the method on experimentally acquired images of immunolabeled microtubules in fixed U2OS cells (**Fig. 5c, Fig. S6b**). We gathered 50 volumes of matched confocal and 1D SIM data, trained an RCAN, and applied the trained network to unseen data, jointly deconvolving the output of six rotations for near isotropic output (**Fig. 5c, d**). Decorrelation analysis showed average spatial resolution improved from 386 ± 42 nm (raw confocal input) to 176 ± 8 nm (*N* = 10 xy slices, **Fig. 5c**), confirming our visual impression that filaments were better resolved in 2D SIM mode than the 1D SIM deep learning output or confocal input (**Fig. 5e**).

To demonstrate the potential for interrogating subcellular dynamics over extended duration, we imaged live, fluorescently labeled Jurkat T cells, often used to model the formation and signaling of the immune synapse^50–52^. We imaged Jurkat cells transiently transfected with EMTB-3XGFP and F-actin tdTomato, and activated on antibody-coated coverslips, recording 150 single-view volumes at two second intervals. When we applied RCANs trained from 124 volumes (73 fixed cells expressing EMTB-3XGFP and 51 fixed cells expressing F-actin tdTomato) and performed joint deconvolution on six rotations, we saw improved resolution and contrast relative to the confocal input, and clearly resolved ruffling actin dynamics at the periphery of the T cell as well as microtubule dynamics closer to the cell center (**Fig. 5f, Movie S6**).

In another example, we applied the same training and deconvolution scheme to Jurkat T cells expressing H2B-GFP (cyan) and 3xEMTB-mCherry (magenta, marking microtubules), volumetrically imaged every 5 s for 200 time points during activation (**Movie S6**). The superior resolution offered by the 2D SIM output revealed H2B nanodomains that were obscured in the raw confocal data and barely evident in deconvolved raw data (**Fig. 5g**). Close inspection of the nanodomains revealed dynamic splitting and merging (**Fig. 5h-j**). Intriguingly, the size and dynamics of these nanodomains were consistent with 2D observations of ‘chromatin blobs’ imaged in live U2OS cells^53^. We visualized relocation of the centrosome towards the cell-substrate contact zone in axial views of the microtubule channel, as well as concomitant squeezing and deformation of the nucleus as the cell spread on the activating surface (**Fig. 5k, Fig. S6d**). In addition to gross nuclear shape changes, we also observed indentation of the nucleus near microtubule bundles (**Fig. 5l**). Such deformations can induce chromatin reorganization, capable of altering gene expression during hematopoietic differentiation^54^ and disrupting nucleocytoplasmic transport in Tau-mediated frontotemporal dementia neuronal cells^55^. Whether microtubule-induced nuclear shape changes affect T cell activation is an open question. Finally, we observed that microtubules wrap tightly around the nucleus (**Fig. 5m**), and that the centrosome produces a concave deformation, pulling and rotating the nucleus as it becomes docked at the immune synapse (**Fig. S6d**). To the best of our knowledge such coherent motion of the centrosome and nucleus has not been reported. The high resolution and fast volumetric imaging possible with this system will enable studies of how the dynamics of cytoskeletal proteins correlate with and modulate changes in chromatin structure and dynamics.

### Deep learning enhances multiview super-resolution imaging

The ability to 1) combine diffraction-limited volumetric views (‘triple-view deconvolution’); 2) produce 1D SIM in each volumetric view, combining the associated super-resolved views by joint deconvolution (‘triple-view 1D SIM’, **Fig. 4**); and 3) employ deep learning to either provide 2D super-resolution imaging in any individual view (‘2D SIM’, **Fig. 5**), or combine such volumetric views (‘triple-view 2D SIM’) enables a highly versatile imaging platform (**Fig. 6a**). Given the substantial improvement in volumetric resolution theoretically expected for the triple-view 2D SIM mode vs. the single-view high NA confocal view, we benchmarked the different microscopy modes against each other on biological specimens (**Fig. 6b-p, Fig. S6d, e**).

**Fig. 6.**
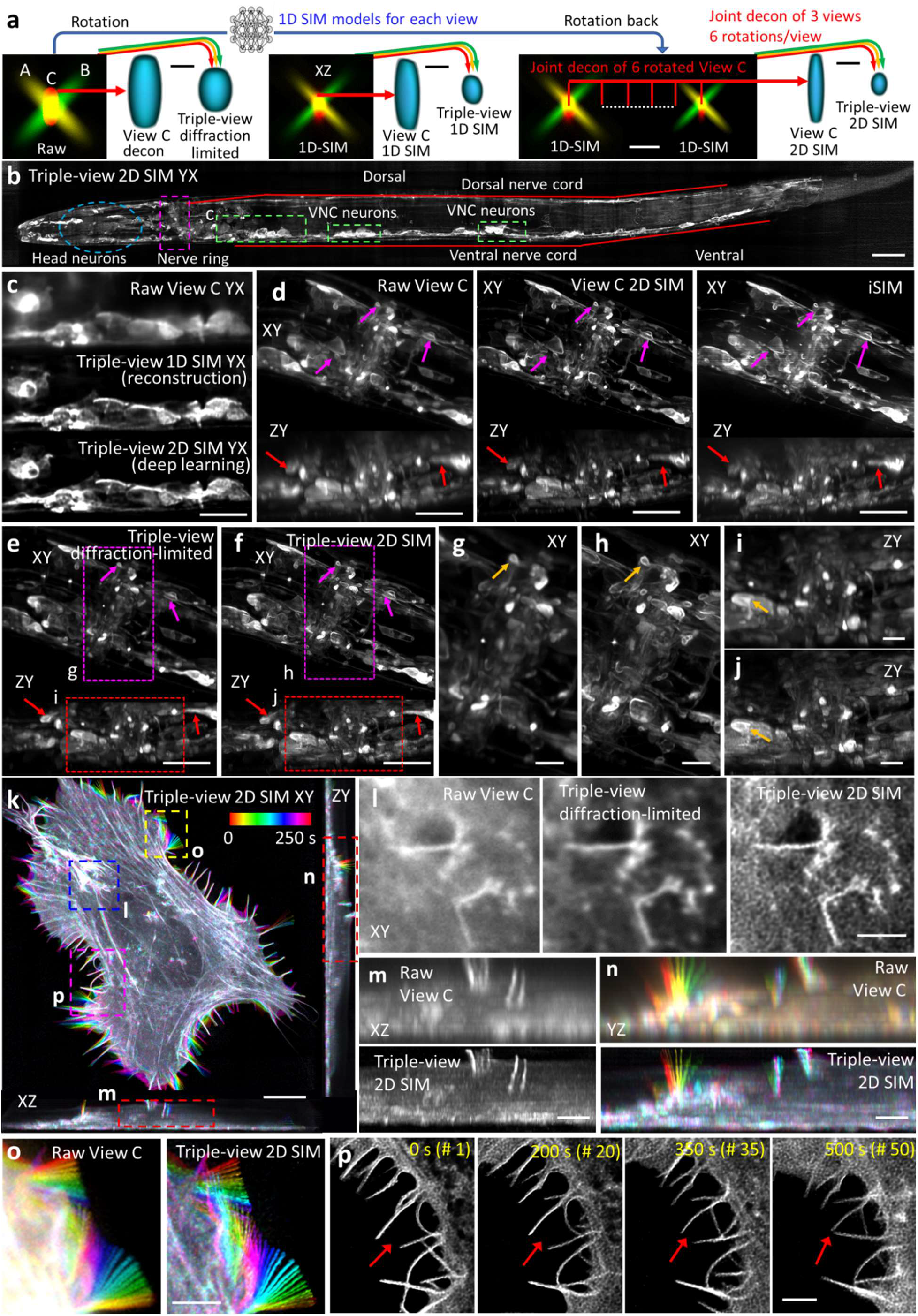
Deep learning enhances multiview super-resolution imaging. **a)** Different methods of combining data. *Left:* Diffraction limited volumes acquired from views A (yellow), B (green), C (red) may be combined with joint deconvolution to yield triple-view diffraction-limited data (RYG arrows). *Middle:* Alternatively, 5 volumes per view may be collected and processed as in **Fig. 4b** for 1D SIM, and the 3 1D SIM volumes combined using joint deconvolution to reconstruct triple-view 1D SIM data. *Right:* Instead, confocal data from each view may be passed through 1D SIM networks, and the data combined via joint deconvolution. Combining 6 rotations (for clarity only two are shown in figure) from view C yields View C 2D SIM data (**Fig. 5**). If the procedure is repeated for Views A, B, and the data combined with joint deconvolution, triple-view 2D SIM data may be obtained. XZ cross sections through PSFs are shown in black boxes. Blue volumes are relative sizes of PSFs that result from each process. **b)** Maximum intensity projection of fixed L2 stage larval worm expressing membrane targeted GFP primarily in the nervous system, imaged in triple-view 2D SIM mode. Anatomy as highlighted. **c)** Higher magnification views (single slices 6 μm into volume) of dashed cyan rectangle, highlighting VNC neurons as viewed in diffraction-limited View C (upper), triple-view 1D SIM obtained by processing 15 volumes (5 per view, middle), and triple-view 2D SIM mode (3 volumes, 1 per view, lower). **d)** Lateral (upper) and axial (bottom) maximum intensity projections from anesthetized L4 stage larval worm expressing the same marker, comparing dense nerve ring region imaged in diffraction-limited View C (left), View C 2D SIM mode (middle), and iSIM (right). Purple, red arrows highlight labeled cell bodies or membranous protrusions for comparison. **e, f)** As in **d)**, but comparing triple-view diffraction-limited vs. triple-view 2D SIM. Higher magnification lateral (**g, h**) and axial (**i, j**) views of purple, red dashed regions in **e, f)** are also shown, with orange arrows indicating regions for comparison. See also **Movie S7. k)** Example lateral (left) and axial (right, bottom) maximum intensity projection of U2OS cell expressing Lifeact tdTomato, volumetrically imaged in triple-view 2D SIM mode every 10 s for 100 time points. Images are color coded to indicate temporal evolution. **l)** Higher magnification single plane view of blue dashed region in **k)**, comparing raw View C (left), triple-view diffraction-limited (middle) and triple-view 2D SIM images. **m, n)** Higher magnification axial views of dashed red regions in **k)**, comparing raw View C vs. triple-view 2D SIM images. Note that **m)** shows single time point, **n)** a temporal projection. **o)** Higher magnification view of yellow dashed rectangular region in **k**), color coded to illustrate filopodial dynamics and comparing raw View C (left) and triple-view 2D SIM (right) images. **p)** Higher magnification views of pink rectangular region, highlighting filopodial dynamics. Maximum intensity projection is derived from 500 nm axial extent. Red arrow indicates filopodia moving together. See also **Movie S7**. Scale bars: **a)** white: 1 μm, black: 200 nm, **b, d-f, k)** 10 μm, **c)** 5 μm, **g-j, l-p)** 3 μm.

In a first example, we imaged a fixed, 18 μm thick L2 stage *C. elegans* larva expressing a GFP membrane marker primarily in the nervous system (**Fig. 6b**). Compared to the raw confocal View C, the triple-view 1D SIM result derived from 15 volumes (5 volumes per view, 3 views) showed clearly improved resolution and contrast (**Fig. 6c**). To investigate the performance of the triple-view 2D SIM mode, we collected 75 matched confocal and 1D SIM volumes of the same marker from 25 worms, allowing us to train an RCAN for 1D super-resolution prediction. Applying the RCAN enabled us to derive the triple-view 2D SIM result (**Fig. 6b, c**), which closely resembled the triple-view 1D SIM reconstruction, despite requiring only 3 input volumes (i.e., one for each view). Both triple-view 1D- and 2D SIM results outperformed a commercial OMX 3D SIM system, which showed obvious artifacts likely arising from out-of-focus contamination in this relatively thick sample (**Fig. S6c**).

In a second example, we imaged the densely labeled nerve ring region in a fixed, 28 μm thick L4 stage *C. elegans* larva embedded in a polymer gel with the same refractive index as water^56^ (**Fig. 6d**), first comparing the raw View C confocal input and the associated 2D SIM output derived from deep learning. The 2D SIM reconstruction was obviously sharper than the confocal input, better resolving cell bodies and fine neural structure, including membrane protrusions. We also obtained comparative images of the same sample using iSIM, finding that the 2D SIM mode provided similar lateral but better axial views of the specimen, perhaps because of the superior suppression of out-of-focus signal offered by our line confocal acquisition. However, all these single-view methods suffered worsened axial resolution and deterioration at the far side of the larvae, as expected in this thicker volume. To remedy this, we also acquired confocal views from the other two views, using them to obtain triple-view deconvolved (**Fig. 6e**) and triple-view 2D SIM reconstructions (**Fig. 6f**). As expected, both triple-view reconstructions (**Fig. 6e, f**) offered more nearly isotropic images than their single-view counterparts (**Fig. 6d**), also better defining the upper region of the image volume. The triple-view 2D SIM mode offered superior resolution in both lateral (**Fig. 6g, h**) and axial (**Fig. 6i, j**) views, evident in volumetric reconstructions (**Movie S7**). Quantifying spatial resolution via decorrelation analysis confirmed the superior volumetric resolution obtained in triple-view 2D SIM mode, more than ten-fold better than the raw confocal data (**Fig. S6d**).

Finally, we reconstructed actin dynamics in a U2OS cell transiently transfected with Lifeact tdTomato, imaging three confocal views every 10 s for 100 time points (**Fig. 6k, Movie S7**). Lateral views confirmed the progressive increase in resolution from raw View C to triple-view diffraction-limited to triple-view 2D SIM output (**Fig. 6l**), with axial views of the latter showing much clearer views of fine, vertically oriented protrusions than the raw data (**Fig. 6m, n**). The improved spatial resolution (**Fig. S6e**) also enabled clear views of retrograde flow (**Movie S7**) and fine filopodial dynamics (**Fig. 6o, p**).

## Discussion

Here we synergize concepts from multiview imaging and structured illumination microscopy to create a highly flexible confocal imaging platform that can be tailored to a wide variety of applications and samples (**Fig. 7**). We demonstrate improved performance relative to state-of-the-art implementations of confocal microscopy and spot-scanning SIM^46^, leading to better quality imaging at depth, more nearly isotropic imaging, and improvements in volumetric resolution (**Fig. 1, 2, 4**). Moreover, our work provides a blueprint for the integration of deep learning with fluorescence microscopy (**Fig. 3, 5, 6**). Besides successfully deploying neural networks to denoise our images, enabling lower illumination levels and thus extending imaging duration, we also show such networks can be used to improve imaging at depth (**Fig. 3**). Perhaps most notably, we show that a diffraction-limited line confocal microscope can be used to derive 1D super-resolution training data and that the resulting trained network can, in combination with multiview workflows, produce super-resolution images from diffraction-limited input images (**Fig. 5, 6**). Although acquiring the training data and training the network is time-consuming, applying the trained network is not. Most importantly, our method does not degrade temporal resolution or introduce more dose than required for capturing the base diffraction-limited images. We suspect the same method could also be profitably applied to other microscopes with sharp line-like illumination, including lattice light-sheet microscopy^57^, traditional linear structured illumination microscopy^58^, nonlinear structured illumination microscopy^59^, and stimulated emission depletion microscopy with 1D depletion^60^. The latter two combinations may provide a route to considerably better spatial resolution than we obtained here.

**Fig. 7.**
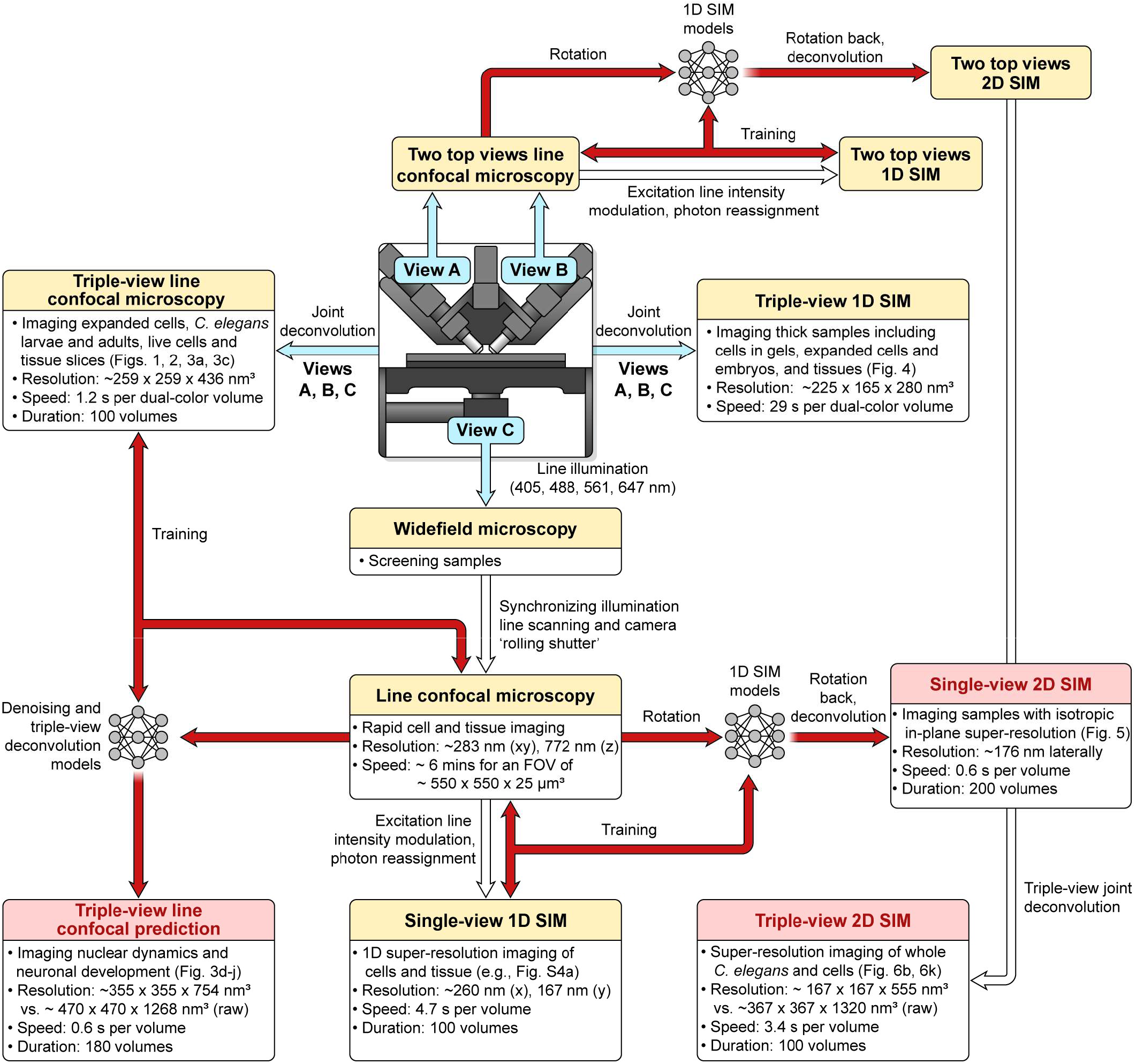
Multi-modality imaging enabled with the multiview line confocal system. Applications include wide-field microscopy, single-view line confocal microscopy (from any of the views), single-view 1D SIM, triple-view diffraction limited imaging, and triple-view 1D SIM. With deep learning (red), triple-view line confocal volumes can be predicted from low SNR single-view input, 1D SIM can be predicted from diffraction-limited input, and combination with joint deconvolution allows further extension to single- and triple-view 2D SIM. Biological and imaging performance examples (resolution, imaging speed and duration) are also provided. Resolution values in line confocal microscopy, triple-view line confocal (without deep learning), and triple-view 1D-SIM (without deep learning) are estimated from immunostained microtubules in fixed U2OS cells. Deep learning resolution values are estimated from fine *C. elegans* embryo neurites (triple-view confocal) or actin fibers (single view 2D SIM, triple-view 2D SIM). See also **Fig. S6f** and **Table S1**.

We envision several extensions and improvements to our work. First, the three objectives we used had relatively long working distances and were chosen to image samples ranging in size from single cells to macroscopic tissue. For studying single cells, higher spatial resolution can be easily attained by using higher NA lenses^61^. Second, our success in imaging expanded tissue suggests that other cleared tissue imaging would also benefit from our methods, provided objective lenses are well matched to the refractive index of the clearing agents being used^21^. Third, although the multi-objective setup used here enables high resolution imaging at depths greater than conventional single-view confocal microscopy, scattering ultimately sets the depth limit. Employing longer wavelength, multiphoton illumination would further improve imaging in thick samples, and could be readily integrated using appropriate fibers^62^. Fourth, although the 50-150 Hz frame rates were well suited to the dynamics studied here, more rapid imaging is conceivable. Five-fold faster 1D SIM imaging is possible using optical reassignment^63^, albeit requiring more emission-side optics, compromising sensitivity. Faster and more light-efficient multiview imaging may be realized by simultaneously recording from all three views (using more complex deconvolution^21,61^ to compensate for the resulting spatially varying PSF), instead of the sequential acquisition scheme employed here. Finally, given that the diffraction-limited line-scanners we developed can be easily modified for lower illumination NA, converting our line-scanners into light-sheets, we anticipate improving spatial resolution in light-sheet microscopy by leveraging cross-modality deep learning techniques^37,38^.

## Author Contributions

Conceived idea: Y.W., H.S. Designed and assembled line-scanners: M.G., J.D. Tested line-scanners: Y.W., X.H., M.G., J.D., H.S. Designed optical setup: Y.W., X.H., H.S. Built optical setup: Y.W., X.H. Designed reconstruction algorithms: Y.W., C.S., P.J.L., H.S. Wrote software: Y.W., X.H., J.L., C.S. Developed deep learning nuclear segmentation pipeline: J.L. Developed cloud and local pipelines for deep learning: Y.W., J.L., J.C., and H.S. Designed experiments: Y.W., X.H. Y.Su, T.S., I. R-S., R.F., A.P., S.R., A.U., D.C-R., H.S. Performed experiments: Y.W., X.H., Y.Su, T.S., I.R-S., R.F., A.P., X.W. Prepared samples: Y.W., X.H., Y.Su, T.S., I.R-S., R.F., A.P., C.C., J.S., R.C. Developed expansion microscopy procedures: Y.Su. Provided reagents or equipment: C. C., X.W., L.B., Y.Sun, L.H.D., Y.P., Y.Shi. All authors analyzed data. Wrote paper: Y.W., X.H., Y.Su, H.S. with input from all authors. Supervised research: Y.W., J.D., Y.P., Y.Shi, E.M., S.R., A.U., D.C-R., P. L.R., H.S. Directed research: H.S.

## Acknowledgements

We thank Dr. Chengyu Liu and Dr. Fan Zhang from the NHLBI Transgenic Core for making the TOMM20-mNeonGreen transgenic mouse line; Erik Jorgensen, Christian Frøkjær-Jensen, and Matthew Rich for sharing the EG6994 *C. elegans* strain; Larry Samelson for the gift of the Jurkat T cells; George Patterson for the H2B-GFP plasmid; John Hammer for the F-tractin-tdTomato plasmid; Noelle Koonce and Lin Shao for sharing images of larvae imaged with spinning disk and iSIM; SVision LLC for maintaining and updating the 3D RCAN Github site; Xuesong Li for assistance with imaging samples on the OMX 3D SIM; Ethan Tyler and Alan Hoofring (NIH Medical Arts) for help with figure preparation; Richard Leapman, Hank Eden, Sapun Parekh, and Min Guo for critical feedback on the manuscript; Qionghai Dai for supporting X.H.’s visit to H.S.’s lab; and Clare Waterman for supporting R.F’s participation in this project. We thank the Research Center for Minority Institutions program, the Marine Biological Laboratories (MBL), and the Instituto de Neurobiología de la Universidad de Puerto Rico for providing meeting and brainstorming platforms. H.S., P.L.R. and D.A.C-R. acknowledge the Whitman and Fellows program at MBL for providing funding and space for discussions valuable to this work. Research in the D.A.C-R. lab was supported by NIH grant No. R24-OD016474, NIH R01NS076558, DP1NS111778 and by an HHMI Scholar Award. X.H. was supported by an international exchange fellowship from the Chinese Scholar Council. This research was supported by the intramural research programs of the National Institute of Biomedical Imaging and Bioengineering; National Institute of Heart, Lung, and Blood; Eunice Kennedy Shriver National Institute of Child Health and Human Development; and the National Cancer Institute within the National Institutes of Health. C.S. acknowledges funding from the National Institute of General Medical Sciences of NIH under Award Number R25GM109439 (Project Title: University of Chicago Initiative for Maximizing Student Development [IMSD]) and NIBIB under grant number T32 EB002103. Y.P. and Y.S. are supported by the Center for Cancer Research, the Intramural Program of the National Cancer Institute, NIH (Z01-BC 006150). This research is funded in part by the Gordon and Betty Moore Foundation. A.U. acknowledges support from NIH R01 GM131054. We thank the Office of Data Science Strategy, NIH, for providing a seed grant enabling us to train deep learning models using cloud-based computational resources.

Disclaimer: The NIH and its staff do not recommend or endorse any company, product or service.

## STAR METHODS

### RESOURCE AVAILABILITY

#### Lead contact

Further information and requests for resources and reagents should be directed to and will be fulfilled by the Lead Contact, Yicong Wu.

#### Materials availability

All cell lines were previously published or available from commercial sources. *C.elegans* strains are available from the Lead Contact.

#### Data and code availability

The data that support the findings of this study are available upon request. The custom code used in this study are available upon request, with most software and test data publicly available at (https://github.com/hroi-aim/multiviewSR, and https://github.com/AiviaCommunity/3D-RCAN).

## EXPERIMENTAL MODEL AND SUBJECT DETAILS

### Cell lines

Mouse melanoma (B16-F10, ATCC CRL-6475), human ovarian carcinoma (HEY-T30, ATCC CRL-3252), human osteosarcoma (U2OS, ATCC HTB-96), human colon carcinoma (HCT-116, ATCC CCL-247), and human T lymphocyte (Jurkat E6-1, ATCC TIB-152) cell lines were used in this study.

### C.elegans

Five nematode strains were used in the study, including EG6994 (unc-119(ed3) III; oxSi469[Pdpy-30::GFP::H2B::tbb-2 cb-unc-119(+)] IV), RW10230 (unc-119(ed3) III; zuIs178 [his-72(1kb 5’ UTR)::his-72::SRPVAT::GFP::his-72(1KB 3’UTR) + 5.7 kb XbaI-HindIII unc-119(+)] V; stIS10024 [pie-1p::H2B::GFP::pie-1 3’ UTR + unc-119(+)]; stIS10224 [lin-39::H1-wCherry + unc-119]), DCR6633 (lim-4p::mCherry::unc54 UTR injected at 25 ng/μL+ unc42p_ZF1_PHD_GFP_unc54 UTR injected at 100ng/μL+ lim-4p::SL2::zif-1::unc54 UTR injected at 25ng/μL+unc-122p::RFP injected at 30 ng/μL), DCR8528 with the genotype syg-1p:PH:GFP injected at 30 ng/μL+ unc-122p:RFP injected at 30 ng/μL in olaIs67 (inx-1-0.5 kb:mCherry injected at 30 ng/μL+inx-1-1 kb:EGFP:Rab-3 injected at 40 ng/μL), and DCR6681 (olaex3993;ujis113 - strain injected with DACR2669 (Pcasy-1A(3.060kb)_PHD_GFP_unc-54 3’utr in pDEST R4-R3) at 100ng/μl + DACR218 (Punc-122_RFP) at 50ng/μl into ujis113 (BV514). Additional information on the Casy-1 promoter is available at promoters.wormguides.org.

### Drosophila

*Drosophila* larval wing discs were imaged this study. Transgenic fly strains were obtained from Bloomington Stock Center: *lexo*-CD2:GFP (BL 66544), *ths*-*GAL4* (BL 77475), and *UAS*-NLS:mCherry (BL 38425). To simultaneously express fluorescently labeled marked in myoblasts and wing disc cells, flies harboring *ths-GAl4/CyO; htl-LexA* were crossed to flies harboring *UAS-*NLS:mCherry*; lexo-*CD2:GFP^33^.

### Mice

Mouse tissue sections (kidney, esophagus, heart and brain) and cardiac myocytes were used in this study. The tissue sections were dissected from 12-week old C57BL male mice generated by the section on Molecular Morphogenesis, Eunice Kennedy Shriver National Institute of Child Health and Human Development (NICHD). The cardiac myocytes were isolated from TOMM20-mNeonGreen expressing male mice aged 15 to 20 weeks from a transgenic line generated by the Transgenic Core, National Heart, Lung and Blood Institute (NHLBI, Dr. Chengyu Liu and Dr. Fan Zhang)^34^. All animal studies were performed in a manner consistent with the recommendations established by the Guide for the Care and Use of Laboratory Animals (National Institutes of Health), and all animal protocols were approved by the Animal Care and Use Committees in NICHD and NHLBI.

## METHOD DETAILS

### Cells

For immunolabeled microtubule samples (**Fig. 1c, Fig. 4b**), U2OS cells were fixed with 4% PFA/PBS and 0.25% glutaraldehyde (Sigma, G6257) for 15 minutes at RT and washed 3 times in 1X PBS. Cells were permeabilized in 0.1% Triton-X 100/PBS for 2 minutes and washed 3 times in 1X PBS. Microtubules were labeled with mouse-α-alpha tubulin (Thermo Fisher Scientific, A11126), α-mouse-biotin (Jackson ImmunoResearch, 715-066-151), and Alexa Fluor 488 Streptavidin (Jackson ImmunoResearch, 016-540-084), each at 1 μg/mL for 1 hour at RT in 1XPBS. In imaging live cells (**Fig. 3a**), HCT-116 human colon carcinoma cells were labeled with 100nM of MitoTracker Green (Thermofisher, M7514) and 5 0nM of LysoTracker Deep Red (Thermofisher, L12492) in imaging media at 37°C for 25 minutes. Cells were washed 3 times in imaging media before being imaged.

For T cell imaging (**Fig. 5f-m**), E6-1 Wild Type Jurkat cells (gift from the Samelson Lab, NIH) were cultured in RPMI 1640 supplemented with 10% fetal bovine serum and 1% Penn-Strep antibiotics. Cells were transiently double transfected with a mixture of mCherry-EMTB and H2B-GFP or EMTB-3XGFP and F-tractin-tdTomato plasmids using the Neon (ThermoFisher Scientific) electroporation system two days before imaging and following the manufacturer’s protocol. Coverslips attached to 8 well Labtek chambers were incubated in Poly-L-Lysine (PLL) at 0.01% W/V (Sigma Aldrich, St. Louis, MO) for 10 min. PLL was aspirated and the slide was left to dry for 1 hour at 37 °C. T cell activating antibody coating was performed by incubating the slides in a 10 μg/ml solution of anti-CD3 antibody (Hit-3a, eBiosciences, San Diego, CA) for 2 hours at 37 °C or overnight at 4 °C. Excess anti-CD3 was removed by washing with L-15 imaging media immediately prior to the experiment.

For live U2OS cell imaging (**Fig. 6k-p**), cells were cultured to 50% confluency on #1.5 cover glass at 37 °C and 5% CO2 in DMEM media supplied with 10% fetal calf serum. Cells were transfected with 100 μL of transfection buffer containing 2 μL of X-tremeGENE™ HP DNA transfection reagent (Sigma, 6366244001), 2 μL of plasmid DNA (tdTomato-Lifeact-7, 500 ng/ μL, Addgene, 54528), and 96 μL of PBS. Cells were imaged 1 day after transfection.

### Expanded cells

For expanded samples with DNA and mitochondrial labels (**Fig. 1e**), U2OS cells were fixed with 4% PFA/PBS and 0.25% glutaraldehyde (Sigma, G6257) for 15 minutes at RT and washed 3 times in 1X PBS. Cells were permeabilized in 0.1% Triton-X 100/PBS for 2 minutes and washed 3 times in 1X PBS. Mitochondria were labeled by rabbit-α-Tom20 (Abcam, ab186735), α-rabbit-biotin (Jackson ImmunoResearch, 711-065-152), and Alexa Fluor 488 Streptavidin, each at 1 μg/mL for 1 hour at RT in 1XPBS. DNA was labeled by 5 μg/mL of mouse-α-DNA (Progen, 61014) and 1 μg/mL of α-mouse-JF549 (Novus, NBP2-60648JF549), each for 1 hour at RT in 1X PBS. For expanded samples with microtubule and mitochondria labels (**Fig. 4g**), the same protocol was applied using mouse-α-alpha tubulin and rabbit-α-Tomm20 as primary antibodies. Stained cells were expanded as described^38^.

### Cells in gels

B16-F10 melanoma cells (ATCC, CRL-6475) were cultured in complete culture media, i.e., DMEM (Thermo Fisher Scientific, 11965-084) with 10% fetal calf serum (R&D Systems, S11150). For collagen (Corning, 354249) gel cultures^47^, cells were trypsinized and resuspended into a 1.6 mg/mL collagen gel in DMEM without serum supplemented with 1:100 dilution of Matrigel (Corning, 354234). The gel was polymerized onto a plasma cleaned coverslip mounted in the imaging chamber at 37C for one hour, and then overlayed with complete culture media. Cells were allowed to migrate in the collagen gel for 16 hours, fixed with 4% paraformaldehyde (Electron Microscopy Sciences, 15710) in cytoskeleton buffer (CB, 20 mM PIPES pH 6.8, 138 mM KCl, 3 mM MgCl2, 2 mM EGTA) and then permeabilized and stained overnight at 4C in CB with 0.5% Triton X-100 (Sigma, T9284) and 2 units/mL Alexa Fluor 488 phalloidin (Thermo Fisher Scientific, A12379). Collagen gel samples were rinsed in PBS and imaged in PBS.

HEY-T30 cells (ATCC CRL-3252) were cultured in RPMI (Thermo Fisher Scientific, 11835-030) with 10% fetal calf serum. For collagen gel cultures, cells were trypsinized and resuspended in 1.6 mg/mL collagen supplemented with 20 mg/mL fibronectin (Millipore, FC010) in DMEM without serum. Gel was polymerized onto a plasma cleaned coverslip for 1 hour at 37°C, then overlaid with complete culture medium and incubated overnight at 37°C. For imaging, media was exchanged with media without serum and supplemented with 300 nM MitoTracker Red CMXRos (Thermo Fisher Scientific, M7512) and incubated at 37°C for 60 minutes. The gel was rinsed with room temperature PBS, fixed in CB buffer with 3% paraformaldehyde, and permeabilized with CB buffer with 0.5% Triton X-100 and 2 units/ml Alexa Fluor 488 Phalloidin overnight at 4C. Labeled cells were stored in PBS until imaging (**Fig. 4c, e**).

### *C. elegans* embryos, larvae, and adults

Nematode strains included EG6994 with nuclei labeled with NucSpot 488 (for the whole L1 larva, **Fig. 1i, j**; and adult, **Fig. 2a-f**), RW10230 with pan-nuclear, GFP-histone label (embryos, **Fig. 3e, f**), DCR6633 expressing GFP-AIB neurons (embryos, **Fig. 3g-j**), DCR6681 expressing GFP-membrane marker (expanded embryo immunostained with rabbit-anti-GFP, anti-rabbit-biotin, and streptavidin Alexa Fluor 488 to amplify GFP signal marking neuronal structures, **Fig. 4i**; expanded embryos stained with DAPI, **Fig. S5b**; and L2 larva, **Fig. S6c**), DCR8528 expressing membrane targeted GFP primarily in the nervous system (a whole worm L2 larva, **Fig. 6b**; L4 larva, **Fig. 6d-j**).

The transgenic strains DCR6633 and DCR8528 were generated via microinjection with the plasmid of interest at 25-100 ng/mL into reference N2 Bristol strains or an integrated AIB marker strain, *olaIs67*, using standard techniques. The *unc-122p: RFP* construct was used as a co-injection marker. We acquired diSPIM and triple-view line confocal volumes of transgenic embryos with one or both AIB neurons labeled^14,39,40^. Cell-specific AIB labeling was achieved by a subtractive labeling strategy^43–45^.

All worms were cultivated at 20°C on NGM plates seeded with OP50 E. coli. Embryos were dissected from gravid adults, placed on poly-L-lysine-coated coverslips and imaged in M9 buffer, as described^35^. When acquiring training data for the triple view line confocal imaging (**Fig. 3e-j**), sodium azide was added to the M9 buffer with a final concentration of 0.3% to slow embryo twitching. For expanding embryos, see more details in the section *C. elegans* embryo expansion. For imaging worm larval and adults, worms were put in imaging chambers, ‘hot-fixed’ in 4% PFA in PBS at 50°C for 5 minutes, and then washed three times using PBS before imaging.

### *C. elegans* embryo expansion

*C. elegans* embryos (strain DCR6681) were immobilized on Poly-L-Lysine (PLL) coated glass bottom dishes, permeabilized with bleach and Yatalase, fixed, immunolabeled, and expanded. The procedure takes approximately four days and is described in detail below (see also **Fig. S5a**).

First, Poly-L-Lysine (PLL) coating glass bottom dishes were prepared. Poly-L-Lysine (Sigma, P7405) powder was reconstituted in distilled water to 1mg/mL, aliquoted, and stored at −20°C. Prior to experiments, 30-50 μL of Poly-L-Lysine was placed on the glass bottom dish (MatTek, P35G-1.5-14-C) and air dried at room temperature (RT). Coated coverslips were usually prepared up to 1 day before pre-treatment of C. elegans for expansion microscopy.

Second, C. elegans embryos were digested and fixed. Gravid adult C. elegans worms were deposited in a petri dish in PBS buffer and cut with a surgical blade to release eggs. Eggs were immobilized on a PLL coated glass bottom dish in PBS. Eggs could be processed immediately or stored at 25°C in M9 buffer until the embryos developed to the desired stage. Embryos were treated by 1% hypochloric acid (Sigma, 425044) for 2-3 minutes in 0.1M NaOH/water, rinsed 3 times in PBS, digested in 50 mg/mL Yatalase in PBS (Takara Bio, T017) for 40 minutes at RT and rinsed 3 times with PBS. We found that using yatalase proved more effective than chitinase in dissolving the *C. elegans* eggshell, a key factor in achieving isotropic expansion. It was important to treat eggs with bleach only after immobilization on the PLL surface, otherwise embryos tend to detach from the glass at later steps. Digested embryos were fixed in 4% paraformaldehyde/PBS (Electron Microscopy Sciences, RT 15710) for 1 hour, then rinsed 3 times with PBS and permeabilized with staining buffer (0.1% Triton X-100/PBS, Sigma, 93443) for 1 hour before immunolabeling.

Third, *C. elegans* embryos were immunolabeled. *C. elegans* embryos were stained by an anti-GFP primary antibody (Abcam, ab290) in the staining buffer at 4°C overnight. After primary antibody labeling, embryos were washed 3 times (30 min intervals between washes) in the staining buffer and labeled using donkey-anti-rabbit-biotin secondary antibody (Jackson ImmunoResearch, 711-065-152) in the staining buffer at 4°C overnight. After secondary antibody labeling, the embryos were washed 3 times in the staining buffer (30 mins intervals between washes) and labeled with Alexa Fluor 488 Streptavidin (Jackson ImmunoResearch, 016-540-084). Labeled embryos were washed 3 times in the staining buffer (30 minutes between washes) before being processed for expansion microscopy.

Last, immunolabeled *C. elegans* embryos were expanded. Immunolabeled *C. elegans* embryos were treated with 1mM MA-NHS (Sigma, 730300) in PBS for 1 hour at RT. Samples were rinsed 3 times in PBS, and treated with monomer solution, which was made up of acrylamide (Sigma, A9099), sodium acrylate (Santa Cruz, 7446-81-3), N, N’-methylenbisacrylamide (Sigma, 146072) and 4-Hydroxy-TEMPO (Sigma, 176414), diluted with PBS, with a final concentration of 10%, 19%, 0.1%, and 0.01%, respectively. After the treatment for 1 hour at RT, the monomer solution was replaced by gelation solution. The gelation solution had the same ingredients and concentration with monomer solution, with the additions of tetramethylethylenediamine (TEMED, ThermoFisher Scientific, 17919, reaching a final concentration of 0.2%) and ammonium persulfate (APS, ThermoFisher Scientific, 17874, reaching a final concentration of 0.2%). APS was added at last, and the fresh gelation solution was immediately applied to the embryos sandwiched between the glass bottom dish and another coverslip surface for 2 hours at RT. It was important to control the gelation speed with 4-hydroxy-TEMPO (Sigma, 176414) as premature gelation can distort embryos and result in poor expansion quality. The space between the glass bottom and the coverslip was controlled by a pair of 51 μm spacers (Precision Brand, 44910). The polymerized embryo-hydrogel hybrid was cut out by a razor blade and digested with 0.2 mg/mL proteinase K (Thermofisher Scientific, AM2548) in the digestion buffer (sodium Chloride (Quality Biological, 351-036-101) with 0.5 M; guanidine hydrochloride (Sigma, G9284) with 0.8M; and Triton X-100 (Sigma, 93443) with 0.5% concentration) at 45°C overnight. Digested embryos were expanded ^~^3.3-fold in distilled water (Fig. S5b), and fresh distilled water was exchanged every 30 min until the samples were fully expanded. Expanded samples were mounted on PLL coated #1.5 coverslips (VWR, 48393-241) and secured in an imaging chamber. Finally, the chamber was filled with distilled water before commencing imaging (Fig. 4i).

### Drosophila samples

All flies were raised at 25 °C with 12 h/12 h light/dark cycle. Third instar larva were dissected in 1XPBS (Hardy Diagnostics) and fixed in 4% PFA in PBS for 25 minutes. The larvae were then washed three times for 5 minutes each using 0.1% PBST (0.1% TritonX-114 in PBS). The wing discs were mounted in 1XPBS in an imaging chamber prior to imaging (**Fig. 2g, Fig. S2c-d**).

### Tissue section preparation and immunolabeling

Organs from C57BL mice were fixed in 4% PFA/PBS at 4°C overnight. Fixed tissue was incubated in a serial dilution (10%, 20%, 30%) of sucrose/water solution (Sigma, S0389) for 3 hours each at RT, followed by equilibration in 30% sucrose at 4°C overnight. Tissue was cut into cubes using a razor blade and submerged in Optimal Cutting Temperature solution (O.C.T, Fischer Scientific NC9806257) in a mold (Fischer Scientific, 22-363-552), and frozen in pure ethanol/dry ice solution for 10 minutes. Frozen tissue was sectioned to various thicknesses using a Leica CM1850 cryostat, then stored in 1X PBS.

Esophagus and kidney tissue sections were incubated in antigen retrieval buffer (Abcam, ab93684) at 60°C for 1 hour. Esophagus tissue sections were incubated with 1μg/mL of mouse-α-Tubulin primary antibody in 0.1% Triton-X/PBS at 4°C overnight, washed in 0.1% Triton-X/PBS for 1 hour at RT, and incubated with 1μg/mL of α-mouse-JF549 and 10μM of phalloidin-Alexa Fluor 488 (Thermo Fisher Scientific, A12379) in 0.1% Triton-X/PBS for 1 hour at RT. Kidney tissue sections were incubated with 1μg/mL of mouse-α-Tubulin primary antibody and Goat-α-CD31 primary antibody (R&D Systems, AF3628) in 0.1% Triton-X/PBS at 4°C overnight, washed in 0.1% Triton-X/PBS for 1 hour at RT, and incubated with 1μg/mL of α-mouse-JF549, α-goat-AF647 (Jackson ImmunoResearch, 705-605-147), 10μM of phalloidin-Alexa Fluor 488, and 1μg/mL of DAPI (Thermo Fisher Scientific, D1306) in 0.1% Triton-X/PBS at 4°C overnight. Heart tissue sections were incubated in 0.1% Triton X-100/PBS with 1μM of ATTO 647N-NHS ester (Sigma, 18373) and 1X NucSpot Live 488 (Biotium, 40081) at RT for 30 minutes. Stained tissue sections were washed in 0.1% Triton-X/PBS at RT for 1 hour and mounted on a poly-L-lysine coated coverglass (VWR, 48393-241) for imaging (**Fig. 2m, Fig. 4l, Fig. S2e-g**).

### Cardiomyocytes

TOMM20-mNeonGreen expressing male mice were anaesthetized with an intraperitoneal injection of sodium pentobarbital (50 mg/kg). Heparin (0.1 ml of 1000 U/ml) was co-administered to prevent blood clotting during the isolation procedure. The heart was excised from the animal and retrogradely perfused for 6-8 minutes with Ca^2+^ free Tyrode’s solution (mM, pH7.4): 140 NaCl, 4 KCl, 1 MgCl_2_, 5 HEPES, 10 D-glucose (equilibrated with 100% O_2_ at 37°C). Subsequently, hearts were enzymatically digested by perfusing with the same solution plus 100 μM Ca^2+^ and Liberase TM (Roche, final 0.25 collagenase Wunsch units/ml) for 6-8 minutes. Following perfusion digestion, the heart is removed from the cannula and the left ventricle is cut into small pieces in the modified Liberase Tyrode solution. Tissue was gently agitated to dissociate individual myocytes. The digestion was stopped by adding 5 mg/ml BSA and allowed to settle for another 20 min at room temperature, during which time Ca^2+^ is added incrementally back up to 1.2 mM. Subsequently myocytes were separated by centrifugation (150 x g for 1 min) at room temperature. Pelleted myocytes were resuspended in Tyrod’s solution containing 1.2 mM CaCl_2_. For imaging, a small aliquot of suspended cells was transferred to a coverslip or a 35 mm glass (no. 1.5) bottom dish (MatTek Life Sciences, P35G-1.5-14-C) and incubated with 100 nM of MitoTracker Green (ThermoFisher Scientific, M7514) in imaging media (ThermoFisher Scientific, A14291DJ). Cells were rinsed 3 times in imaging media and imaged at room temperature.

### Triple-view microscope frame

The microscope frame and objective configuration (**Fig. S1a**) is similar to that used for triple-view wide-field and light sheet microscopy^20^. A modular microscope frame with a motorized XY stage (Applied Scientific Instrumentation, RAMM and S551-2230A) served as the base, and an XY piezoelectric stage (Physik Instrumente, P-545.2C7, 200 μm × 200 μm lateral travel) was bolted on the top of the stage to provide precise translation of the sample. Rectangular coverslips containing samples were placed in an imaging chamber (Applied Scientific Instrumentation, I-2450), the chamber was placed into a stage insert (Applied Scientific Instrumentation, PI545.2C7-3078), and the insert was mounted to the piezo stage.

Three line-scanning confocal modules were coupled to the microscope base, each containing a microelectromechanical systems (MEMS)-based line scanner (**Fig. S1b**, Applied Scientific Instrumentation, Scan-SL, MM-CYL-SL-2.4-FA20), an excitation tube lens (Applied Scientific Instrumentation, C60-TUBE-160 with C60-NRADJ-C-MOUNT), tip/tilt adjustable holders for mirrors and filters (Applied Scientific Instrumentation, C60-CUBE-III and C60-D-CUBE), an objective lens housed within a piezoelectric stage (Applied Scientific Instrumentation, APZOBJ-150), an emission tube lens (Applied Scientific Instrumentation, C60-TUBE-B) and an sCMOS camera (PCO, Edge 5.5). Two modules were equipped with 40x, 0.8 NA water-immersion objectives (OBJ A and OBJ B in **Fig. 1a** and **Fig. S1a**, Nikon Cat. # MRD07420) and accompanying piezoelectric stages (Applied Scientific Instrumentation, APZOBJ-150, used for fine alignment and for acquiring volumes with OBJ A/B) and held in a perpendicular configuration above the imaging chamber. These modules were mounted on a motorized z translation stage (Applied Scientific Instrumentation, LS-50-AMCLH), so their axial position can be adjusted relative to the sample chamber. The third module contained a 60x, 1.2 NA water objective (OBJ C in **Fig. 1a** and **Fig. S1a**, Olympus UPLSAPO60XWPSF) and accompanying piezoelectric stage (Physik Instrumente, PIFOC-P726), placed directly below the imaging chamber. The third module is also mounted on a z translation stage (Applied Scientific Instrumentation, LS-50-AMERL).

The motorized XY stage was used for (1) coarse sample positioning before imaging; (2) accessing multiple sample positions (e.g., imaging multiple embryos in time lapse imaging); (3) extending the field of view by translating samples. Objective piezo stages were used to create imaging volumes for all three views. For the two top views, imaging volumes can be also created by stepping the sample through the stationary confocal plane with the XY piezo stage. More detail is provided in the sections *Triple-view data acquisition* and *Triple-view data processing*.

### Triple-view line confocal excitation and detection optics

Excitation and detection optics (**Fig. S1c**) for triple-view line confocal microscopy are similar to those used in fiber-coupled diSPIM^23,39^, except that sharply focused line illumination is employed to collect fluorescence in epi-mode. 405 nm (Spectra-Physics, Excelsior 4050-200-CDRH), 488 nm (Newport, PC14584), 561 nm (Coherent, Sapphire 561-300 CW) and 647 nm (Coherent, Cube 647-100C, 1190357) laser diode outputs were combined with dichroic mirrors (Semrock, Di02-R405-25×36, Di02-R488-25×36, Di02-R561-25×36) and passed through an acousto-optic tunable filter (AOTF, Quanta Tech, AOTFnC-400.650-TN) for power and shutter control. Two beam splitters, 70:30 ratio (Thorlabs, BS058), and 50:50 ratio (Thorlabs, BS010), were used to obtain three beams with similar power. Each beam was directed into its own single-mode fiber via a galvanometric mirror (Thorlabs, GVSM001). The galvanometric mirrors served as shutters, switching the excitation on in series for sequential triple-view imaging. They were also used to finely tune the power of each arm by changing the fiber coupling efficiency. Illumination power varied between 0.2 – 0.6 mW in this work, measured after the objective.

Each fiber (Qioptiq, kineFLEX-P-2-S-488.640-0.7-APC-P2) is coupled into a MEMS line scanner (**Fig. S1b**, Applied Scientific Instrumentation, MM-CYL-SL-2.4-FA20). The optics within the scanner collimate the fiber-coupled beam, pass it through an aperture stop, focus it with a cylindrical lens to form line illumination, scan it with a MEMS mirror (2.4 mm diameter), and image it to a field stop at a conjugate sample plane via lens pairs L1 and L2 (**Fig. S1c**). The MEMS mirror scans in two dimensions, one for vertically scanning the line to create an imaging plane, the other for adjusting the lateral position of the excitation line. Two aperture sliders optionally allow adjustment of the lateral and vertical input beam size (affecting the NA of the illumination) or cropping of the illumination along the line’s long axis (avoiding unnecessary excitation).

After the line scanner, the scanned beam was relayed via an excitation tube lens (L3, focal length FL = 160 mm, Applied Scientific Instrumentation, C60-TUBE-160 with C60-NRADJ-C-MOUNT) and the objective (OBJ) to the sample plane, after reflection from a dichroic mirror (Chroma, ZT405/488/561/647rpc). This optical magnification guaranteed that the NA of the objective could be fully used: once the MEMS mirror was filled after fiber collimation, the maximum beam diameter at the back focal plane of the objective is *D* x cos(22.5°) x FL_L3_/ FL_L2_ = ^~^2.22 mm x 160 mm/39 mm = ^~^9.1 mm, as the MEMS mirror has diameter (*D*) 2.4 mm and is tilted at 22.5° relative to the main axis of L2 and L3. This beam diameter overfills the back apertures of the OBJ A/B and OBJ C, which are given by 2 x NA x FL = 2 x 0.8 x 200/40 mm = 8 mm and 2 x 1.2 x 180/60 mm = 7.2 mm, respectively, indicating that diffraction-limited excitation is expected barring additional aberrations.

Fluorescence was collected in epi-mode, transmitted through the dichroic mirror, reflected by a mirror (Applied Scientific Instrumentation, C60-30mm-CUBE-RA-MIRROR) mounted within a kinematic adjuster (Applied Scientific Instrumentation, C60-CUBE-III), further filtered by a notch emission filter (Chroma, ZET405/488/561/647m) to reject excitation light, and imaged via an emission tube lens (L4, f = 200 mm, Applied Scientific Instrumentation, C60-TUBE-B) onto a scientific CMOS camera (PCO, Edge 5.5). Lens pair L1, L2 were placed in 4f imaging configuration, as were lens pair L3, OBJ. Lenses L1, L2, L3, and L4 were the same in all three views. The resulting image pixel sizes for the top views were 162.5 nm, and for the bottom view, 97.5 nm. Cameras were operated in ‘light-sheet mode’^20,23^, synchronizing the scanned beam with the rolling shutter readout to achieve confocality (**Fig. S1d**, see also *Triple-view data acquisition*). These optical configurations allowed us to achieve diffraction-limited confocal imaging with a field of view of ^~^ 260 μm x 260 μm in the top views and ^~^175 μm x 175 μm in the bottom view (**Fig. S1e**).

### Triple-view system alignment and calibration

Alignment is critical for optimal triple-view imaging. First, to achieve the sharpest excitation on the sample plane, the focused, scanned beam produced by the line scanner must be properly relayed to the sample plane. The thickness of the excitation line at the sample was first visually checked by imaging a dye solution (e.g., fluorescein), coarsely adjusting the distance between the fiber tip and the collimating lens via the rotation of an eccentric screw focusing tool on the collimating lens’s cartridge to translate the axial position of the beam waist (i.e., to narrow the width of the line as viewed in the camera). Next, we finely tuned the distance between the line scanner and the excitation tube lens with the non-rotating adjustable C-mount (Applied Scientific Instrumentation, C60-NRADJ-C-MOUNT), first minimizing the apparent line width in dye and second minimizing a more accurate estimate of the illumination beam waist at various points in the field. The latter was obtained by scanning the line illumination over stationary 100 nm fluorescent beads (Thermo Fisher Scientific, F8803) sparsely deposited on a coverslip, acquiring an image at each position of the line focus, and inspecting the intensity as a function of scan position (i.e., estimating the width of the illumination PSF). After optimization, this method yielded 361 ± 22 nm FWHM for the top two views and 267 ± 23 nm for the bottom view (**Fig. S1e**, *N* = 10 measurements across the 260 x 260 μm^2^ top view field, and 175 x 175 μm^2^ bottom view field).

Second, to achieve optimal confocality, it is important to minimize both the curvature of the excitation line (here defined as the offset between the midpoint of the line and the endpoints, ideally zero) and its rotation angle when scanned, ensuring that the scanned lines are as parallel as possible to each other. We achieved this by again using fluorescent dye solution, iteratively steering the excitation beam entering the objective by tilting the dichroic mirror via kinematic knobs on its holder (Applied Scientific Instrumentation, C60-CUBE-III), while also steering the emission beam by rotating the relevant mirror (located after the dichroic mirror) via another set of knobs on its mount (Applied Scientific Instrumentation, C60-CUBE-III). The maximum curvature and rotation angles of the excitation line were measured to be less than 0.09 μm and 2×10^−5^ radian/mm, respectively, over 80% of the image field for Views A/B, and 0.19 μm and 8×10^−5^ radian/mm, over 80% of the image field for View C. Global rotation of the scanned lines, ensuring that they are perpendicular to the direction of camera readout, was achieved by rotating the camera with a rotating C-mount adapter (Applied Scientific Instrumentation, C60-3060-CMR).

Last, we encountered two challenges in synchronizing the MEMS mirror with the rolling shutter:

(1) a ^~^2 ms delay between the driving waveform and the MEMS actuator; (2) a non-linear relationship between the driving voltage and the MEMS position. The first issue is easily dealt with by padding the imaging region extra pixel rows (typically ^~^100); these rows are then discarded from the final image. To address the second issue, the voltage-position response must be calibrated for each view to guarantee synchronization of the camera’ rolling shutter and the excitation line scan. We obtained the calibration curves (i.e., the MEMS driving waveforms) by (1) acquiring a series of images of a dye solution, similar to **Fig. S1e**, but with fine voltage steps every 5 mV from 0 to 5 V; (2) localizing the vertical positions of each excitation line on the camera chip; (3) interpolating this nonlinear response curve so that each pixel row on the camera chip corresponds to a MEMS driving voltage. This calibration curve is quite stable, and only needs to be redone if the optics are re-aligned.

### Triple-view data acquisition

Control software with a graphical user interface (GUI) was written in MATLAB (Mathworks, R2019b, and with Image Processing Toolbox), running on a HP Z820 workstation (Intel Xeon CPU E5-2630 v2 @ 2.60 GHz, 24 threads, 64 GB memory, and 1 TB SCSI hard drive disk). Waveforms for driving the MEMS mirror, stage and objective piezos, and AOTF were produced with three data acquisition (DAQ) cards (National Instruments, PCI 6733, BNC 2110) installed in a DAQ card chassis (National Instruments, PXI-e-1073). Dynamic-link library (DLL) files provided by National Instruments and PCO were called in MATLAB to control the DAQ cards and cameras. We also used our software to control the non-piezo XY translational stage and Z translation stages via a controller hub (Applied Scientific Instrumentation, TG16) by sending or receiving serial commands (e.g., moving the stage to a defined location, or reading the current position).

The three views were acquired sequentially in the order A->B->C. **Fig. S1d** illustrates the control waveforms for one view, when acquiring diffraction-limited data. A pulse train was used to trigger the camera, and the camera output then used to trigger all driving waveforms. We noticed that the timing between the camera trigger input and the actual exposure start was unpredictable (varying from 0 to 100 μs); choosing the camera output as the master signal avoids this jitter.

Once the camera output trigger was received, a waveform was generated to drive the MEMS mirror, thereby scanning the excitation line in the sample during the camera exposure time. The duration of the excitation was controlled by the AOTF waveform. The camera was run in rolling shutter mode with a tunable slit width (colloquially known as ‘light-sheet readout mode’) so that the camera exposure was synchronized with the scanning line excitation. The slit width was set to 3 - 6 pixels (i.e., ^~^ 0.3 - 0.6 μm for View C, or ^~^0.5 - 1 μm for Views A/B) for balancing the rejection of out of focus light and signal intensity. Example images acquired in *C. elegans* embryos expressing GFP-histones with and without the rolling slit are shown in **Fig. S1f**.

Analog step-wise waveforms were used to drive the piezoelectric objective stages for all three views (or the piezo XY stage only for Views A/B) for volumetric acquisition. When imaging small samples (e.g., *C. elegans* embryos and cells), we used ‘objective piezo scan’ mode, i.e., triple-view volumes were acquired by moving the objectives via their own piezo stages. In this case, there was no pause introduced when switching views. When imaging large and flat samples (e.g., worm adults and tissue), we employed ‘stage scan’ mode so that the top two view volumes were acquired by laterally moving the sample via the XY piezo stage. In this case, after acquiring Views A/B in the ‘stage scan’ mode, the XY piezo stage returned to its original position and paused for 200 ms to settle before View C was acquired in ‘objective piezo scan’ mode. For accessing multiple sample positions or extending the field of view by translating samples, the required positions of the motorized MS-2000 translation stage were first compiled and saved in the MATLAB program; then after each triple-view acquisition, the translational stage was directed to move to the next position. For multi-color imaging, the piezo stages moved after all the colors were acquired, i.e., we first acquired all colors per plane and then moved to acquire the next plane.

For triple-view 1D SIM imaging, five consecutive images with illumination patterns phased ^~^2pi/5 apart were acquired for super-resolution reconstruction (see section *Triple-view data processing*). At each phase, the excitation was turned on for 1 row of pixels, off for 4 rows of pixels, and this duty cycle repeated over the camera chip. We achieved this on/off modulation by using the AOTF to rapidly modulate the laser illumination during each camera exposure. For the next phase, the ‘on’ rows were shifted by one row relative to the previous row. We explored alternate on/off duty cycles (**Fig. S4c**), but settled on the five-phase pattern as it gave good modulation contrast (necessary for noticeable resolution enhancement) while still requiring relatively few raw images.

Volumetric acquisition time was determined by the camera line exposure time, number of rows of pixels in the image, camera readout time, and the number of z slices. For example, in the live cell imaging shown in **Fig. 3a**, the line exposure time was set to 20 μs, yielding a frame exposure time of 6.4 ms for Views A/B and a 9 ms frame exposure time for View C; when setting a 1 μm z step size for Views A/B, and 0. 5 μm z step size for View C, we were able to collect each dual-color, triple-view diffraction-limited volume spanning 32 μm x 32 μm x 16 μm in ^~^1.2 s. **Table S1** details the volume size and corresponding acquisition time for all the data presented in the paper.

### Triple-view data processing

Raw image data from the three views were merged to produce a single volumetric view, after processing steps that include background subtraction, digital photon reassignment of 5 phase images in 1D SIM mode (more detail is provided in the section *1D SIM data reconstruction*), interpolation, transformation, registration, and deconvolution. The data processing software was written in MATLAB (Mathworks, R2019b, with Image Processing and Parallel Computing Toolboxes), running on a HP Z840 workstation (CPU: Intel Xeon CPU E5-2660 v3 @ 2.60 GHz, 48 threads, RAM: 256 GB; GPU: NVIDIA GPU TITAN RTX, 24 GB memory). Background subtraction, interpolation, and transformation procedures that transform the data to the perspective of the coverslip (i.e., the bottom view’s perspective), either in ‘objective piezo scan’ mode or ‘stage scan’ mode, are similar to those previously described^20^.

After producing three views with the same perspective and same pixel size as view C (i.e., 97.5 nm in diffraction-limited mode, 48.75 nm in SIM mode), a Fourier-based phase correlation method (implemented in MATLAB with GPU processing) was used to coarsely register (translation only) the two top views to the bottom view, followed by fine registration using GPU-based 3D affine registration (initially written in C++/CUDA, called the DLL file in MATLAB)^21^.

As with previous dual-view^14^ and triple-view^20^ light sheet deconvolution, the algorithm we developed to fuse triple-view line confocal datasets is based on Richardson-Lucy deconvolution (RLD)^24,25^, implemented in the Fourier domain using GPU processing in MATLAB. We programmed both alternating RLD updates (Equations 1) and additive RLD updates (Equations 2) for processing different samples with 15-30 iterations (**Table S1**). For dim samples, the additive method usually yields better reconstructions than the alternating joint deconvolution updates, but the latter converges faster than the former^20^. For thick, scattering samples (**Fig. 2b, Fig. S2a**), fusion of all three views with the additive method provides complementary information from each view, but signal in the sample interior still appears dim relative to the peripheral regions (**Fig. 2d**). To remedy this issue, we explicitly incorporate decay functions in the additive RLD updates (Equations 3) by modifying the convolution term in the forward model (i.e., the convolution of the sample estimate and PSF). We derived the decay functions from the data by fitting exponential functions to the average depth-dependent attenuation in each view (**Fig. S2b**). This new deconvolution method enables much clearer reconstructions throughout the sample volume (**Fig. 2e**). To compute the convolutions in Equations 1-3, we used simulated PSFs (both in diffraction-limited mode and 1D SIM mode, see the section *PSF Modelling*).

Alternating RLD:

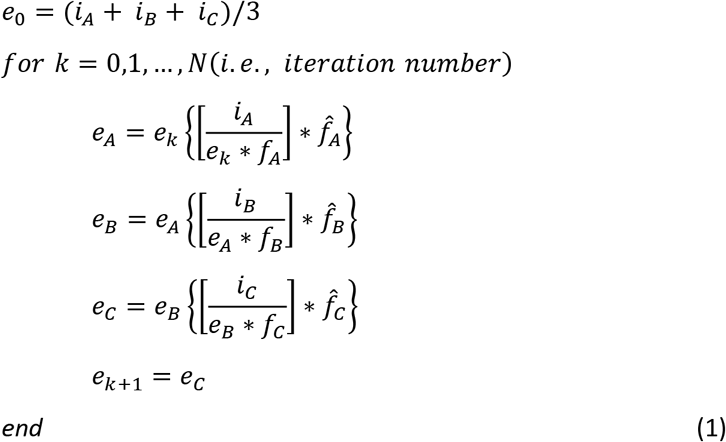

where *i*_*A*_, *e*_*A*_, *f*_*A*_, 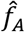, *i*_*B*_, *e*_*B*_, *f*_*B*_, 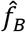, *i*_*C*_, *e*_*C*_, *f*_*C*_, 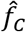 are the raw images, estimates, point spread functions (PSFs), and adjoint PSFs, corresponding to views A, B and C respectively; and * denotes convolution.

Additive RLD:

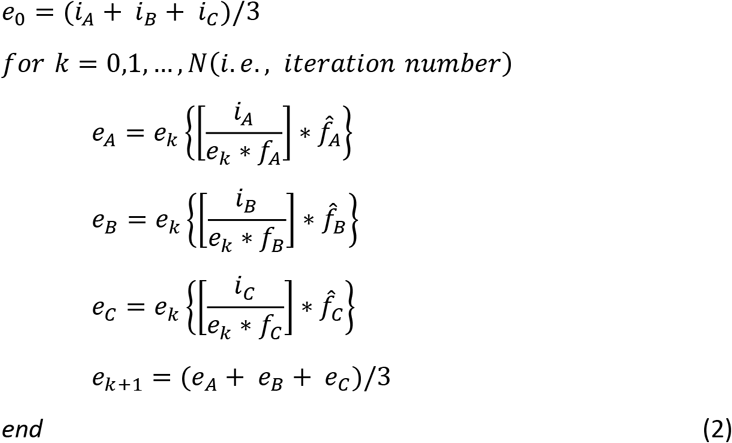

where *i, e, f,* 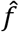 are defined as above and the subscripts *A, B, C* indicating each view.

Additive RLD with attenuation correction:

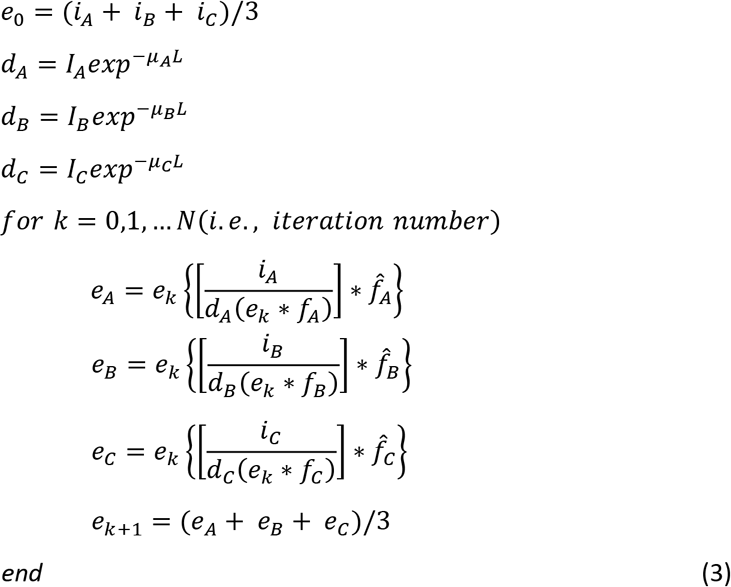

where *i, e, f,* 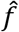 are defined as above and the subscripts *A, B, C* indicate each view. Here *d*_*A*_, *d*_*B*_, *d*_*C*_ are the exponential attenuations (with attenuation coefficient *μ*_*A*_, *μ*_*B*_, *μ*_*C*_) as a function of depth (*L*) for each raw view, estimated from the data; *I*_*A*_, *I*_*B*_, *I*_*C*_ are the intensity at the sample surface for each view.

As the GPU-based triple-view deconvolution requires ^~^20 3D variables and 6 convolutions, the maximum size of the deconvolved volume that can be processed with the GPU in our workstation (24 GB memory) is only ^~^0.3 GB. For datasets that exceed this limit (e.g., the expanded cell sample with 1400 x 1400 x 530 pixels shown in **Fig. 1e**, also see **Table S1**), we adopted the postprocessing previously developed for registering and jointly deconvolving large cleared-tissue diSPIM datasets^21^. We cropped each single-view volume into overlapping subvolumes, registered and deconvolved the subvolumes, removed the boundary regions of the deconvolved subvolumes (40 pixels from each edge, in all three dimensions) to eliminate edge artifacts, and finally stitched the resulting reconstruction back into the original size. Linear blending was performed on the overlapped regions of the adjacent subvolumes (40 pixels from each edge, in all three dimension) to lessen stitching artifacts. This processing pipeline currently allows the handling of triple-view datasets up to hundreds of GB scale (e.g., 3 views, 2 colors, 21 tiles, 1200 x 1200 x 500 pixels per tile, in the esophageal tissue images in SIM mode, **Fig. 4l**, see **Table S1**).

### 1D SIM data reconstruction

We processed sets of images excited with sparse, periodic, phase-shifted line illumination (3-6 phases when choosing the number of phases for reconstruction, **Fig. S4a-h**; 5 phases in all other SIM data presented in this paper, **Figs. 4-6**) into super-resolution images using digital photon reassignment^8,9^. The processing steps resemble those derived for multifocal SIM^9^ but are simpler as only 1D reassignment is needed here. The reconstruction code was written in MATLAB (Mathworks, R2019b, using the Image Processing Toolbox).

The first key step is to precisely estimate the illumination patterns, obtaining the spatial positions of all intensity maxima with subpixel precision. In each image, we averaged the intensity of pixels in each row to create a 1D intensity profile across the scan direction, then digitally upsampled the scan dimension by 10-fold to obtain a more finely sampled profile. The illumination offsets *ϕ*_*n*_ are initially estimated by finding the phase shift that maximizes the cross-correlation between the 1D intensity profile and a cosine function with period 2π/*N*, where *N* is the number of phases, i.e.

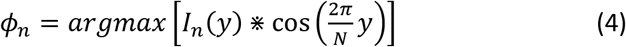

Here *n* = 1, 2, …., *N*; *I*_*n*_ is the 1D intensity profile of the *n*th phase image, and ⋇ denotes cross correlation.

We repeated this process for all images, obtaining a set of offsets *ϕ*_1_ *… ϕ*_*N*_. To minimize the effects of noise, we then aligned all phases to the first by subtracting the expected phase difference between images (successive illumination patterns are expected to be shifted by exactly one camera row, or 2π/*N* phase difference), and averaged them to obtain an improved estimate of the first phase:

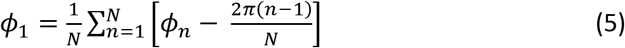

We then shifted each phase back to its expected position, now based on the improved estimate of the first phase:

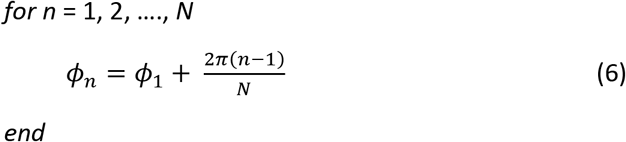

When imaging densely labeled samples, the periphery of the volume was sometimes dominated by out of focus background, and the above method was unreliable on the associated image planes. In this case, rather than running the above method on each image plane, we instead ran it on the XY maximum intensity projection (MIP) of the whole volume. Since the illumination pattern for a given phase shift is expected to be the same at each plane, using the MIP-derived offsets allowed reasonable pattern estimation.

After deriving the positions of the illumination maxima, the second key step is to apply digital photon reassignment in the vicinity of each maximum, thus improving the resolution by ^~^1.4x along the scanning direction. We extracted sub-images (7 rows, approximately twice the width of the system PSF) centered at each maximum and re-assigned the signal to a new location such that the distance between reassigned maxima was twice the distance in the raw data. This procedure shrinks the effective pixel size along the line scan direction by a factor of two (e.g., 49.75 nm for View C, instead of the raw value of 99.5 nm). Next, we multiplied the sub-images by a 1D Gaussian window and summed them to reconstruct a super-resolution image. The window (kernel size = 7 pixels, sigma = 1.3 pixels) served as a digital pinhole to further reject out-of-focus light in the reconstruction, also helping to smooth the overlap regions between the adjacent sub-images. Finally, we digitally upsampled the non-scan dimension so that the 1D super-resolution image had isotropic pixels. The processing time for this procedure scales with the volume size, e.g., ^~^ 15 s for reconstructing a 1D SIM volume with a final size of 1280 x 1280 x 480 pixels). Single-view RLD deconvolution (with 10 iterations) with a 1D SIM PSF (see *PSF Modeling*) was performed on the reconstructed 1D SIM images to further boost signals at high spatial frequencies (e.g., **Fig. S4a-b**). For some comparisons, we summed the raw phase images to obtain an average image (diffraction-limited), interpolating the result to the same final pixel size as the super resolution image, and performed single-view deconvolution if needed (e.g., **Fig. 4a**).

#### View C and triple-view 2D SIM processing

After our RCAN predicted 1D super resolved images at six rotations per view, joint RLD deconvolution (with 10-20 iterations, Equations 7) was performed to achieve isotropic in-plane super-resolution images for View C (**Fig. 5**) or triple-view 2D SIM (**Fig. 6**). Code was written in MATLAB (Mathworks, R2019b, using the Image Processing and Parallel Computing Toolboxes).

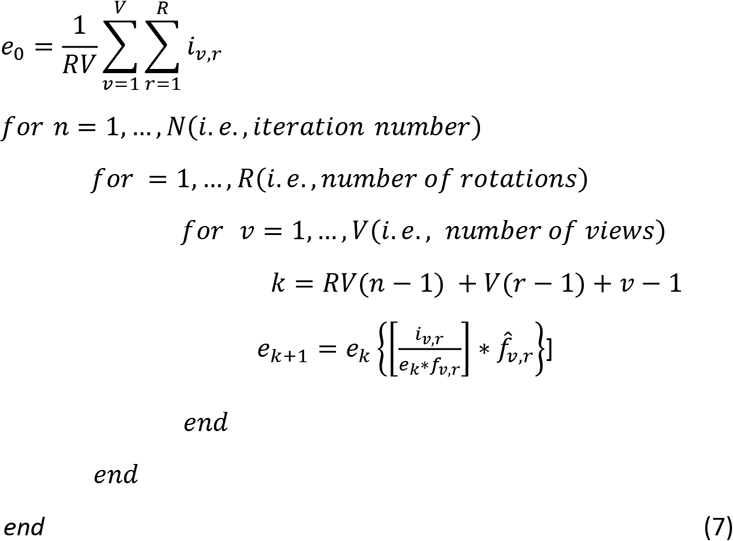

where *i*_*v,r*_, *f*_*v,r*_,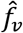, respectively, the 1D super-resolved images, point spread functions (PSFs), and adjoint PSFs, corresponding to the view ν (i.e., view C only when *V* = 1, or views A/B/C when *V* = 3) and rotation *r* (i.e., 0°, 30°, 60°, 90°, 120°, and 150° when *R* = 6, or 0° and 90° when *R* = 2). *e*_*k*_ is the estimate at *k* = *RV*(*n* − 1) + *V*(*r* − 1) + *ν* − 1; and * denotes convolution.

The joint deconvolution of multiple volumes (e.g., 18 volumes for triple-view 2D SIM) requires ^~^100 3D variables that could not all be loaded into the GPU at once due to memory constraints, thus we only loaded the variables required at each iteration 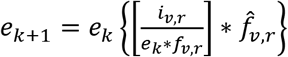. We also adopted the above-mentioned postprocessing pipeline (i.e., cropping the views/rotations into subvolumes, deconvolving the subvolumes, and stitching the results, see the section *Triple-view data processing*) to scale up the data processing ability (this procedure allowed us to scale up the processing to larger volumes, taking e.g. ^~^ 45 min for reconstructing a triple-view 2D SIM volume with a final size of 1560 x 1720 x 164 voxels, including file reading/writing, **Fig. 6k**).

### PSF Modeling

3D system PSFs (Equations 8) used in deconvolving triple-view line scanning diffraction-limited data were computed in MATLAB (Mathworks, R2019b) for each view.

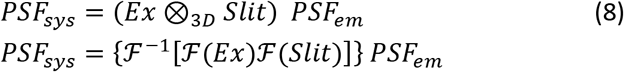

where *Ex* is the 3D line excitation pattern; *Slit* models the rolling shutter, a rectangle function with width equivalent to the number rows in the rolling shutter at the *z* = 0 plane, and defined to be zero at other *z* planes; *PSF*_*em*_ the 3D emission PSF; and ⊗_*3D*_ the 3D convolution operation, implemented in Fourier domain (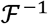 denotes Fourier transform and 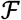 inverse Fourier transform).

Assuming the imaging system is aberration free, we derived *Ex* and *PSF*_*em*_ by inverse Fourier transforming the scalar pupil function (Equations 9^26^):

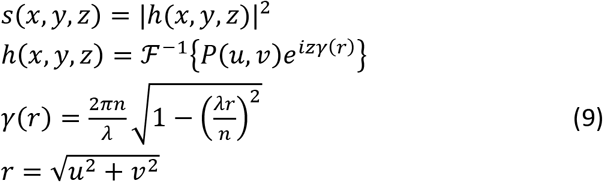

where *S*(*x, y, z*) and *h*(*x, y, z*) are the incoherent and coherent PSFs; *P*(*u*, *v*) is the pupil function, *u, v* are pupil plane coordinates, *γ*(*r*) is the defocus function, *r* is the radial coordinate in the pupil plane coordinate; *x, y* are lateral coordinates and *z* the axial coordinate in the spatial domain, *λ* the wavelength, and *n* the refractive index.

For line excitation, we model the excitation pupil function *P*_*ex*_(*u, v*) as a single line centered at *u* = 0, bounded by the cutoff frequency of the objective:

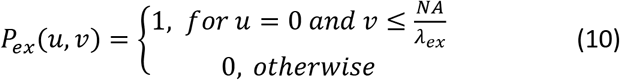

where *NA* is the numerical aperture of the objective and *λ*_*ex*_ is the excitation wavelength.

For the detection, we model the pupil function in focus *P*_*em*_(*u, v*) as a binary mask that transmits all spatial frequencies transmitted by the objective:

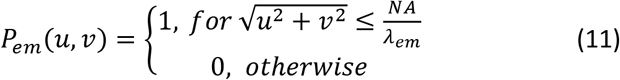

where *NA* is the numerical aperture of the objective and *λ*_*em*_ is the emission wavelength.

When replacing *P*(*u, v*) in Equations 8 with *P*_*ex*_(*u, v*) (Equation 10) and *P*_*em*_ (*u, v*) (Equation 11), we obtain *Ex* and *PSF*_*em*_, and thus *PSF*_*sys*_ according to Equations 8. We set *NA* to 0.8 for the top views and 1.2 for the bottom view, *n* to 1.33 (water), *Slit* at the in-focus plane was a rectangular window with five pixels width (pixel size 97.5 nm), excitation wavelength to 530 nm and emission wavelength to 590 nm (to best match measured PSFs derived from 100 nm fluorescent beads). We normalized PSFs to the total energy (i.e., divided each by the sum of all PSFs). Top PSFs were rotated to the conventional viewing direction, i.e., visualized from the bottom view.

To deconvolve SIM images, the system PSFs for each view were simulated based on the SIM image formation process described in *1D SIM reconstruction*. For a given position of the line illumination, the detected fluorescence image of the sample (here assumed to be a 1 voxel source) is given by Σ_*z*_[(*Ex* x *Sample*) ⊗_*2D*_ *PSF*_*em*_] x *Slit*_*2D*_, where we loop over *z* in *Ex*, *Sample*, and *PSF*_*em*_, perform the 2D convolution ⊗_*2D*_ for each *Z* plane, sum over the *Z* coordinate, and multiply with the 2D slit function (i.e., *Slit*_*2D*_, a rectangle function of width equal to the number of rows in the rolling shutter). In 1D SIM mode, the line excitation is shifted row by row along the scanning direction, and modulated (on for 1 row and off for *N* − 1 rows, where *N* is the number of phases); thus we obtained a phase image *I*_*n*_ on the camera by accumulating the detected fluorescence while shifting *Ex* and *Slit*_*2D*_ in synchrony. This process was repeated for all phases to generate *N* phase images (Equation 12):

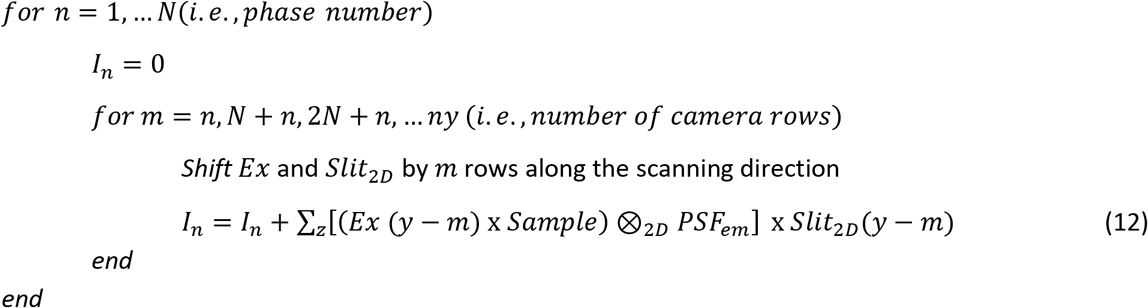

We applied the digital photon reassignment procedure (described in *1D SIM reconstruction*) to all *N* images, thereby reconstructing a SIM image of the source. We repeated this process for all *Z* slices by axially translating the point source, thus deriving the system 3D PSF in 1D SIM mode. Equation 12 was also used to simulate the fluorescence patterns (**Fig. S4c**) in evaluating the optimum phase number *N* required for 1D SIM reconstruction. In this case, the sample was modeled as a uniform layer (i.e., one pixel thick) of fluorophores, and the simulation was performed on the in-focus plane. The parameters used in this simulation were the same as for deriving PSFs.

### Neural networks for image restoration and segmentation

We used four neural networks including (1) content aware image restoration (CARE^36^) networks to denoise the raw, low SNR images of developing nuclei and neurons in *C. elegans* embryos captured in each view, and to predict the high SNR triple-view deconvolution output from the raw View C (**Fig. 3d-j**, **Fig. S3e-h**); (2) DenseDeconNet^21^ for predicting the diSPIM output from single-view light sheet images of embryonic nuclei (**Fig. 3f, Fig. S3k**); (3) a modified segmentation routine based on Mask-RCNN^30^ for segmenting nuclei in adult worms and embryos imaged with the line scanning confocal system and diSPIM (**Fig. 2f, Fig. 3f, Fig. S3i-k**); (4) a 3D residual channel attention network (RCAN^38^) to predict 1D super-resolution images from diffraction-limited input and derive single-view 2D SIM (**Fig. 5, Fig. S6a-b, Fig. S6d**) and triple-view 2D SIM (**Fig. 6, Fig. S6e-f**) reconstructions. Mostly, model training and application were performed on an NVIDIA TITAN RTX GPU installed on a local workstation. We also performed some RCAN training and model application in the Google Cloud Platform (GCP), using a virtual machine with two NVIDIA Tesla P100 GPUs (each with 16GB memory). We used Tensorflow framework version 2.4.1 and Python version 3.7.1. Peak-signal-to-noise-ratio (PSNR) and the structural similarity index (SSIM) of the network output and ground truth (**Fig. S3f**) were evaluated with built-in MATLAB (Mathworks) functions.

CARE software was installed from GitHub (https://github.com/CSBDeep/CSBDeep). For training, patches of size 64 × 64 × 32 were randomly cropped from the data, and 10% of the patches set aside for validation. For the embryonic nuclei data, we used 100 matched volumes (low SNR ^~^4 to high SNR ^~^60, volume size 320 x 220 x 40 voxels, SNR values were estimated by averaging pixel values in regions surrounding nuclei, and dividing by the standard deviation of the pixel values in a background region outside the embryo) for Views A/B, and 100 matched volumes for View C (size 350 x 540 x 32 voxels) to train a denoising model for each view; then 68 matched volumes (size 350 x 540 x 328 voxels) to train a second network to predict the triple-view joint deconvolution result from the denoised View C data. For embryonic neuron data, 40 matched volumes (low SNR ^~^4 to high SNR ^~^40, SNR estimated by computing the average intensity value along a neurite, dividing by the standard deviation of the pixel values in a background region outside the embryo) were used to train denoising models for each view (450 x 330 x 48 voxels for Views A/B and 380 x 640 x 40 voxels for View C); 40 matched volumes were also used for predicting the triple-view deconvolved result from the denoised View C data (volume size of 380 x 640 x 411 voxels).The training learning rate was 2 × 10^4^, the number of epochs 100, the number of steps per epoch 200, and mean absolute error (MAE) was used as loss function. Training took a total of ^~^3 days for all models, and applying the trained models took ^~^ 20 seconds with the two-step deep learning strategy.

For the studies using DenseDeconNet, we used our published single-input neural network (https://github.com/eguomin/regDeconProject/tree/master/DeepLearning). The objective function incorporated three terms: the mean square error (MSE), the structural similarity (SSIM) index and the minimum value of the output (MIN). The input data was raw single-view light sheet image volumes, and the ground-truth data consisted of jointly deconvolved diSPIM data. Training and validation data with size 240 x 360 x 240 voxels were derived from 180 volumetric time points, and cropped into subvolumes with size 120 x 120 x 240. 80% of the volumes were randomly selected for training and the remaining 20% were used for validation. The learning rate was 0.04 and the training iteration number was 7000. It took ^~^2.5 h to train the network, and ^~^ 1 s per volume for applying the trained network.

For Mask RCNN^30^, which is a state-of-the-art segmentation framework, we adopted the Keras and Tensorflow based implementation (https://github.com/matterport/Mask_RCNN) to perform nuclear segmentation. We manually segmented 8 volumes (3 diSPIM, 3 iSPIM, and 2 triple-view confocal, 1963 nuclei in total) for training. Of these 8 volumes, 6 volumes with 1688 nuclei were used for training and 2 volumes with 275 nuclei were used for validation. We used a ResNet-50 feature pyramid network model as the backbone, initialized the model using weights obtained from pretraining on the MS COCO^31^ dataset and proceed to train all layers in three stages. In total we trained for 180 epochs using stochastic gradient descent with a momentum of 0.9, starting with a learning rate of 0.001 and ending with a learning rate of 0.0001. We used a batch size of 2 on a single NVIDIA GPU P6000. Gradients were clipped to 5.0 and weights decayed by 0.0001 each epoch. Training took ^~^10 hours and applying the model took ^~^ 3 minutes per volume. After nuclear segmentation by Mask RCNN, we also applied a marker-controlled watershed to separate touching nuclei. In this post-processing step the minima of the distance transform of each Mask RCNN output mask were extracted to replace region minima. Then, the watershed transformation of the distance transforms was used to separate touching nuclei (**Fig. S3j**).

For the studies employing RCAN, we used our recently developed 3D RCAN model, appropriate for restoring image volumes (https://github.com/AiviaCommunity/3D-RCAN)^38^. For the simulated 2D phantoms (**Fig. 5b and Fig. S6a**), patches of size 256 × 256 were randomly cropped from 1000 2D frames with size 512 x 512 pixels (see the section *Simulation of mixed structures for 1D super-resolution prediction*). For all 3D data, patches of size 128 × 128 × nz (nz is the number of planes) were randomly cropped from the data. Training used 50 volumes (each 1600 x 1600 x 40 voxels) for the cells expressing microtubule-GFP (**Fig. 5c**), 73 volumes (each 416 x 620 x 40 voxels) for the cells expressing EMTB-3XGFP (**Fig. 5f**), 51 volumes (each with size 604 x 766 x 40 voxels) for the cells expressing F-actin-tdTomato (**Fig. 5f**), 139 volumes (each with size 272 x 360 x 32 voxels) for the cells expressing H2B-GFP (**Fig. 5g**), 108 volumes (each with size 356 x 430 x 32 voxels) for the cells expressing EMTB-mCherry (**Fig. 5g**), 26 volumes (each with size 730 x 440 x 32 voxels for Views A/B, 800 x 1200 x 48 voxels for View C) for the larval worms expressing membrane targeted GFP (**Fig. 6b, Fig. 6d-e**), and 50 volumes (each with size 640 x 800 x 64 voxels for Views A/B, and 1786 x 1260 x 16 voxels for View C) for the cells expressing actin-GFP (**Fig. 6k**). In all RCAN training, the learning rate was 2 × 10^−4^, the number of epochs for training 200, the number of residual blocks 5, the number of residual groups 5, the number of channels 32, the steps per epoch 400; and the MAE was used as the loss function. The training time for each model varied from ^~^3-50 hours. For example, it took ^~^5 h for the Jurkat T cell dataset expressing H2B-GFP (**Fig. 5g**), and ^~^4 hours to apply the model to recover a 100 time-point dataset with size 430 x 376 x 32 voxels and 6 rotations (total 600 volumes), including the processing time for image rotation, network prediction, rotation back, and file reading/writing.

### Simulation of mixed structures for 1D super-resolution prediction

We simulated 2D images (1000 pairs for training, and 50 for validation) with mixed structures of dots, lines, rings, and solid circles for validating the deep learning method that provides isotropic, super resolution from diffraction-limited input (**Fig. 5b, Fig. S6a**). Simulated structures were created in MATLAB (Mathworks, R2019b, with the Imaging Processing Toolbox). Each image was composed of 150 lines, 50 dots, 150 solid circles, and 150 rings, randomly located in a 1024 x 1024 grid. The 150 lines were generated with random angles (1-360 degree), random lengths (1-200 pixels), and a random intensity (10-60 counts); the 50 dots were generated with random intensity (30-60 counts); the 150 solid circles were generated with random diameter (1-25 pixels) and random intensity (10-60 counts); the 150 rings were generated with random outer diameter (1-35 pixels), random thickness (1-3 pixels) and random intensity (40-100 counts). Structures were blurred with a 2D Gaussian function (sigma = 1.5 pixels in both x and y dimensions), then shrunk 2-fold to 512 x 512 pixels, mainly to slightly smooth the generated structures. These smoothed structures served as ‘simulated objects’, assuming a pixel size of 30 nm (**Fig. 5b**). Then we generated two sets of images: one blurred with a 2D Gaussian function (sigma = 4 pixels in both x and y dimensions), normalized, corrupted with Gaussian (mean = 0; variance = 0.0005) and Poisson noise, serving as the ‘simulated raw input’ (**Fig. 5b**); the other blurred with a 2D Gaussian function (sigma = 4 pixels in x, 2 pixels in y), normalized, and again corrupted with Gaussian (mean = 0; variance = 0.0005) and Poisson noise, serving as the 1D SIM image (i.e., the ground truth for training).

### Estimating resolution

Lateral and axial resolution measures derived from fluorescent beads and actin fibers (**Fig. 6k-p, Fig. S6f**) were estimated by computing the FWHM of line intensity profiles along XY and XZ views. The thickness of the line excitation pattern (**Fig. S1e**) was also estimated by finding the FHWM of the intensity of a bead as a function of scan position through it. Statistical results (mean ± standard deviation) were obtained from *N* = 8-40 samples. All FWHM calculations were obtained in MATLAB, after interpolating line intensity profiles 10-fold. For rough resolution comparisons, we plotted the Fourier transforms of the images (**Fig. 1d, Fig. 4b, Fig. S4b, Fig. S4f, Fig. S6a-b**).

All other resolution estimates were based on decorrelation analysis^27^. This method estimates average image resolution from the local maxima of a series of decorrelation functions, providing a metric that corresponds to the highest spatial frequency with sufficient SNR, rather the Abbe resolution limit. For estimating anisotropic resolution in a given plane (**Fig.4g, Fig. 5b-c**), we performed a modified sector-based decorrelation analysis^38^, in which Fourier space was subdivided into 16 sectors (each with 22.5 degrees angular span). For estimating lateral and axial resolution (e.g., for **Fig.1e, Fig. S6e**), we first interpolated the stacks along the axial dimension to achieve isotropic pixel size; then we performed sectorial resolution estimate on xy and xz slices (with mean and standard deviations derived from the slices), by using three sectorial masks to capture spatial frequencies predominantly along the x, y, and z dimensions.

### Estimating expansion factor

For the expanded cells (**Fig. 1e, Fig. 4g**), the expansion factor was estimated to be 3.9-4.1, by comparing pre- and post-expansion data in 3D Slicer after performing landmark-based registration^38^. For the expanded embryo stained with DAPI (**Fig. 4i, Fig. S5b, c**), the expansion factor was estimated by imaging and registering 7 pre- and post-expansion volumes. We first unsampled the pre-expanded embryo image volume 3.2-fold, manually rotated the volume to coarsely overlap with the post-expanded data, and finally registered the coarsely aligned volumes with a GPU-based 12-degree affine registration method^21^. The normalized cross correlation of the volumes after registration was 0.72 ± 0.03. We then computed the expansion factor Ē-from the following operations:

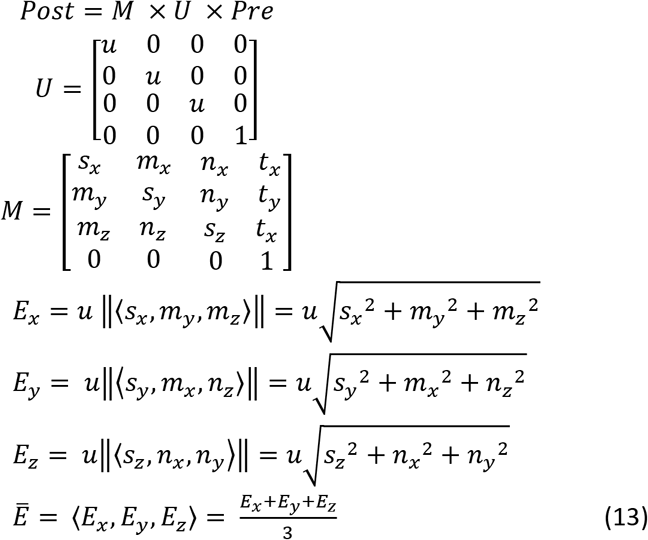

where *U* describes the 3.2-fold upsampling (i.e., *u* = 3.2), *M* is the transformation matrix (extracted from the affine registration) that maps the 3.2x upsampled pre-expansion data (*Pre*) to post-expansion data (*Post*), *E*_*x*_, *E*_*y*_, *E*_*z*_ represents the scaling factor for x, y, and z dimensions, respectively; and Ē-is the average scaling factor. Our results show that the expansion factor for whole embryos is 3.29 ± 0.14, with nearly isotropic expansion (3.24 ± 0.14, 3.39 ± 0.14, 3.26 ± 0.17 for x, y and z dimensions, respectively).

### Point scanning confocal imaging for comparison with triple-view line confocal imaging

A commercial point scanning confocal microscope (Leica SP8, 60x/1.2 NA water lens) was used to image a whole fixed adult *C. elegans* (strain EG6994) labeled with NucSpot Live 488, for comparison with triple-view line confocal imaging. Point scanning confocal volumes (**Fig. S2a**) were acquired in ^~^70 mins (stitched from 10 subvolumes with size 1024 x 1024 x 192 voxels, and 10 subvolumes with size 1024 x 1024 x 60 voxels, with a resonant scanning rate of 600 Hz so that the imaging rate was ^~^1.7s per frame). For each subvolume, the xy pixel size was 100 nm per pixel, and z 200 nm. After stitching and z interpolation, the final resulting volume size was 8364 x 1024 x 500 voxels (corresponding to ^~^836 x 102 x 50 μm). Comparative triple-view line confocal volumes (**Fig. 2h, Fig. S2c**) were acquired in ^~^160 s (stitched from 10 subvolumes, 16 s per subvolume). For each subvolume, we acquired 56 planes at 1 μm spacing for all three views. Post triple-view deconvolution of each subvolume and stitching, the final volume size was 8930 x 1280 x 574 voxels (corresponding to ^~^ 870 μm x 125 μm x 56 μm). Table S1 provides further details.

### Spinning disk confocal imaging for comparison with triple-view line confocal imaging

An Andor spinning disk confocal microscope mounted in a Nikon Eclipse Ti frame and equipped with iXon 897 EM-CCD camera and 40x 1.3 NA oil immersion objective (resulting in a pixel size of 333.3 nm), was used to image *Drosophila* larvae for comparison with the triple-view line confocal system. Spinning disk dual-color volumes (488 nm and 561 nm excitation, **Fig. S2d**) with 295 planes (each 512 x 512 pixels, 100 ms exposure time and 299 EM gain) and a spacing of 0.395 μm were acquired in 59 s. Triple-view line confocal volumes (**Fig. 2h, Fig. S2c**) were also acquired in 59 s (160 planes at 1 μm spacing for each top view in stage-scanning mode, and 120 planes at 0.5 μm spacing for the bottom view).

### diSPIM imaging for comparison with triple-view line confocal imaging

A fiber-coupled diSPIM^23,39^ with two 40x, 0.8 NA water objectives (Nikon Cat. # MRD07420), resulting in a pixel size of 162.5 nm, was used to image transgenic embryos (strain RW10230 expressing GFP-nuclei, and strain DCR6633 expressing GFP-AIB neurons, described in detail in the *Worms* section). When imaging nuclei (**Fig. 3f**), stacks were also acquired from both views (50 planes at 1 μm spacing per view, 0.91 s per dual-view volume) at 3-minute intervals until the embryo hatched (*N* = 16 embryos). In imaging AIB neurons (**Fig. 3i**), dual-view stacks were acquired (50 planes at 1 μm spacing per view, 1.75 s per dual-view volume) at 3-minute intervals for 540 minutes. Embryonic twitching was stereotyped and started ^~^435 minutes post fertilization (m.p.f) under our imaging conditions. The images were rotated to represent lateral views of the AIB neuron.

Volumetric time-lapse imaging of the same strains was also conducted with View C in the triple-view line confocal system (0.6 s per volume for nuclear imaging, every 3 mins, N = 16 embryos, **Fig. 3e**; 0.75 s per volume for AIB neurons, every 5 mins, N = 10 embryos, **Fig. 3h, i). Table S1** provides further details.

### iSIM imaging for comparison with triple-view SIM

A home built iSIM system^10^ with 60X, NA = 1.2 water objective (Olympus UPLSAPO60XWPSF) and an sCMOS camera (PCO, Edge 5.5), resulting in a pixel size of 55 nm, was used to image stained cells in gels (**Fig. 4f**) and C. *elegans* larvae (**Fig. 6d**) for comparison with line confocal SIM. All images were background subtracted and deconvolved using the Richardson-Lucy algorithm with 15 iterations.

Collagen gel samples containing fixed B16-F10 cells stained with MitoTracker Red CMXRos and Alexa Fluor 488 phalloidin were placed in an imaging chamber filled with PBS. Fiducial marks were made on the coverslip so that we could find the same cells on both the triple-view system and the iSIM. We acquired triple-view 1D SIM volumes of the cell in 14.5 s (72 planes at 1 μm spacing for each top view, 32 planes at 0.5 μm spacing for the bottom view, **Fig. 4e**) and iSIM volumes of the cells in 2 s (40 planes at 0.25 μm spacing, **Fig. 4f**).

A DCR8528 L4 stage larval worm was transferred from an agar plate into a drop (^~^10 μL) of UV-curable polymer BIO-133^56^ (MY Polymers Ltd.) on a #1.5 cover glass (24 mm x 50 mm, VWR, 48393-241). The droplet was positioned between two 50 μm spacers (Precision Brand, 44910), and was compressed by a glass slide followed by 2-minute polymerization under a UV lamp (365nm, Spectroline ENB-280C). The immobilized worm was placed in the imaging chamber, and M9 buffer containing 0.25% sodium azide was added to further stop the worm from moving. The same chamber was used for imaging the larva on both the triple-view system and the iSIM. Triple-view volumes (diffraction-limited mode) were acquired in 1.5 s (40 planes at 1 μm spacing for each top view, 56 planes at 0.5 μm spacing for the bottom view, **Fig. 6e**) and iSIM volumes in 4.25 s (85 planes at 0.3 μm spacing, **Fig. 6d**).

### Traditional 3D SIM imaging for comparison with triple-view SIM

Worm larvae (strain DCR6681, described in detail in *Worms* section) were ‘hot-fixed’ in 4% PFA in PBS at 50°C for 5 minutes and then washed three times using PBS. A commercial OMX-SR 3D SIM system equipped with a 60× 1.3 NA silicone objective was used for imaging the fixed worms with a raw pixel size of 80 nm (**Fig. S6c**). We put a small drop of PBS containing fixed worms between a glass slide and a coverslip and imaged through the coverslip using the sample holder provided with the OMX. 169 z slices (1024 x 1024 pixels, 3 angles and 5 phases for each z slice) with a spacing of 0.125 μm were acquired in 25 s, while the reconstruction (**Fig. S6c**, softWoRx 7.2.0 Suite) was carried out on the 40 slices (5 μm stack) near the coverslip to minimize reconstruction artifacts. The PSF/OTF required for reconstruction was acquired with 100 nm yellow-green beads using the same objective lens and coverslip as used for the larvae.

