## Supplementary Materials for "Multiview super-resolution microscopy"

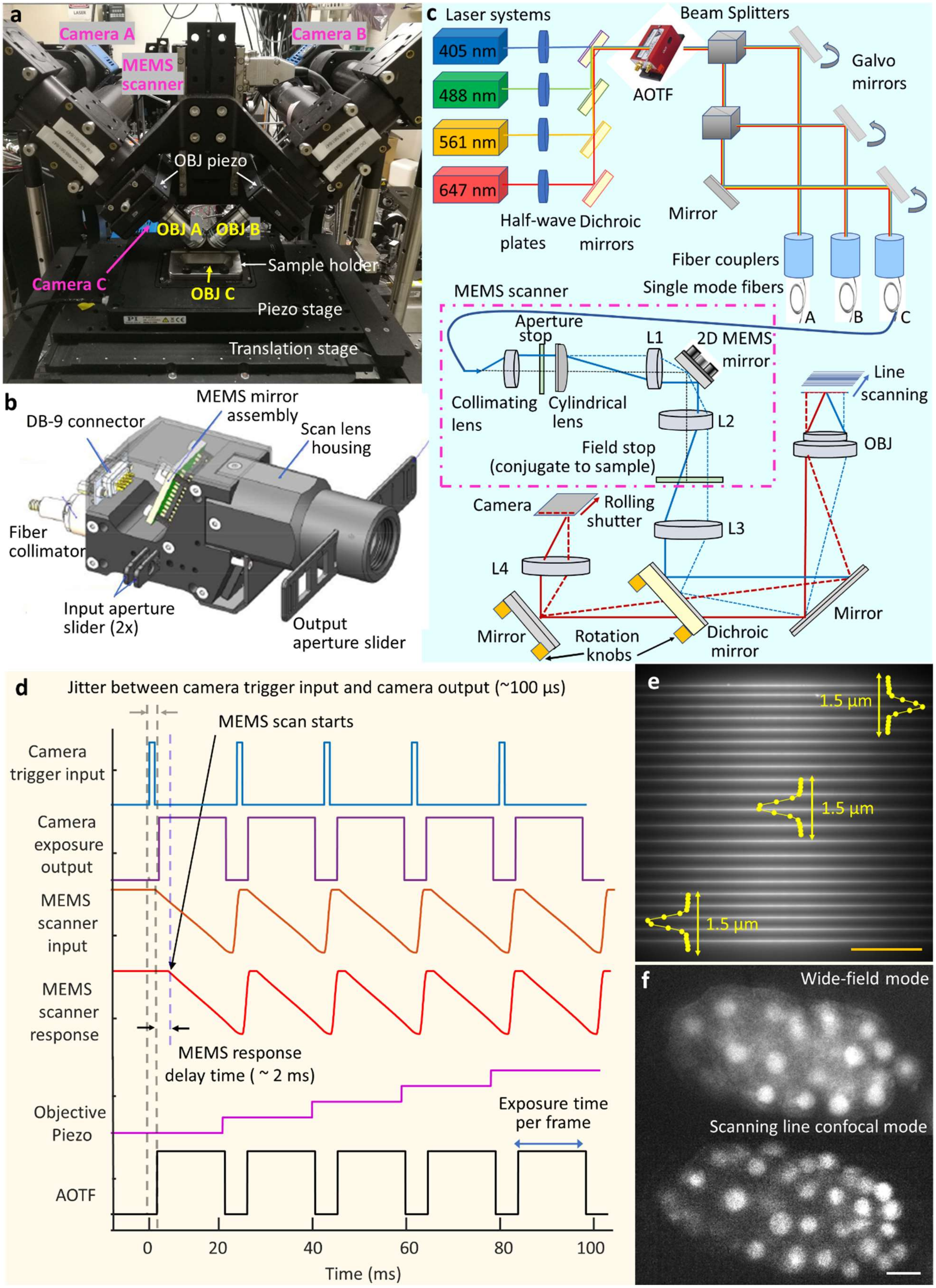
a
Camera A
Camera B
MEMS scanner
OBJ piezo
OBJ A
OBJ B
Sample holder
Camera C
OBJ C
Piezo stage
Translation stage
b
MEMS mirror assembly
DB-9 connector
Scan lens housing
Fiber collimator
Input aperture slider (2x)
Output aperture slider
c
Laser systems
405 nm
488 nm
561 nm
647 nm
Half-wave plates
Dichroic mirrors
AOTF
Beam Splitters
Galvo mirrors
Mirror
Fiber couplers
Single mode fibers
A
B
C
MEMS scanner
Aperture stop
L1
2D MEMS mirror
Collimating lens
Cylindrical lens
L2
Field stop (conjugate to sample)
Line scanning
OBJ
Camera
Rolling shutter
L3
L4
Mirror
Rotation knobs
Dichroic mirror
Mirror
d
Jitter between camera trigger input and camera output (~100 µs)
MEMS scan starts
Camera trigger input
Camera exposure output
MEMS scanner input
MEMS scanner response
MEMS response delay time ( ~ 2 ms)
Objective Piezo
Exposure time per frame
AOTF
0
20
40
60
80
100
Time (ms)
e
1.5 µm
1.5 µm
1.5 µm
f
Wide-field mode
Scanning line confocal mode

1

1

**Fig. S1, Instrument overview, related to Fig. 1**. **a)** Photograph of instrument, highlighting objectives, cameras, sample area. **b)** CAD rendering of MEMS line scanner. **c)** Optical setup. Diode lasers are combined and passed through an acousto-optic tunable filter (AOTF) for shuttering and power control, before being directed into broadband single-mode fibers. Galvanometric mirrors are used to direct beams into each fiber, and to adjust the power of the beams entering the fiber. Each fiber is then fed into a MEMS line scanner (purple dot-dashed line, here only one shown for clarity), with optics as indicated. The scanner serves to collimate the fiber output, focus it with a cylindrical lens, scan it with a MEMS mirror, and image the scanned output to a field stop at a conjugate sample plane. The beam is then relayed via lens L3 and the objective to the sample plane, after reflection from a dichroic mirror. Fluorescence is collected in epi-mode, transmitted through a dichroic mirror, and imaged via tube lens L4 onto a scientific CMOS camera where it is synchronized to the rolling shutter readout. **d)** Control waveforms issued to camera, MEMS scanner, objective piezo, and AOTF, used to acquire volumetric data. See also **Methods**. **e)** Example line illumination at various lateral positions on camera chip, as imaged with fluorescent dye in lower view C. Excitation PSF measurements (FWHM 267 ± 22.5 nm, N = 10 measurements) are taken at various positions in the field. Scale bar: 50 μm. **f)** Example images acquired in *C. elegans* embryo expressing GFP-histones, as visualized in bottom view C with widefield mode (top) and line-scanning mode (slit width 0.58 μm, bottom). Scale bar: 5 μm.

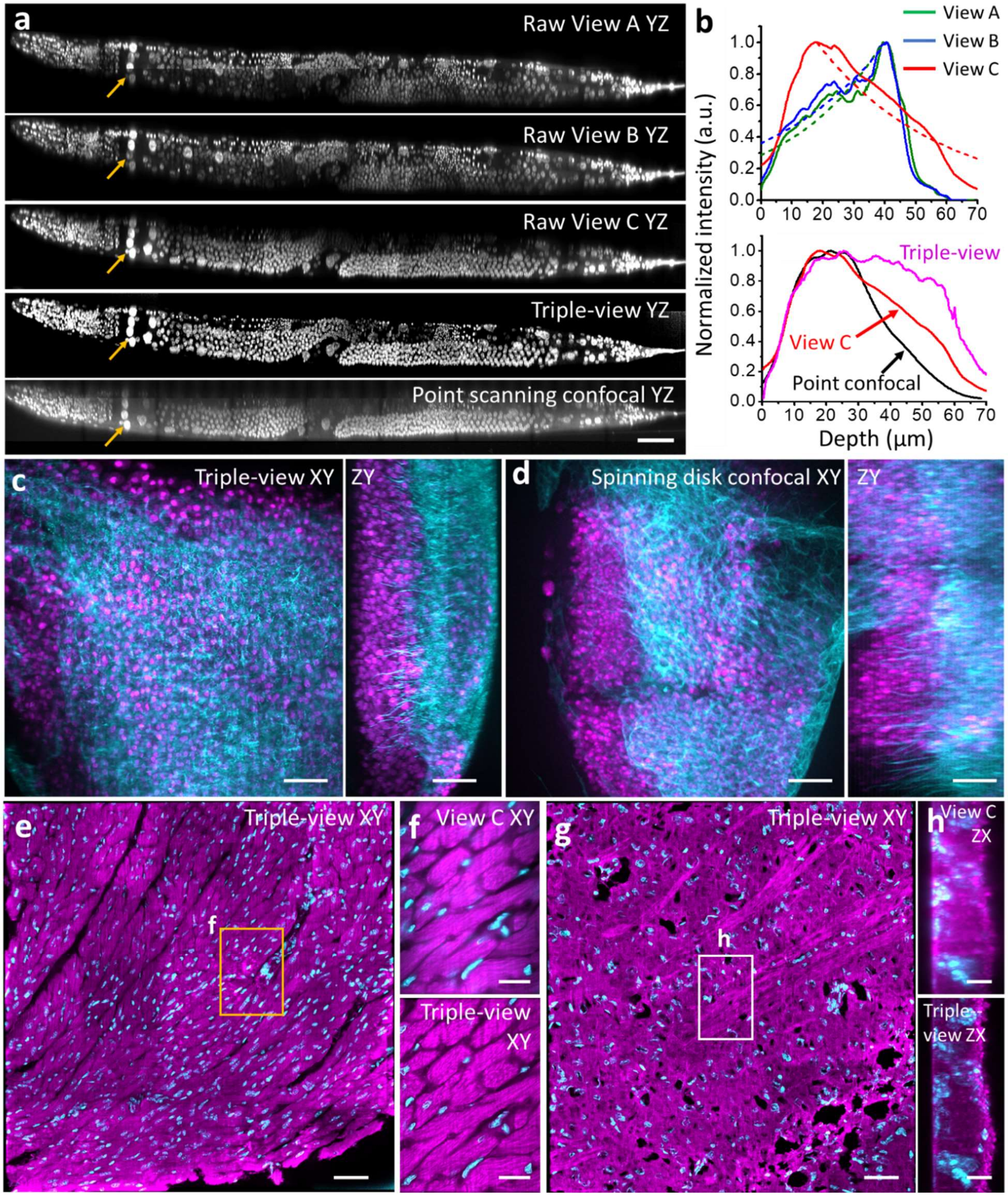


**Fig. S2, Additional triple-view comparisons in scattering tissue, related to Fig. 2. a)** Axial views of whole fixed worm labeled with NucSpot Live 488, comparing raw views gathered by objectives A, B, C; triple-view reconstruction; and point-scanning confocal microscope. Arrows highlight example region with uniformly good quality in triple-view reconstruction, but showing attenuation either at bottom (Views A, B) or top (View C, point-scanning confocal) of stack with other methods. See also **Fig. 2a-e**. Scale bar: 50 μm. **b)** Average attenuation measured through axial extent of the worm, as measured by raw views A, B

C (top graph) and triple-view reconstruction and point-scanning confocal (bottom graph). Exponential fits to the data are also shown with dashed lines. See also **Methods**. **c)** Lateral (left) and axial (right) maximum intensity projections of the larval wing disc shown in **Fig. 2h**. Notum nuclei (NLS-mCherry, magenta) and myoblast membranes (CD2-GFP, cyan) are indicated. **d)** As in **c)** but showing another larval wing disc imaged with a spinning-disk confocal microscope with NA =1.3. Scale bars in **c, d**: 15 μm. **e)** Triple-view reconstruction of mouse cardiac tissue (~536 x 536 x 37 $\mu m^3$ section, shown is a maximum intensity projection of axial slices from 10-30 μm). Magenta: Atto 647 NHS, nonspecifically marking proteins, cyan: NucSpot Live 488, marking nuclei. Scale bar: 50 μm. **f)** Higher magnification view of orange rectangular region in **e)**, comparing raw View C (top) to triple-view reconstruction (bottom). Scale bar: 20 μm. **g)** Labels as in **e)** but showing triple-view reconstruction of mouse brain tissue (~536 x 536 x 25 $\mu m^3$ section, shown is a maximum intensity projection of axial slices from 5-15 μm). Scale bar: 50 μm. **h)** Axial maximum intensity projection computed over 120 μm vertical extent of white rectangular region shown in **g)**, comparing raw View C (top) to triple-view (bottom) result. Scale bar: 10 μm.

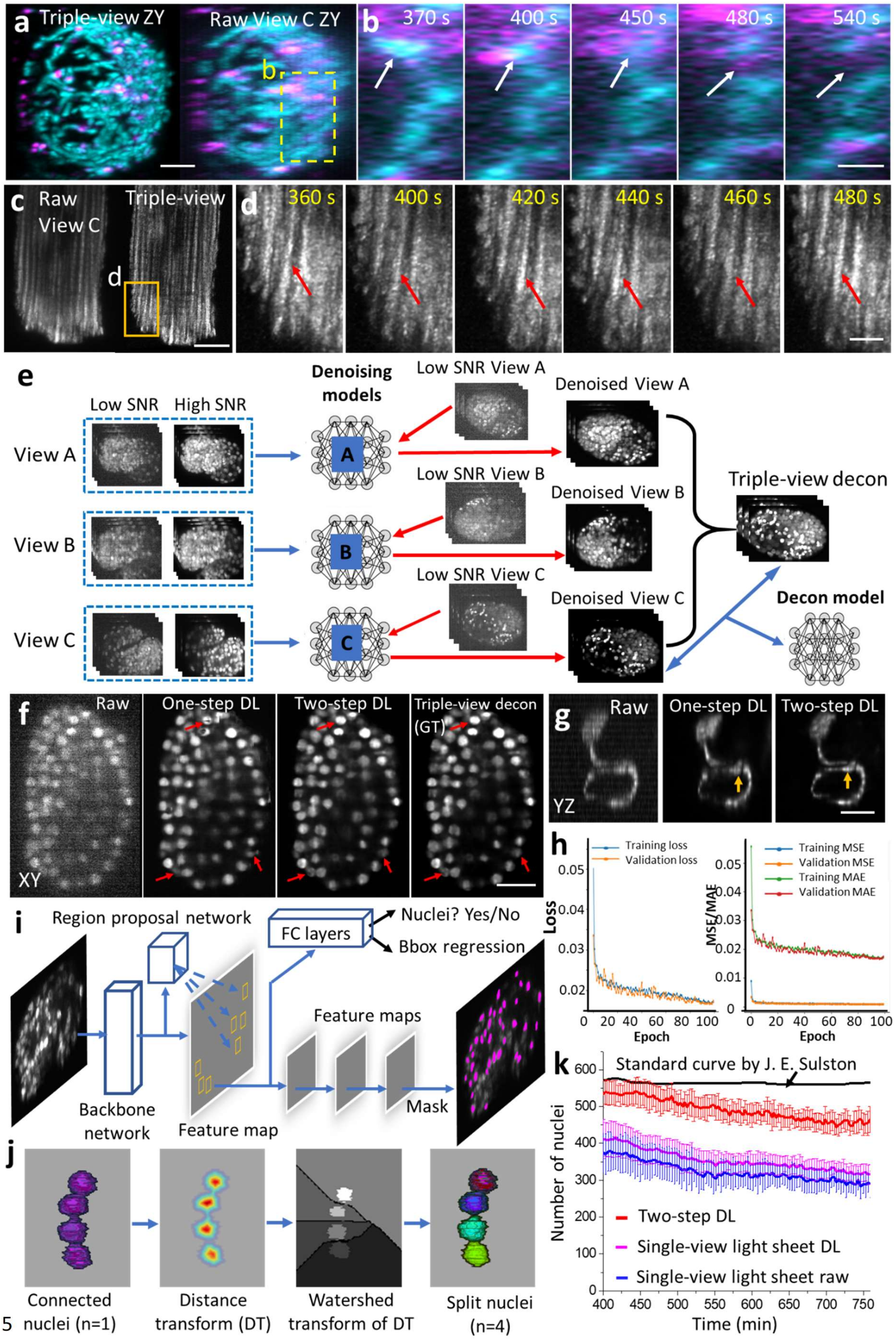
a Triple-view ZY
Raw View C ZY
b
370 s
400 s
450 s
480 s
540 s
c Raw View C
Triple-view
d
360 s
400 s
420 s
440 s
460 s
480 s
e
Low SNR
High SNR
View A
View B
View C
Denoising models
A
B
C
Low SNR View A
Low SNR View B
Low SNR View C
Denoised View A
Denoised View B
Denoised View C
Triple-view decon
Decon model
f
Raw
One-step DL
Two-step DL
Triple-view decon (GT)
XY
g
Raw
One-step DL
Two-step DL
YZ
h
Training loss
Validation loss
Loss
Epoch
Training MSE
Validation MSE
Training MAE
Validation MAE
MSE/MAE
Epoch
i
Region proposal network
FC layers
Nuclei? Yes/No
Bbox regression
Backbone network
Feature map
Feature maps
Mask
j
Connected nuclei (n=1)
Distance transform (DT)
Watershed transform of DT
Split nuclei (n=4)
k
Standard curve by J. E. Sulston
Number of nuclei
Time (min)
Two-step DL
Single-view light sheet DL
Single-view light sheet raw

**Fig. S3, Additional live triple-view data and neural network schematics, related to Fig. 3**. **a)** Lysosomes (magenta) and mitochondria (cyan) in HCT-116 human colon carcinoma cells, comparing triple-view axial maximum intensity projection (left) to raw View C data (right). See also **Fig. 3a**. Scale bar: 3 μm. **b)** Higher magnification views of white dashed rectangular regions in **a)**. Scale bars: 2 μm. See also **Fig. 3b. c)** Maximum intensity projection of cardiomyocyte stained with MitoTracker Red CMXRos, shown in **Fig. 3c**. Raw View C (left) is also shown for comparison. Scale bar: 10 μm. **d)** Higher magnification view of orange rectangular region in **c)**, highlighting mitochondrial fluctuations (red arrows) over time. Scale bar: 3 μm. **e)** Workflow for two-step deep learning procedure used in **Fig. 3d, e**. Denoising neural networks are trained for views A, B, C, by using matched high and low SNR volumes derived from embryos paralyzed with sodium azide. The denoising output of each model are combined with joint deconvolution (triple-view decon) and are combined with the denoised view C output to train a second neural work (Decon model). In this manner, noisy raw data from View C can be transformed into a high SNR, high resolution prediction. **f)** Example data (lateral slice through GFP-histone expression *C. elegans* embryo) from left to right: raw View C data; single neural-network prediction (raw input View C to high SNR, triple-view deconvolved result); two-step denoised and deconvolved prediction; high SNR denoised triple-view deconvolved result (i.e., ground truth used in training the second neural network in **e)**). The two-step prediction is noticeably closer to the ground truth (red arrows highlight regions for comparison; 3D SSIM 0.89 ± 0.05, PSNR 43.4 ± 2.2 (N = 7 embryos) than the single network (3D SSIM 0.72 ± 0.17, PSNR 27.8 ± 6.4, N = 7). Scale bar: 10 μm. See also **Fig. 3e**. **g)** Example raw input (left), output of single neural network (middle), and two-step output (right) for AIB neuron. Axial views are shown, orange arrows that fine features are better preserved in two-step rather than one-step neural network. Scale bar: 5 μm. See also **Fig. 3h, i**. **h)** Training and validation loss (left) and error (right) as a function of epoch number, for the second step in the neural network, i.e., denoised view C input and triple-view deconvolved output in **e)**. MSE: mean square error. MAE: mean absolute error. **i)** Mask RCNN used for segmenting nuclear data in **Fig. 3**. Key components of the Mask RCNN include a backbone network, region proposal network (RPN), object classification module, bounding box regression module, and mask segmentation module. The backbone network is a convolutional neural network that extracts features from the input image. The RPN scans the feature map to detect possible candidate areas that may contain objects (nuclei). For each bounding box containing an object, the object classification module (containing fully connected (FC) layers) classifies objects into specific object class(es) or a background class. The bounding box regression module refines the location of the box to better contain the object. Finally, the mask segmentation module takes the foreground regions selected by the object classification module and generates segmentation masks. **j)** Post processing after Mask-RCNN. Nuclei (here four are shown) are often connected and need to be split. We apply a watershed to split the nuclei based on a distance transform. **k)** Number of nuclei segmented by two-step deep learning, raw single-view light-sheet imaging, and single-view light-sheet imaging passed through DenseDeconNet, a neural network designed to improve resolution isotropy. See also **Fig. 3f**.

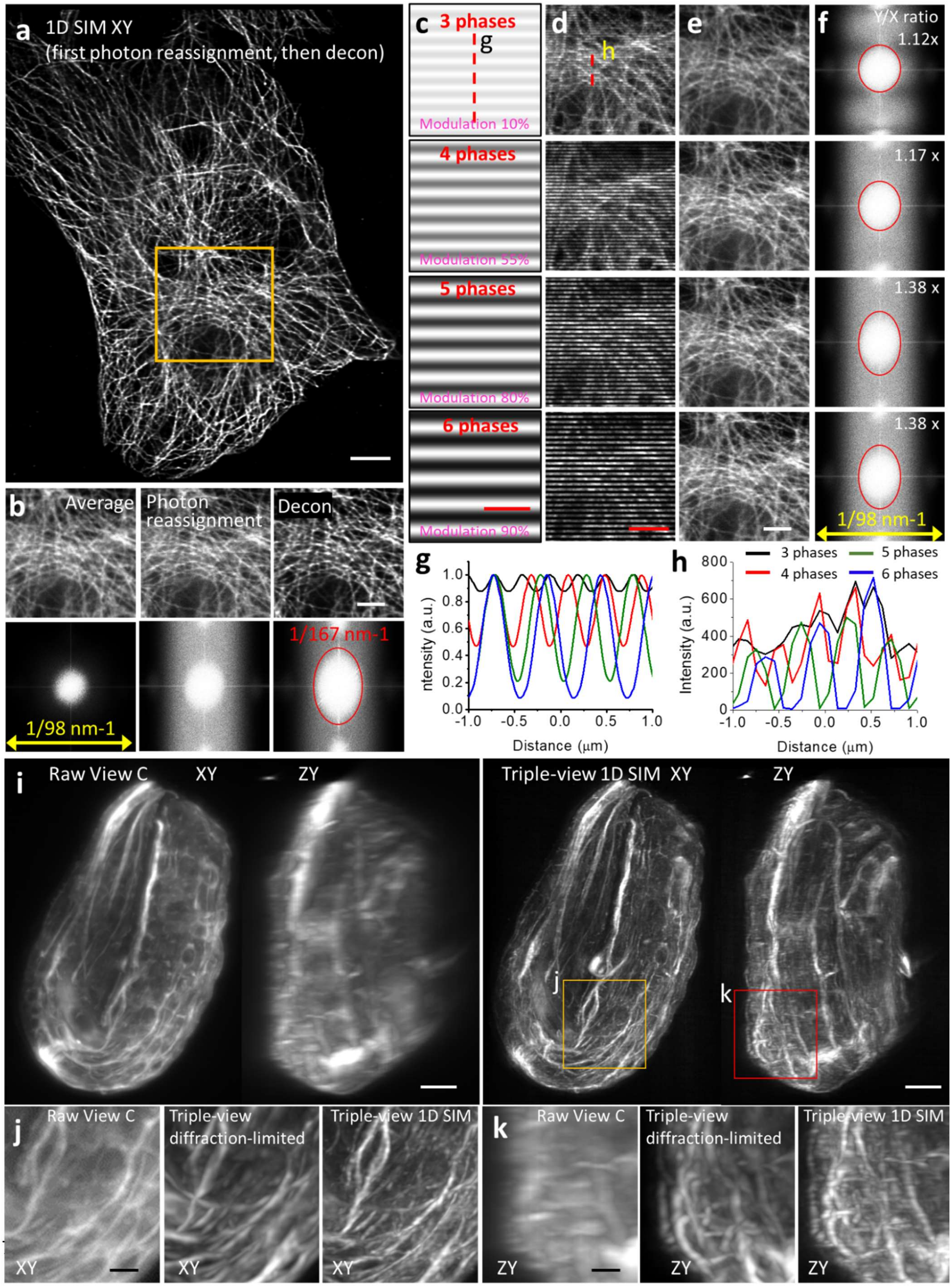
a
1D SIM XY
(first photon reassignment, then decon)
b
Average
Photon reassignment
Decon
1/98 nm-1
1/167 nm-1
c
3 phases
g
Modulation 10%
4 phases
Modulation 55%
5 phases
Modulation 80%
6 phases
Modulation 90%
d
h
e
f
Y/X ratio
1.12x
1.17 x
1.38 x
1.38 x
1/98 nm-1
g
Intensity (a.u.)
Distance (μm)
h
3 phases
4 phases
5 phases
6 phases
Intensity (a.u.)
Distance (μm)
i
Raw View C
XY
ZY
Triple-view 1D SIM
XY
ZY
j
k
j
Raw View C
Triple-view diffraction-limited
Triple-view 1D SIM
XY
XY
XY
k
Raw View C
Triple-view diffraction-limited
Triple-view 1D SIM
ZY
ZY
ZY

**Fig. S4, Triple-view 1D SIM data and methods, related to Fig. 4. a)** Immunolabeled microtubules in a fixed U2OS cell, imaged from the bottom view in 1D SIM mode (View C), as in **Fig. 4b**. Photon reassignment is used to improve resolution, followed by deconvolution. Single lateral plan from image volume is shown. See also **Methods**. Scale bar: 5 μm. **b)** Orange rectangular region in **a)**, comparing line confocal image obtained after averaging 5 phases (left), photon reassignment (middle), and photon reassignment followed by deconvolution (Decon, right). Scale bar: 3 μm. Corresponding Fourier transforms are shown at bottom, red ellipse indicates extended resolution with 167 $nm^{-1}$ vertical extent. **c)** Simulated fluorescence patterns elicited by (top to bottom) 3, 4, 5, 6 phase illumination. Modulation contrast for each pattern is also indicated. Scale bar: 1 μm. See also **Methods**. **d)** Raw data corresponding to orange rectangular region in **a),** as excited with 3, 4, 5, 6 phase illumination. **e)** Corresponding deconvolved result after photon-reassignment. Scale bars in **d-e**): 3 μm. **f)** Fourier transforms corresponding to **e)**. Vertical/horizontal resolution improvement corresponding to red ellipse also indicated. **g)** Line profiles for 3, 4, 5, 6 phase illumination, corresponding to dotted red line in **c)**. **h)** As in **g)** but corresponding to dotted line in **d)**. **i)** Fixed *C. elegans* embryo (strain DCR6681) with tubulin immunolabeled with α-alpha tubulin primary, Streptavidin Alexa Fluor 568, imaged via lower View C (left) and triple-view 1D SIM mode (right). Lateral (left) and axial (right) maximum intensity projections are shown in each case. Scale bars: 5 μm. Higher magnification lateral **j)** views and axial **k)** views of yellow and red rectangular regions in **i)** are also indicated, highlighting progressive improvement in resolution from view C (left), to triple-view diffraction-limited result (middle), to triple-view 1D SIM result (right). Scale bars: 2 μm.

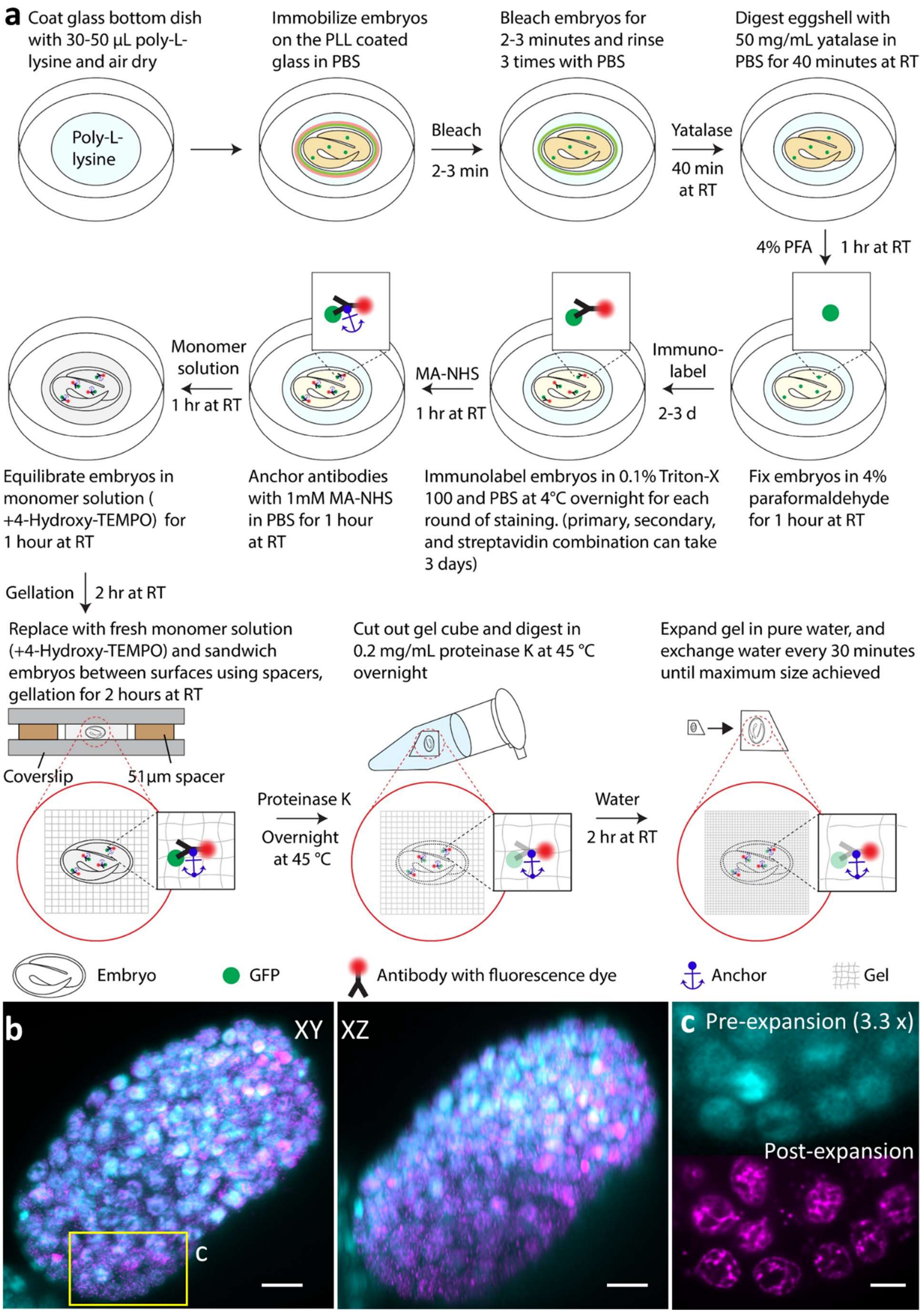
a
Coat glass bottom dish with 30-50 µL poly-L-lysine and air dry
Immobilize embryos on the PLL coated glass in PBS
Bleach embryos for 2-3 minutes and rinse 3 times with PBS
Digest eggshell with 50 mg/mL yatalase in PBS for 40 minutes at RT
Poly-L-lysine
Bleach
2-3 min
Yatalase
40 min at RT
4% PFA
1 hr at RT
Monomer solution
1 hr at RT
MA-NHS
1 hr at RT
Immuno-label
2-3 d
Equilibrate embryos in monomer solution (+4-Hydroxy-TEMPO) for 1 hour at RT
Anchor antibodies with 1mM MA-NHS in PBS for 1 hour at RT
Immunolabel embryos in 0.1% Triton-X 100 and PBS at 4°C overnight for each round of staining. (primary, secondary, and streptavidin combination can take 3 days)
Fix embryos in 4% paraformaldehyde for 1 hour at RT
Gellation
2 hr at RT
Replace with fresh monomer solution (+4-Hydroxy-TEMPO) and sandwich embryos between surfaces using spacers, gellation for 2 hours at RT
Cut out gel cube and digest in 0.2 mg/mL proteinase K at 45 °C overnight
Expand gel in pure water, and exchange water every 30 minutes until maximum size achieved
Coverslip
51µm spacer
Proteinase K
Overnight at 45 °C
Water
2 hr at RT
Embryo
GFP
Antibody with fluorescence dye
Anchor
Gel
b
XY
XZ
C
c
Pre-expansion (3.3 x)
Post-expansion

**Fig. S5, Expansion workflow and imaging results for *C. elegans* embryos, related to Fig. 4i. a)** Immobilization, permeabilization, fixation, immunostaining, and expansion takes approximately four days. See also **Methods**. **b)** Overlay of embryo stained with DAPI before and after expansion after 12-dgree affine registration (cyan: pre-expansion; magenta: post-expansion). The embryo was imaged with diSPIM. Single-view maximum intensity projection along XY and XZ views are shown for comparison. The registration results of 7 embryos (normalized cross correlation: 0.72 ± 0.03) show that the expansion factor for whole embryos is 3.29 ± 0.14, with nearly isotropic expansion (3.24 ± 0.14, 3.39 ± 0.14, 3.26 ± 0.17 for x, y and z dimensions, respectively). See also **Methods**. Scale bar: 5 µm in pre-expansion units. **c)** Magnified view of the rectangle shown in **b),** showing no visible local distortions in nuclei shape during expansion. Scale bar: 2 µm in pre-expansion units.

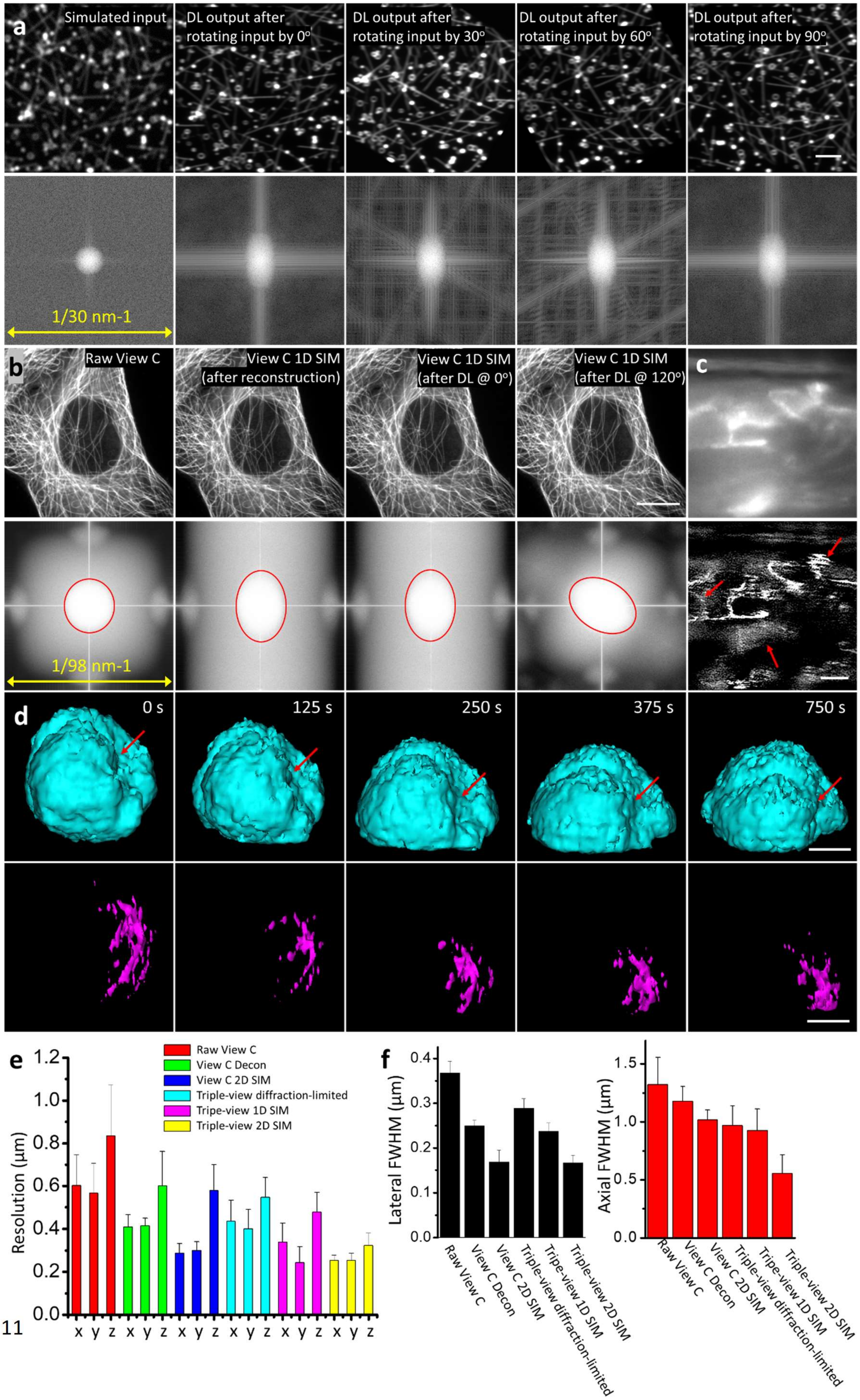

a
Simulated input
DL output after rotating input by 0°
DL output after rotating input by 30°
DL output after rotating input by 60°
DL output after rotating input by 90°
1/30 nm-1
b
Raw View C
View C 1D SIM (after reconstruction)
View C 1D SIM (after DL @ 0°)
View C 1D SIM (after DL @ 120°)
c
1/98 nm-1
d
0 s
125 s
250 s
375 s
750 s
e
Raw View C
View C Decon
View C 2D SIM
Triple-view diffraction-limited
Tripe-view 1D SIM
Triple-view 2D SIM
Resolution (μm)
x y z x y z x y z x y z x y z x y z
f
Lateral FWHM (μm)
Axial FWHM (μm)
Raw View C
View C Decon
View C 2D SIM
Triple-view diffraction-limited
Tripe-view 1D SIM
Triple-view 2D SIM


11

**Fig. S6, Additional multiview super-resolution simulations and experiments, related to Figs. 5, 6. a)** Simulated lines and hollow spheres showing (from left to right): diffraction-limited input; deep learning 1D super-resolution output with no rotation; deep learning output after rotating input by 30 degrees; deep learning output after rotating input by 60 degrees; deep learning output after rotating input by 90 degrees. Fourier transforms in bottom row confirm 1D resolution enhancement regardless of rotation angle. Scale bar: 2 μm. Fourier transforms in bottom row confirm resolution enhancement in images in top row. See also **Fig. 4b. b)** Immunolabeled microtubules in fixed U2OS cells, same sample as shown in **Fig. 4c, d.** Shown are raw input (View C); 1D SIM output after physics-based reconstruction with five images; deep learning 1D super-resolution output after rotation at 0 degrees; deep learning 1D super-resolution output after rotation at 120 degrees and rotation back to the original frame. Scale bar: 10 μm. Accompanying Fourier transforms are shown in bottom row. **c)** Fixed *C. elegans* L2 larvae (strain DCR6681) expressing GFP-membrane marker imaged in commercial OMX 3D SIM system. A single slice ~2 μm from the bottom surface of the worm is shown, derived from 5 μm stack. Raw data (top) and reconstruction (bottom) are shown. No modulation is evident in raw data, and reconstruction shows obvious artifacts (red arrows). Scale bar: 5 μm. See also **Fig. 6b, c. d)** Imaris 3D renderings of nucleus (H2B-GFP, cyan) and segmented centrosome (EMTB-mCherry, magenta) at indicated time points. The centrosome produces a concave nuclear deformation (red arrows), pulling and rotating the nucleus as it becomes docked at the immune synapse. Scale bar: 5 μm. See also **Fig. 6l. e)** Decorrelation resolution analysis from the images of worm L4 larval (strain DCR8528) expressing membrane targeted GFP primarily in the nervous system. See also **Fig. 6d-j**. Data are derived from 45 measurements (3 animals, 15 planes per animal). Given the ~2.3-fold improvement laterally and ~2.6-fold improvement axially, the triple-view 2D SIM result offers a volume resolution improvement of ~13.8-fold over the raw view C data. **f)** Apparent widths (mean ± standard deviation) of 8 actin fibers from the cell presented in **Fig. 6k-p**, comparing lateral (left) and axial (right) full width at half maximum in different microscope modalities. Given the ~2.2-fold improvement laterally and ~2.4-fold improvement axially, the triple-view 2D SIM result offers a volume resolution improvement of ~11.6-fold over the raw view C data.

**Table S1, Data acquisition and processing parameters for data acquired with the triple-view line confocal platform, related to Figs. 1-6.** The volume size W x H x D represent pixel size in x, y, and z dimensions, C the number of colors and P the number of phases (1 means no phase modulation). The volume acquisition time includes all colors and phases.

| Figure # | Sample/structure (excitation wavelength) | Imaging mode | Acquisition mode | Views A/B volume size W x H x D C x P (z step size) | View C Volume size W x H x D C x P (z step size) | Volume acquisition time | Volume or subvolume # (interval) | Volume size after reconstruction (sample size, $\mu m^3$) | Decon method | Iteration # |
|---|---|---|---|---|---|---|---|---|---|---|
| Fig. 1b | 100 nm yellow-green beads (488 nm) | Triple-view diffraction-limited | Stage scan | 960 x 960 x 50<br>1 x 1<br>(0.5 μm) | 960 x 960 x 50<br>1 x 1<br>(0.5 μm) | 19 s | 1 | 260 x 900 x 200<br>(~25 x 88 x 20) | Addition | 30 |
| Fig. 1c | Fixed U2OS cell, microtubules (488 nm) | Triple-view diffraction-limited | Objective piezo scan | 640 x 480 x 240<br>1 x 1<br>(0. 25 μm) | 960 x 800 x 32<br>1 x 1<br>(0. 25 μm) | 33 s | 1 | 980 x 980 x 82<br>(~ 96 x 96 x 8) | Alternating | 30 |
| Fig. 1e | Expanded U20S cell, mitochondria (488 nm) and DNA (561 nm) | Triple-view diffraction-limited | Stage scan | 1440 x 640 x 168<br>2 x 1<br>(0. 5 μm) | 1920 x 1520 x 104<br>2 x 1<br>(0. 5 μm) | 200 s | 1 | 1440 x 1440 x 530<br>(~ 140 x 140 x 52) | Alternating | 30 |
| Fig. 1i | Fixed *C. elegans* L1 larva, nuclei (488 nm) | Triple-view diffraction-limited | Objective piezo scan | 1920 x 640 x 56<br>1 x 1<br>(0.5 μm) | 640 x 1630 x 96<br>1 x 1<br>(0.5 μm) | 15 s | 2 | 600 x 2300 x 200<br>(~59 x 224 x 20) | Addition | 30 |
| Fig. 2a | Fixed *C. elegans* adult, nuclei (488 nm) | Triple-view diffraction-limited | Objective piezo scan | 1600 x 480 x 72<br>1 x 1<br>(1 μm) | 1120 x 1200 x 72<br>1 x 1<br>(1 μm) | 16 s | 10 | 8930 x 1280 x 574<br>(~ 871 x 125 x 56) | Addition with decay function | 30 |
| Fig. 2h | Fixed *Drosophila* third instar larva, wing disc, myoblast membrane (488 nm) and notum nuclei (561 nm) | Triple-view diffraction-limited | Stage scan | 1280 x 960 x 160<br>2 x 1<br>(1 μm) | 1280 x 1280 x 120<br>2 x 1<br>(0.5 μm) | 59 s | 1 | 1280 x 1280 x 615<br>(~125 x 125 x 60) | Alternating | 15 |
| Fig. 2m | Fixed mouse kidney tissue, nuclei (405 nm), actin (488 nm), tubulin (561 nm), and CD31 (647 nm) | Triple-view diffraction-limited | Stage scan | 960 x 640 x 144<br>4 x 1<br>(1 μm) | 1280 x 1280 x 72<br>4 x 1<br>(0.5 μm) | 145 s | 1 | 1280 x 1280 x 369<br>(~125 x 125 x 36) | Addition | 30 |
| Fig. S2e | Fixed mouse cardiac tissue, | Triple-view diffraction-limited | Stage scan | 960 x 640 x 160<br>2 x 1<br>( 1 μm) | 1280 x1280 x 96<br>2 x 1<br>(0.5 μm) | 32 s | 30 | 5500 x 5500 x 380<br>(~536 x 536 x 37) | Addition | 30 |

| | | | | | | | | | | |
|---|---|---|---|---|---|---|---|---|---|---|
| | nuclei (488 nm) and nonspecific protein (647nm) | | | | | | | | | |
| Fig. S2e | Fixed mouse brain tissue, nuclei (488 nm) and nonspecific marking proteins (647nm) | Triple-view diffraction-limited | Stage scan | 960 x 640 x 168<br>2 x 1<br>( 1 µm) | 1280 x1280 x 88<br>2 x 1<br>(0.5 µm) | 33 s | 30 | 5500 x 5500 x 260<br>(~536 x 536 x 25) | Addition | 30 |
| Fig. 3a | Live HCT-116 cell, mitochondria (488 nm) and lysosome (561 nm) | Triple-view diffraction-limited | Objective piezo scan | 320 x 200 x 24<br>2 x 1<br>( 1 µm) | 640 x 320 x 24<br>2 x 1<br>( 0.5 µm) | 1.2 s | 60<br>(10 s) | 235 x 218 x 164<br>(~23 x 21 x 16) | Addition | 30 |
| Fig. 3c | Live mouse cardiomyocyte, mitochondria (488 nm) | Triple-view diffraction-limited | Objective piezo scan | 1440 x 640 x 40<br>1 x 1<br>(1 µm) | 1440 x 640 x 32<br>1 x 1<br>(1 µm) | 2.0 s | 100<br>(20 s) | 450 x 550 x 329<br>(~44 x 54 x 32) | Addition | 30 |
| Fig. 3e | Live *C. elegans* embryo, nuclei (488 nm) | Triple-view diffraction-limited prediction | Objective piezo scan | N/A | 800 x 480 x 40<br>1 x 1<br>(1 µm) | 0.6 s | 180<br>(3 mins) | 580 x 400 x 369<br>(~57 x 39 x 36) | N/A | N/A |
| Fig. 3h | Live *C. elegans* embryo, AIB neurons (488 nm) | Triple-view diffraction-limited prediction | Objective piezo scan | N/A | 800 x 480 x 56<br>1 x 1<br>(0.7 µm) | 0.75 s | 65<br>(5 mins) | 580 x 370 x 400<br>(~57 x 36 x 39) | N/A | N/A |
| Fig. 4b | Fixed U2OS cell, microtubules (488 nm) | Triple-view 1D SIM | Objective piezo scan | 960 x 640 x 240<br>1 x 5<br>(0. 25 µm) | 1120 x 960 x 40<br>1 x 5<br>(0. 25 µm) | 42 s | 1 | 1300 x 1300 x 200<br>(~ 63 x 63 x 10) | Alternating | 30 |
| Fig. 4c | Fixed B16F10 cell in collagen gel, actin (488 nm) | Triple-view 1D SIM | Objective piezo scan | 640 x 640 x 96<br>1 x 5<br>(1 µm) | 1280 x 960 x 64<br>1 x 5<br>(0.5 µm) | 24 s | 1 | 1045 x 1065 x 438<br>(~ 51 x 52 x 21) | Alternating | 30 |
| Fig. 4e | Fixed HEY-T30 cell in collagen gel, actin (488 nm) and mitochondria (561 nm) | Triple-view 1D SIM | Objective piezo scan | 640 x 640 x 72<br>2 x 5<br>(1 µm) | 960 x 960 x 32<br>2 x 5<br>(0.5 µm) | 29 s | 1 | 1920 x 1920 x 328<br>(~ 94 x 94 x 16) | Alternating | 30 |
| Fig. 4g | Expanded U20S cell, mitochondria (488 nm) and microtubules (561 nm) | Triple-view 1D SIM | Stage scan | 1440 x 640 x 168<br>2 x 5<br>(1 µm) | 1600 x 1500 x 88<br>2 x 5<br>(0.5 µm) | 90 s | 1 | 3200 x 3200 x 900<br>(~ 156 x 156 x 44) | Alternating | 30 |

| | | | | | | | | | | |
|---|---|---|---|---|---|---|---|---|---|---|
| Fig. 4i | Expanded *C. elegans* embryo, neuronal structures (488 nm) | Triple-view 1D SIM | Objective piezo scan | 1280 x 960 x 96<br>1 x 5<br>(1 µm) | 1280 x 1120 x 192<br>1 x 5<br>(0.5 µm) | 76 s | 2 | 2263 x 3462 x 1900<br>(~ 221 x 169 x 93) | Alternating | 30 |
| Fig. 4l | Fixed mouse esophageal tissue, actin (488 nm) and tubulin (561 nm) | Triple-view 1D SIM | Stage scan | 640 x 640 x 88<br>2 x 5<br>(1 µm) | 640 x 640 x 48<br>2 x 5<br>(1 µm) | 39 s | 21 | 3535 x 7502 x 528<br>(~ 172 x 366 x 26) | Alternating | 20 |
| Fig. S4i | Fixed *C. elegans* embryo, tubulin (561 nm) | Triple-view 1D SIM | Objective piezo scan | 640 x 320 x 96<br>1 x 5<br>(0.5 µm) | 640 x 640 x 80<br>1 x 5<br>(0.5 µm) | 59 s | 1 | 740 x 1000 x 821<br>(~ 36 x 49 x 40) | Alternating | 30 |
| Fig. 5c, d | Fixed U2OS cell, microtubules (488 nm) | Single-view 1D SIM | Objective piezo scan | N/A | 1120 x 1120 x 48<br>1 x 5<br>(0.25 µm) | 5 s | 1 | 2240 x 2240 x 246<br>(~ 109 x 109 x 12) | Alternating | 30 |
| | | Single-view 2D SIM prediction | | | 1120 x 1120 x 48<br>1 x 1<br>(0.25 µm) | 1 s | 1 | | Alternating | 20 |
| Fig. 5f | Live Jurkat T cell, EMTB (488 nm) and F-actin (561 nm) | Single-view 2D SIM prediction | Objective piezo scan | N/A | 640 x 480 x 32<br>2 x 1<br>(0.5 µm) | 0.6 s | 150<br>(2 s) | 656 x 640 x 300<br>(~ 32 x 31 x 15) | Alternating | 10 |
| Fig. 5f | Live Jurkat T cell, H2B (488 nm) and EMTB (561 nm) | Single-view 2D SIM prediction | Objective piezo scan | N/A | 640 x 480 x 32<br>2 x 1<br>(0.5 µm) | 0.6 s | 200<br>(5 s) | 492 x 492 x 200<br>(~ 24 x 24 x 10) | Alternating | 10 |
| Fig. 6b, c | Fixed *C. elegans* L2 larva, neuronal structures (488 nm) | Triple-view 1D SIM | Objective piezo scan | 640 x 320 x 32<br>1 x 5<br>(1 µm) | 640 x 320 x 40<br>1 x 5<br>(0.5 µm) | 6.5 s | 8 | 680 x 6356 x 472<br>(~ 33 x 310 x 23) | Alternating | 30 |
| | | Triple-view 2D SIM prediction | | 640 x 320 x 32<br>1 x 1<br>(1 µm) | 640 x 320 x 40<br>1 x 1<br>(0.5 µm) | 1.3 s | | | Alternating | 10 |
| Fig. 6d-f | Fixed *C. elegans* L4 larva, neuronal structures (488 nm) | Triple-view diffraction-limited | Objective piezo scan | 680 X 480 x 40<br>1 x 1<br>(1 µm) | 640 x 640 x 56<br>1 x 1<br>(0.5 µm) | 1.5 s | 1 | 922 x 1168 x 574<br>(~ 45 x 57 x 28) | Alternating | 30 |
| | | Single-view 2D SIM | | N/A | 640 x 640 x 56<br>1 x 1<br>(0.5 µm) | 0.7 s | | | Alternating | 15 |
| | | Triple-view 2D SIM prediction | | 680 X 480 x 40<br>1 x 1<br>(1 µm) | 640 x 640 x 56<br>1 x 1<br>(0.5 µm) | 1.5 s | | | Alternating | 10 |

| | | | | | | | | | | |
|---|---|---|---|---|---|---|---|---|---|---|
| Fig. 6k-p | Live U2OS cell, actin (488 nm) | Triple-view diffraction-limited | Objective piezo scan | 640 X 640 x 56<br>1 x 1<br>( 1 μm) | 960 x 960 x 16<br>1 x 1<br>(0.5 μm) | 3.4 s | 100<br>(10 s) | 1560 x 1720 x 164<br>(~ 76 x 84 x 8) | Alternating | 30 |
| | | Triple-view 2D SIM prediction | | 640 X 640 x 56<br>1 x 1<br>( 1 μm) | 960 x 960 x 16<br>1 x 1<br>(0.5 μm) | 3.4 s | | | Alternating | 15 |

**Movie S1:** Fixed and expanded U2OS cell, immunolabeled to highlight DNA (magenta) and Tomm20 (cyan). Reconstructions compare triple-view deconvolved result vs. raw single view C. See also **Fig. 1e-h**.

**Movie S2:** Whole, fixed adult worm stained with NucSpot Live 488. First part of movie compares reconstructions after imaging with point-scanning confocal, single view line-scanning confocal (View C), and triple-view line-scanning confocal mode with conventional and scattering-compensating deconvolution. Second movie shows triple-view reconstruction with 2136 segmented nuclei overlaid. See also **Fig. 2a-f**.

**Movie S3:** Time lapse imaging in triple-view diffraction-limited mode. Left: HCT-116 cells with MitoTracker Green and Lysotracker Deep Red. Right: cardiomyocyte labeled with MitoTracker Red CMXRos. Comparisons highlight single- vs. triple-view imaging. See also **Fig. 3a-c**.

**Movie S4:** Nuclear imaging (H2B-GFP) in live *C. elegans* embryos. Single-view volume projections (left top) and time-lapse maximum intensity projections (middle) vs. projections from two-step deep learning output (left bottom and right) are compared. hpf: hours post fertilization. See also **Fig. 3e**.

**Movie S5:** Triple-view 1D SIM reconstruction of fixed B16F10 mouse melanoma cells labeled with Alexa Fluor 488 phalloidin, embedded in 1.8% collagen gel. Higher magnification views of deconvolved View C, triple-view diffraction-limited, and triple-view 1D SIM reconstructions are also compared. See also **Fig. 4c, d**.

**Movie S6:** Single-view 2D SIM of Jurkat T cells, also highlighting comparison to raw View C confocal input. Top panels: cell expressing H2B-GFP (cyan) and EMTB-mCherry (magenta); bottom panels: cell expressing EMTB-3XGFP (red) and F-actin-tdTomato (yellow). See also **Fig. 5f-m**.

**Movie S7:** Triple-view 2D SIM of (1) stage L4 larval worm expressing membrane-targeted GFP in nerve ring region (volume projections compare raw View C, triple-view diffraction-limited, 2D SIM, and triple-view 2D SIM) and (2) U2OS cell expressing Lifeact tdTomato (lateral and axial projections compare raw View C vs. triple-view 2D SIM output; inserts show higher magnification views of regions marked with yellow rectangles at different brightness level). See also **Fig. 6d-p**.
